# An unbiased screen identified the Hsp70-BAG3 complex as a regulator of myosin binding protein C3

**DOI:** 10.1101/2022.10.08.511444

**Authors:** Andrea D. Thompson, Marcus J. Wagner, Juliani Rodriguez, Alok Malhotra, Steve Vander Roest, Ulla Lilienthal, Hao Shao, Jaime M. Yob, Benjamin L. Prosser, Adam S. Helms, Jason E. Gestwicki, David Ginsburg, Sharlene M. Day

**Affiliations:** University of Michigan, Department of Internal Medicine, Division of Cardiovascular Medicine, Ann Arbor, MI.; University of Pennsylvania, Department of Internal Medicine, Division of Cardiovascular Medicine and Cardiovascular Institute, Philadelphia, PA.; University of Michigan, Center for Chemical Genomics, Life Sciences Institute, Ann Arbor, MI; University of California San Francisco, Institute for Neurodegenerative Diseases and Department of Pharmaceutical Chemistry, San Francisco, CA.; University of Pennsylvania, Department of Physiology, Philadelphia, PA.; University of Michigan, Departments of Internal Medicine, Human Genetics and Pediatrics, Howard Hughes Medical Institute, and the Life Sciences Institute, Ann Arbor, MI

**Keywords:** Hypertrophic cardiomyopathy, molecular chaperones, MYBPC3, BAG3

## Abstract

**Objective:** We aim to identify regulators of myosin binding protein C3 (MyBP-C) protein homeostasis.

**Background:** Variants in myosin binding protein C3 (*MYBPC3*) account for approximately 50% of familial hypertrophic cardiomyopathy (HCM). Most pathogenic variants in *MYBPC3* are truncating variants that lead to reduced total levels of MyBP-C protein. Elucidation of the pathways that regulate MyBP-C protein homeostasis could uncover new therapeutic strategies that restore normal protein levels.

**Method:** We developed a high-throughput screen to identify compounds that can increase or decrease steady-state levels of MyBP-C in an induced pluripotent stem cell cardiomyocyte (iPSC-CM) model derived from a patient with HCM. To normalize results, we also monitored effects on myosin heavy chain (MYH) and focused on those molecules that selectively modulated MyBP-C levels.

**Results:** Screening a library of 2,426 known biologically active compounds, we identified compounds which either decreased (241/2426, 9.9%) or increased (29/2426, 1.2%) MyBP-C/MYH levels. After a rigorous validation process, including a counter screen for cellular toxicity, two compounds (JG98 and parthenolide) were confirmed as decreasing MyBP-C levels and no compounds were confirmed to increase MyBP-C levels. For further studies, we focused on JG98, which is an allosteric modulator of heat shock protein 70 (Hsp70), inhibiting its interaction with BAG domain co-chaperones. We found that genetic reduction of BAG3 phenocopies treatment with JG98 by reducing MyBP-C protein levels.

**Conclusion:** An unbiased compound screen identified the Hsp70-BAG3 complex as a regulator of MyBP-C stability. Thus, approaches that stimulate this complex’s function may be beneficial in the treatment of HCM.

**Highlights:**

- Hypertrophic cardiomyopathy (HCM) is commonly caused by pathogenic *MYBPC3* variants that reduce total wild-type MyBP-C (the protein encoded by *MYBPC3*).
- It is critical to understand the regulators of MyBP-C protein homeostasis to uncover novel therapeutic strategies.
- We developed and executed a high-throughput chemical screen in iPSC-CMs to identify compounds which alter steady-state levels of MyBP-C protein, revealing two compounds, JG98 and parthenolide, that significantly reduced MyBP-C levels.
- Validation studies suggest that the complex between heat shock protein 70 (Hsp70) and its co-chaperone BAG3 is a dynamic regulator of MyBP-C stability, suggesting that this axis could be a new therapeutic target for HCM.

## Introduction

Hypertrophic cardiomyopathy (HCM) is a genetic cardiomyopathy characterized by left ventricular hypertrophy. Patients can experience a variety of adverse cardiovascular outcomes including left ventricular outflow tract obstruction, heart failure, arrhythmias, and premature death.(1) Cardiac sarcomere genes are the primary genetic basis of HCM, with variants in myosin binding protein C3 (*MYBPC3*), accounting for ∼50% of familial cases.(2)

Significant progress has been made in defining the disease mechanism underlying HCM.(3) For example, greater than 90% of *MYBPC3* variants are known to create premature stop codons (4) that are commonly referred to as “truncating variants.” The mutant mRNA and protein produced by these variants is rapidly cleared from the cell via nonsense mediated decay and the ubiquitin protein system (UPS), respectively.(5–7) This leads to haploinsufficiency of MyBP-C, the protein product of *MYPC3*, with levels reduced by ∼40% in human HCM myectomy tissue.(7–10) Viral delivery of WT *MYBPC3* prevents and reverses disease phenotypes in both animal and cellular models of disease.(11–14) However, in both human inducible pluripotent stem cell-derived cardiomyocytes (iPSC-CMs) and mouse models, a single allelic pathogenic variant is not sufficient to consistently cause disease phenotypes or haploinsufficiency(8, 15), and we previously found that reduced degradation of MyBP-C may compensate for a reduced gene dose.(15) Suggesting compensatory mechanisms could be exploited to restore WT MyBP-C function in patients. However, the factors and pathways that regulate expression, folding, trafficking and/or turnover of MyBP-C remain incompletely defined.

Towards this goal, we developed a high throughput screen using iPSC-CMs derived from a patient with HCM, carrying a *MYBPC3* pathogenic variant. We screened a library of biologically active small molecules to identify those that alter steady-state MyBP-C protein levels. From this screen, we identified the Hsp70-BAG3 complex as a critical regulator of MyBP-C homeostasis, suggesting that this pathway is a potential therapeutic target for HCM due to *MYBPC3* loss of function variants.

## Methods

### Induced pluripotent stem cells (iPSC)

iPSCs used in this study are from two control lines (Ctrl 1,2), three patient-derived lines obtained from patients with HCM who were genotype positive for a pathogenic truncating variant in *MYBPC3* (*MYBPC3* Patient1,2,3) and a gene-edited iPSC homozygous *MYBPC3* knock-out (*MYBPC3* (-/-)) (Supplemental Table 1).(15–17) iPSCs were verified to be free of mycoplasma contamination using MycoAlert Detection Kit (Lonza). Newly generated lines were karyotyped (WiCell Institute, Madison WI) (Supplemental Figure 1).

### Cardiomyocytes (iPSC-CMs)

Details regarding cardiomyocyte production and cell culture are provided in the Supplemental Information Expanded Methods. Unless otherwise noted, iPSC-CMs were pooled from 3 independent differentiations, tested at day 25 of differentiation, and plated at a concentration of 150,000 cells/cm^2^. The iPSC-CMs utilized in our screen demonstrated > 90% purity by immunostaining for α-actinin (Supplemental Figure 2).

### High-throughput Alpha-LISA screen (HTS), 384 well plate format

Alpha-LISA assay development and expanded protocol are provided in Supplemental Information (Expanded Methods, Supplemental Figure 3-5). The screen was performed using *MYBPC3* Patient1 iPSC-CMs, with *MYBPC3* (-/-) iPSC-CMs and 293T cells used as negative controls.

### Alpha-LISA: MyBP-C

Assay mixture included 1:2,000 dilution of mouse monoclonal MyBP-C antibody (C0, Santa Cruz, sc-137180), 1:2,000 dilution of rabbit monoclonal MyBP-C antibody (C5-C7, Samantha Harris, University of Arizona), 1:200 dilution of anti-mouse IgG donor bead (Perkin Elmer, AS104D) and a 1:200 dilution of anti-rabbit IgG acceptor bead (Perkin Elmer, AL104C) dissolved in 1X Alpha-LISA assay buffer (Perkin Elmer, AL000F). Plates read by laser irradiation of donor beads at 680 nm generating chemiluminescent emission from the acceptor bead at 615 nm using EnVision Multimode Plate Reader (Perkin Elmer).

### Alpha-LISA: Myosin Heavy Chain (MYH)

In a separate plate the MYH Alpha-LISA was performed using 1:2,000 dilution of mouse monoclonal BA-G5 antibody to α-MYH (abcam, ab50967), 1:2,000 dilution of rabbit polyclonal β-MYH (abcam, ab228353).

### Compound Libraries

A small molecule library of 2,400 FDA approved or known biologic active compounds (MS2400, Microsource Discovery Systems) was tested. Further, we created a 26-compound custom library of small molecule modulators that target proteins identified to bind or potentially regulate MyBP-C synthesis or degradation (Supplemental Table 2). In total 2,426 compounds were tested.

### Concentration Response Curves

Mosquito X1 (SPT Labtech) was utilized to prepare compound treatment plates and serial dilutions, at the indicated concentrations for concentration response curves performed using compounds from the center for chemical genomics (CCG) library. For powder retesting and secondary assays, fresh powder was obtained (Supplemental Table 3) and dissolved in DMSO to make stock solutions and tested at final DMSO concentration of 0.4%.

### Cell Toxicity Assay

Cellular toxicity was evaluated using CellTiter-Fluor^TM^ cell viability assay (Promega #G6082). *MYBPC3* Patient1 iPSC-CMs were plated and treated with compound in the same manner described above in Black, clear bottom 384 well plates (Breiner Bio-one 781092). After 24 hours of compound treatment, media changes were performed decreasing the volume from 50 μl to 25 μL. Next, 25 μL of substrate-assay buffer was added according to manufactures protocol. Plates were incubated at 37 °C for 30 minutes. A 384-well immuno-grade high-binding black micro plate cover was placed over the clear bottom, and plates were read on fluorometer 380-400 nm excitation with 505 nm emission on pheraSTAR (BMG LABTECH). Buffer only was utilized as positive control (0% viability) and CMs treated with vehicle control was utilized as negative control (100% viability).

### Immunofluorescence

iPSC-CMs, used in the HTS, were plated on coverslips cut from polydimethylsiloxane sheeting (Specialty Manufacturing, Inc) coated with Matrigel at 15,000 cells/cm^2^. The same media and culture conditions were maintained during the compound treatment step. Upon completion of the HTS, immunofluorescence was performed as previously described.(6) Expanded methods including antibodies utilized and image analysis is provided in Supplemental Materials.

### Cycloheximide (CHX) pulse-chase analysis of MyBP-C degradation rates

Neonatal rat ventricular cardiomyocytes (NRVCMs) were prepared as previously described(6). An expanded methods is provided in the supplement. Cells were tested with vehicle control (DMSO). JG98 was tested at its EC_50_ (in NRVCMs this was 1 μM). After 24 hours of compound treatment, CHX chase assay was performed with cells treated with CHX (Sigma Aldrich) 300 μg/ml for the designated durations of 0, 1, 8, 30, 38 hours.

### Adenovirus treatment

Cells were transduced with adenovirus at the indicated multiplicity of infection(MOI) and treatment times. Adenovirus drove expression of scrambled short hairpin RNA (ShRNA)(Vector Biolabs, shADV #207157) or BAG3 ShRNA (Vector Biolabs, shADV #201936). The following adenoviral reagents were provided by Dr. Prosser’s laboratory: scramble shRNA, two vasohibin1 (VASH1) ShRNA, two small vasohibin binding protein SVBP ShRNA, null construct, tubulin tyrosine ligase (TTL).(18) Cells were collected using 1x Laemelii Buffer (Bio-Rad, #1610737) containing protease (Milipore-Sigma, #11836153001) and phosphatase (Roche, #04-906-837-001) inhibitors.

### Western Blot

Immunoblotting was performed as previously described(6). Nitrocellulose membranes were blocked in Intercept Blocking Buffer (Li-COR, #927-70001) for 1 hour. Primary antibodies: rabbit polyclonal BAG3 (Invitrogen, #PA5-78854) 1:1,000; rabbit polyclonal *MYBPC3* (Invitrogen #PA5-79714) 1:500; mouse monoclonal MYBPC3 (Santa Cruz sc-137180) 1:500; monoclonal mouse GAPDH (Sigma, #G8795) 1:1,000; rabbit polyclonal anti-detyrosinated (dTyr) alpha-tubulin antibody (abcam, ab48389); mouse monoclonal anti-alpha tubulin (DM1A) (cell signaling technology #3873) 1:5,000; were resuspended in Intercept Antibody Dilutant Buffer (LI-COR, #927-75001) and stained at 4 °C overnight. Secondary antibodies: IRDye680RD (LI-COR, #926-68070), IRDye800CW (LI-COR, #926-32211), were resuspended in Intercept Antibody Diluent Buffer at room temperature for 1 hour. Nitrocellulose membranes were imaged using an Odyssey Fc (LI-COR). Densitometry analysis was performed using Image Studio analysis software (LI-COR).

### Statistics

Statistical analysis was done using GraphPad Prism software. Kruskal-Wallis nonparametric 1-way ANOVA with Dunn’s post hoc test for multiple comparisons were used for comparisons among control. *P* values < 0.05 were considered statistically significant. Data are reported as mean ± standard error of mean (SEM). Curve fitting was performed as described in the supplemental information.

### Study approval

All animal studies were conducted with the approval of the University of Michigan Institutional Animal Care and Use Committee (IACUC, PRO00009438). All patient samples were collected with the approval of the University of Michigan Institutional Review Board, and all subjects gave written informed consent (HUM00041413, HUM00052165).

## Results

### High Throughput Screening

To identify pathways and targets that regulate MyBP-C stability, we envisioned screening a collection of bioactive small molecules in a cell-based model. To enable this screen, we utilized a commercialized no-wash, homogenous immunoassay technology [Alpha-LISA, Perkin Elmer]. We developed an Alpha-LISA assay targeting MyBP-C using two antibodies linked to donor and acceptor beads (Figure 1A), allowing detection and quantification of MyBP-C within a cellular lysate sample. In choosing cells for this screen, we utilized a well-characterized iPSC cell line derived from a patient with HCM to generate CMs (*MYBPC3* Patient1 (+/c.2373dupG) (15–17) (Figure 1B). Another key design in our screen was the simultaneous measurement of myosin heavy chain (MYH, using antibodies which detect αMYH and βMYH via a parallel Alpha-LISA. This feature allowed us to control for sarcomere content in each well and focus on compounds that selectively modulated MyBP-C via reporting of the MyBP-C/MYH ratio. Expanded results describing the development of the assay and its optimization are provided in Supplemental Information. Briefly, both the MyBP-C and MYH Alpha-LISA exhibited excellent signal-to-noise and a linear relationship between Alpha-LISA signal and total protein concentration when testing *MYBPC3* Patient1 cellular lysates (Figure 1C).

**Figure 1:**
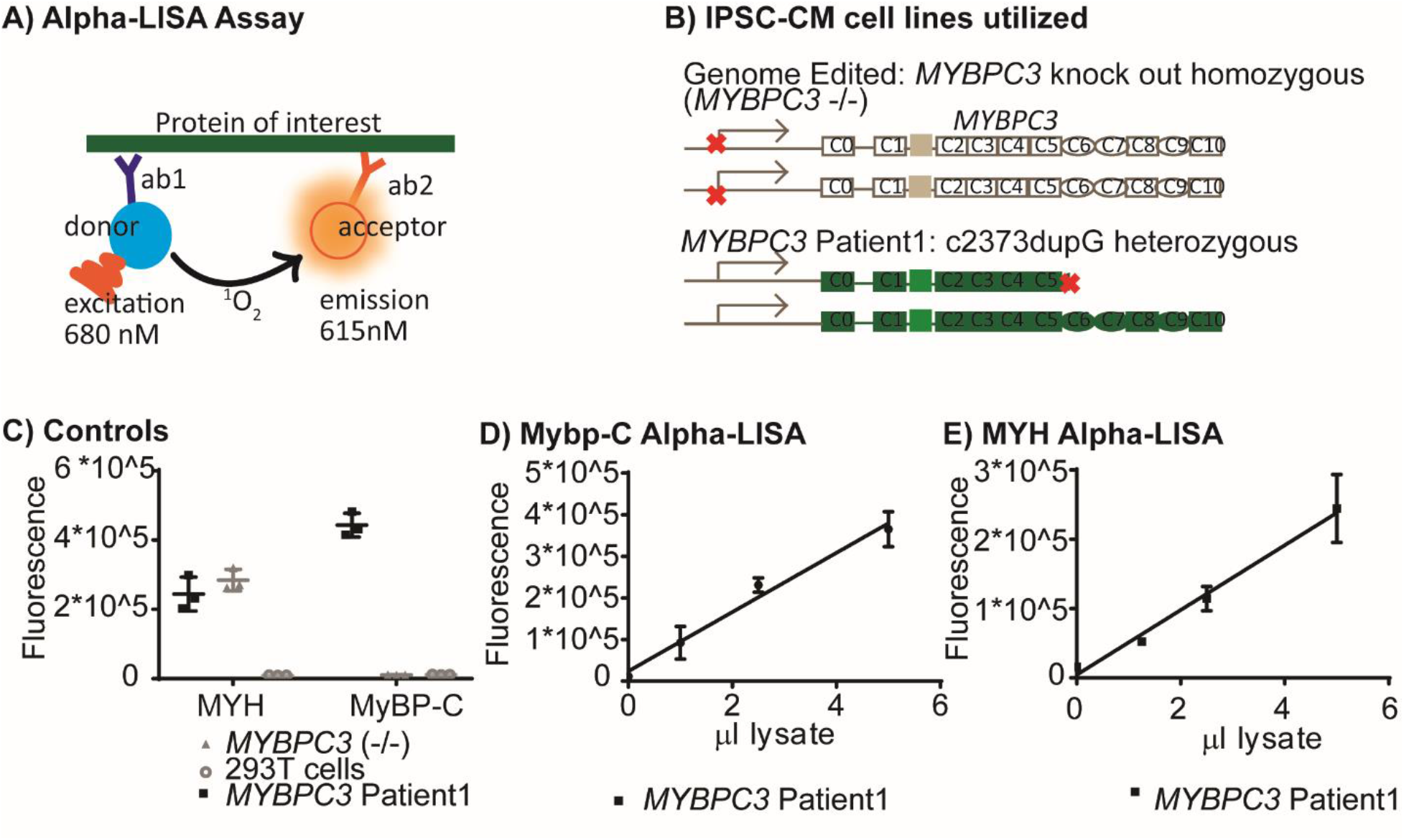
Alpha-LISA MyBP-C/MYH Assay Development. (A) Schematic of Alpha-LISA testing platform. (B) iPSC-CM models used in the screen. (C) MyBP-C and MYH Alpha-LISA raw results under screening assay conditions for positive (*MYBPC3* Patient1) and negative (*MYBPC3* -/-) control iPSC-CMs. 293T cells were also utilized as a negative control. (D &E) Serial two-fold dilution of *MYBPC3* Patient1 iPSC-CM lysate tested in triplicate demonstrates that assay conditions fall within linear range for MyBP-C (D) and MYH (E) Alpha-LISA. Controls C-E were performed for each independent Alpha-LISA experiment.

Using a primary endpoint of MyBP-C/MYH ratio, we screened 2,426 compounds representing FDA-approved molecules, known bioactive compounds and a curated sub-collection of 26 molecules that target known MyBP-C interacting proteins. In these screens, we used in-plate controls to allow determination of the assay robustness, yielding an in-plate Z-factor of 0.66 (0.44-0.83) (Figure 2A, Supplemental Figure 6). Using a criterion of 3 standard deviations from the controls, we identified 241/2426 (9.9%) hit compounds that decreased MyBP-C/MYH levels (Figure 2). We also identified 29/2426(1.2%) hit compounds which increased MyBP-C/MYH levels (Supplemental Table 4).

**Figure 2:**
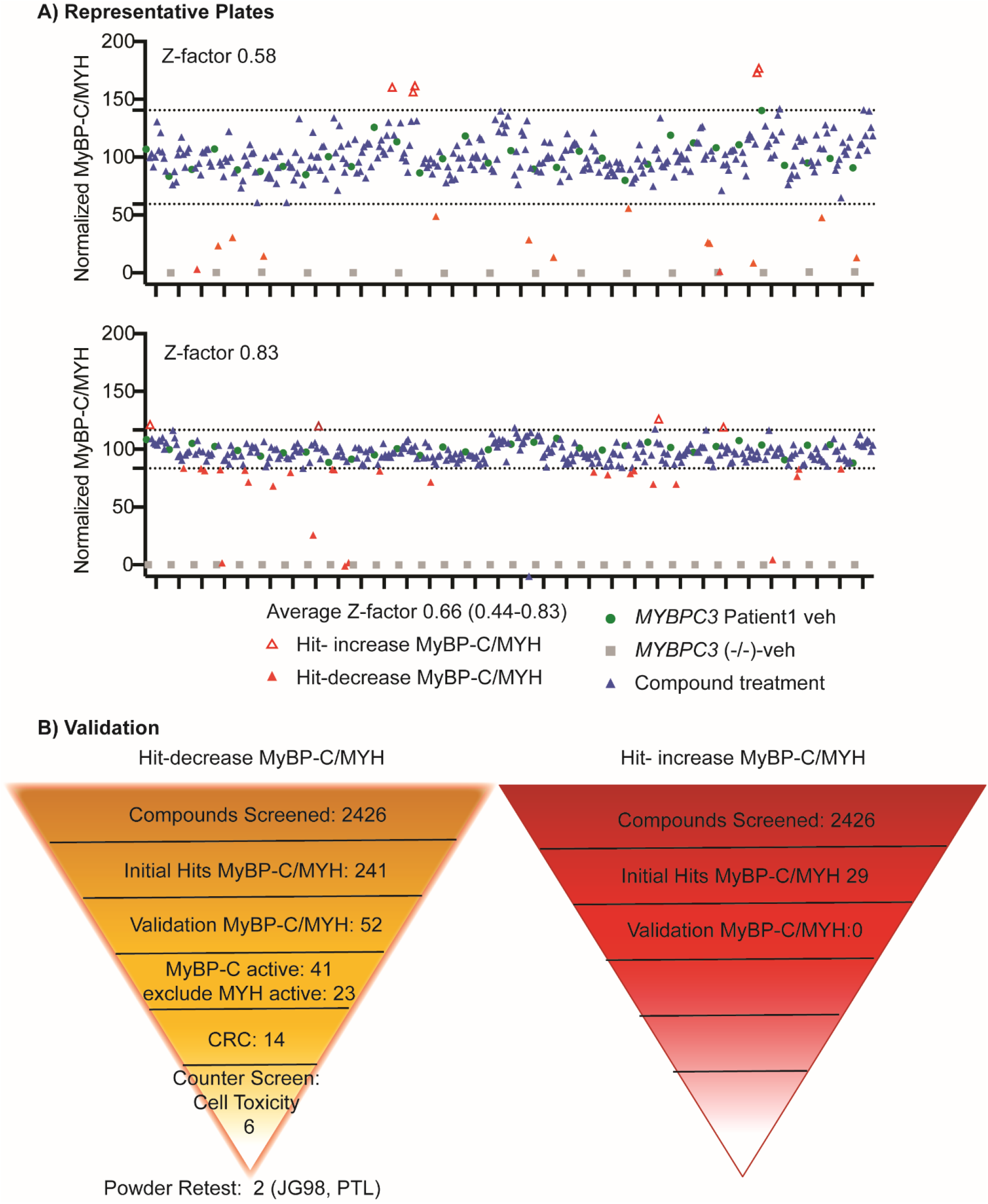
Compound Screen Results. A) Normalized MyBP-C/MYH results from two representative screening plates. 100% is defined by *MYBPC3* Patient1 iPSC-CMs tested with vehicle (DMSO) control (green circles). 0% is defined by *MYBPC3* (-/-) iPSC-CMs tested with vehicle (DMSO) control (grey squares). *MYBPC3* patient1 iPSC-CMs treated with compound are shown in triangles. Hits were defined as compounds which exhibited a MyBP-C/MYH value > 3 standard deviations above (red open triangles) or below (red closed triangles) the control. Most compounds did not significant change MyBP-C/MYH levels (blue triangles). B) Hits subsequently went through a validation process detailed in the flowsheets herein, ultimately identifying two validated hits.

### Validating screening hits

In any HTS campaign, it is important to establish reproducibility of initial hits (Figure 2B, Supplemental Table 5). No compounds that increased MyBP-C/MYH protein levels demonstrated activity after repeat testing. This is likely a result of the low initial hit rate. Further, it is important to use secondary assays to focus on the most robust hits. First, we analyzed MyBP-C and MYH Alpha-LISA separately and selected compounds that reduced MyBP-C levels without significantly altering MYH levels (Figure 2B, Supplemental Table 6-7, Supplemental Figure 7). Finally, we evaluated 14 hits which met the above criteria in a counter screen for cytotoxicity (Supplemental Figure 8, Supplemental Table 8). We then triaged compounds to ensure compounds were non-toxic (EC_50_ in cellular toxicity assay > 60 μM). After these steps, we were left with six non-toxic compounds that decreased MyBP-C/MYH protein levels by decreasing MyBP-C protein levels without a significant effect on MYH protein levels.

To further validate the six active compounds, we re-ordered fresh powders and tested them in the Alpha-LISA (Supplemental Table 3). From these experiments, only JG98 and parthenolide (PTL) retained activity (Figure 3A, Supplemental Figure 9). Specifically, JG98 exhibited an effective concentration for 50% reduction in MyBP-C protein levels (EC_50_) of 12.1 μM (95% CI 8.3-18.0 μM). JG98 also decreased MYH, but at a higher concentration; EC_50_ of 43.8 (29.1-68.4) μM. PTL exhibited an EC_50_ of 43.8 μM (95% CI 29.1-68.4 μM) with no significant activity noted in the MYH assay. This activity was maintained in a control line and two other patient derived lines (Figure 3B, Table 1) and confirmed by western blot analysis (Supplemental Figure 10). JG98 is an allosteric modulator of heat shock protein 70 (Hsp70), which is known to inhibit Hsp70 binding to an important class of co-chaperones, BAG domain proteins.(19, 20) PTL is a natural product that has a variety of reported biologic activities.(21) Thus, these screens identified two compounds that impacted MyBP-C levels.

**Figure 3:**
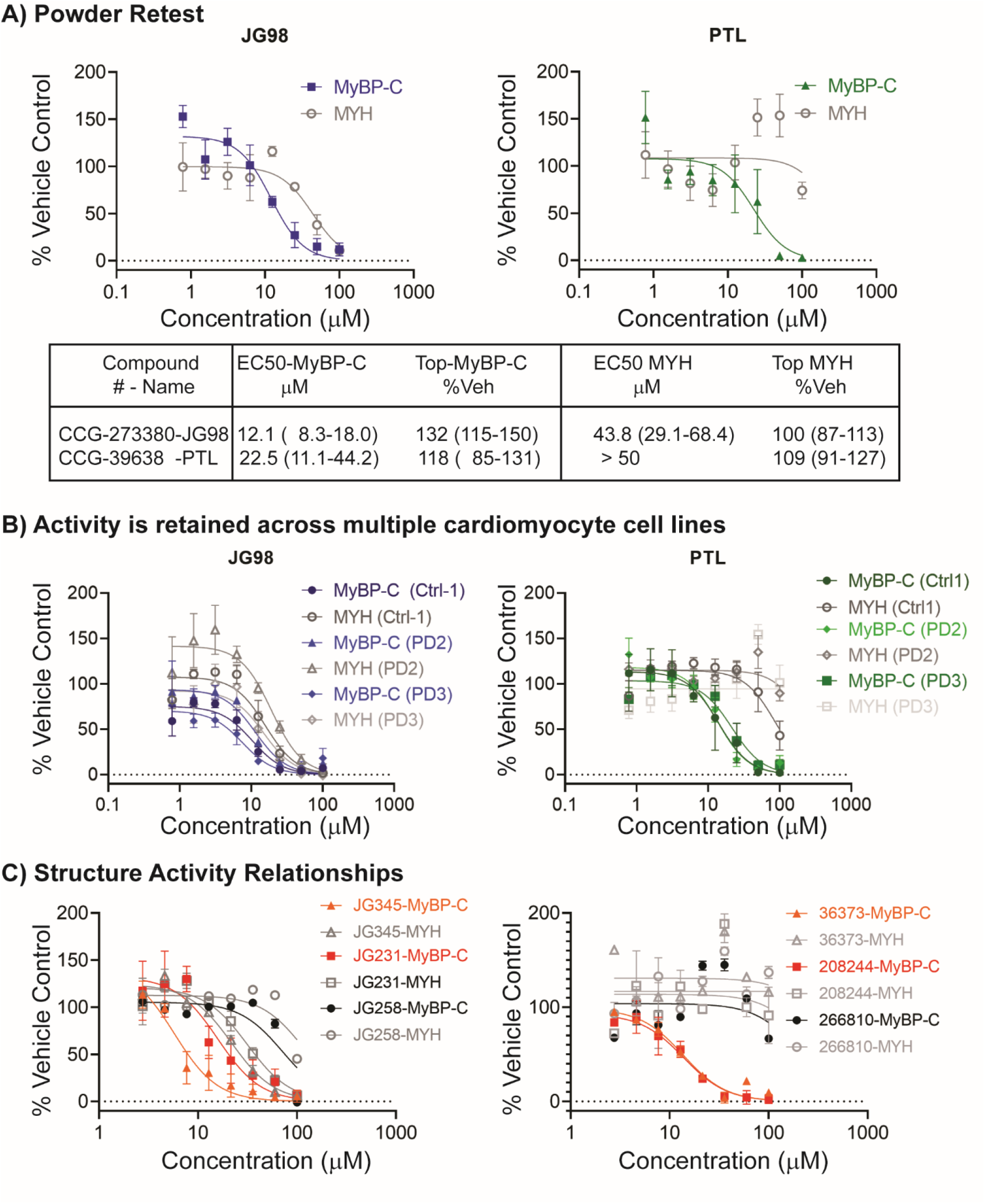
Validated Hits-JG98, PTL. A) Fresh powder was tested, JG98 and PTL retained activity (N = 4). EC_50_ and Top values of curve fitting are reported along with 95% confidence interval (CI). B) Compounds were also tested across a variety of cardiomyocyte cell lines and demonstrated activity throughout (Table1). C) Structure activity relationships (SAR) were explored for JG98 and PTL, the two structurally related compounds with the lowest EC_50_ are shown (orange triangles, red squares). A representative inactive structurally related compound (black circles) is also shown. (Supplemental Table 9).

**Table 1.** Testing PTL and JG98 in multiple cell lines

| Cell<br>lines | MyBP-C |  |  |  | MYH |  |  |  | N |
| --- | --- | --- | --- | --- | --- | --- | --- | --- | --- |
|  | EC<sub>50</sub> | 95% *CI | Top | 95% CI | EC<sub>50</sub> | 95% CI | Top | 95% CI |  |
|  | μM |  | % Veh |  | μM |  | % Veh |  |  |
| JG98 |  |  |  |  |  |  |  |  |  |
| Ctrl-1 | 10.3 | 7.4-14.6 | 75 | 65- 86 | 15.8 | 11.4-22.3 | 107 | 94-121 | 5 |
| †PD-1 | 11.4 | 7.5-17.8 | 94 | 79-109 | 19.2 | 12.2-31.1 | 142 | 120-164 | 7 |
| ‡PD-2 | 7.5 | 4.7-12.6 | 70 | 57- 85 | 14.8 | 9.4-24.4 | 92 | 76-110 | 4 |
| PTL |  |  |  |  |  |  |  |  |  |
| Ctrl-1 | 14.1 | 7.7-26.5 | 113 | 91-138 | >50 | - | 115 | 104-126 | 5 |
| PD-1 | 13.4 | 9.9-18.2 | 118 | 105-131 | >50 | - | 115 | 107-123 | 7 |
| PD-2 | 21.5 | 15.3-31.0 | 104 | 91-116 | >50 | - | 95 | - | 4 |

Legend notes: Concentration response curves were fit using the inhibitor vs. response-
variable slope curve in GraphPad prism. Constraints included Bottom = 0, hill slope = -2.
\*CI = confidence interval (CI). †PD-2 and ‡PD-3 refer to *MYBPC3* Patient2 and Patient3 iPSC-CMs, respectively.

### Structure activity relationships (SAR)

To gain further insight into the validated hits, we tested analogs that are structurally related to JG98 and PTL (Figure 2C, Supplemental Table 9). Our goal was to understand whether the expected SAR would correlate with the observed effects on MyBP-C abundance. In the Alpha-LISA, we found that PTL-related compounds containing the reactive epoxide moiety were active, but those lacking the epoxide were inactive (Supplemental Table 9). One exception was eunicin, but this compound also reduced MYH. These findings are consistent with PTL having a covalent mechanism of action, driven by reactivity of the epoxide moiety.

JG98 is part of a 400+ compound, medicinal chemistry campaign to develop Hsp70-BAG inhibitors (19,22,23). Using this information as a guide, we selected analogs of JG98 to test in the MyBP-C Alpha-LISA. In those screens, we found that the activity of the analogs tracked with their known potency in other systems. For example, JG-258 is a structurally related compound without activity against Hsp70, and it did not reduce MyBP-C protein levels (Figure 2C).(22) Likewise, analogs such as MKT-077, YM-1 and YM-8 are known to have reduced activity against Hsp70, compared to JG98, and they also did not reduce MyBP-C protein levels.(24, 25) (Supplemental Table 9) Conversely, more potent analogs, such as JG-231 and JG-345, reduced MyBP-C protein levels to a greater extent than JG98 (22) (Figure 2C, Supplemental Table 9). Together, these results support the idea that the Hsp70-BAG system is important for MyBP-C stability.

### Mechanism of action for PTL is likely independent of detyrosination of α-tubulin

In cardiomyocytes, one of the biologic activities of PTL that has been well studied is its ability to decrease the amount of detyrosinated α-tubulin.(26) Detyrosinated α-tubulin is a post-translational modification generated by carboxypeptidases (VASH1/2-SVBP) (27, 28) and this process is reversed by tubulin tyrosine ligase (TTL).(29) Thus, it is possible that PTL could reduce MyBP-C through this pathway. To test this possibility, we treated the cells with adenovirus to increase expression of TTL and, separately, knocked down VASH1 with shRNA. We validated that these manipulations reduced detyrosinated α-tubulin, as expected (Supplemental Figure 11). However, there was no change in MyBP-C/MYH. These results suggest that PTL regulates MyBP-C homeostasis via a mechanism distinct from its known effects on detyrosinated α-tubulin. PTL also has many other cellular targets, and further elucidation of the mechanism by which PTL leads to alterations in MyBP-C homeostasis could be pursued in future studies.

### JG98 identifies Hsp70-BAG3 as a key regulator of MyBP-C protein homeostasis

We subsequently focused on understanding the mechanism by which JG98 reduces MyBP-C protein levels. Based on the mechanism of JG98 action on a chaperone complex, we hypothesized that compound treatment would enhance degradation of MyBP-C protein and shorten its cellular half-life. Indeed, a cycloheximide pulse-chase assay performed in JG98 treated NRVCMs showed more rapid degradation of MyBP-C with a corresponding shortening in MyBP-C half-life (Figure 4A). This change occurred without obvious effects on sarcomere integrity (Figure 4C). Next, we focused on the role of BAG3, one of the BAG family of co-chaperones that is highly expressed in cardiomyocytes. This protein localizes to the sarcomere (30, 31), where it is thought to play a role in folding and assembly of sarcomere proteins. Thus, we hypothesized that JG98 induces MyBP-C degradation by disrupting Hsp70-BAG3 interaction (Figure 4D). We performed shRNA knock down of BAG3 and, consistent with our hypothesis, found that this treatment reduces MyBP-C protein levels (Figure 5). Taken together, this result provides strong evidence that the Hsp70-BAG3 axis is critical for regulating MyBP-C levels.

**Figure 4:**
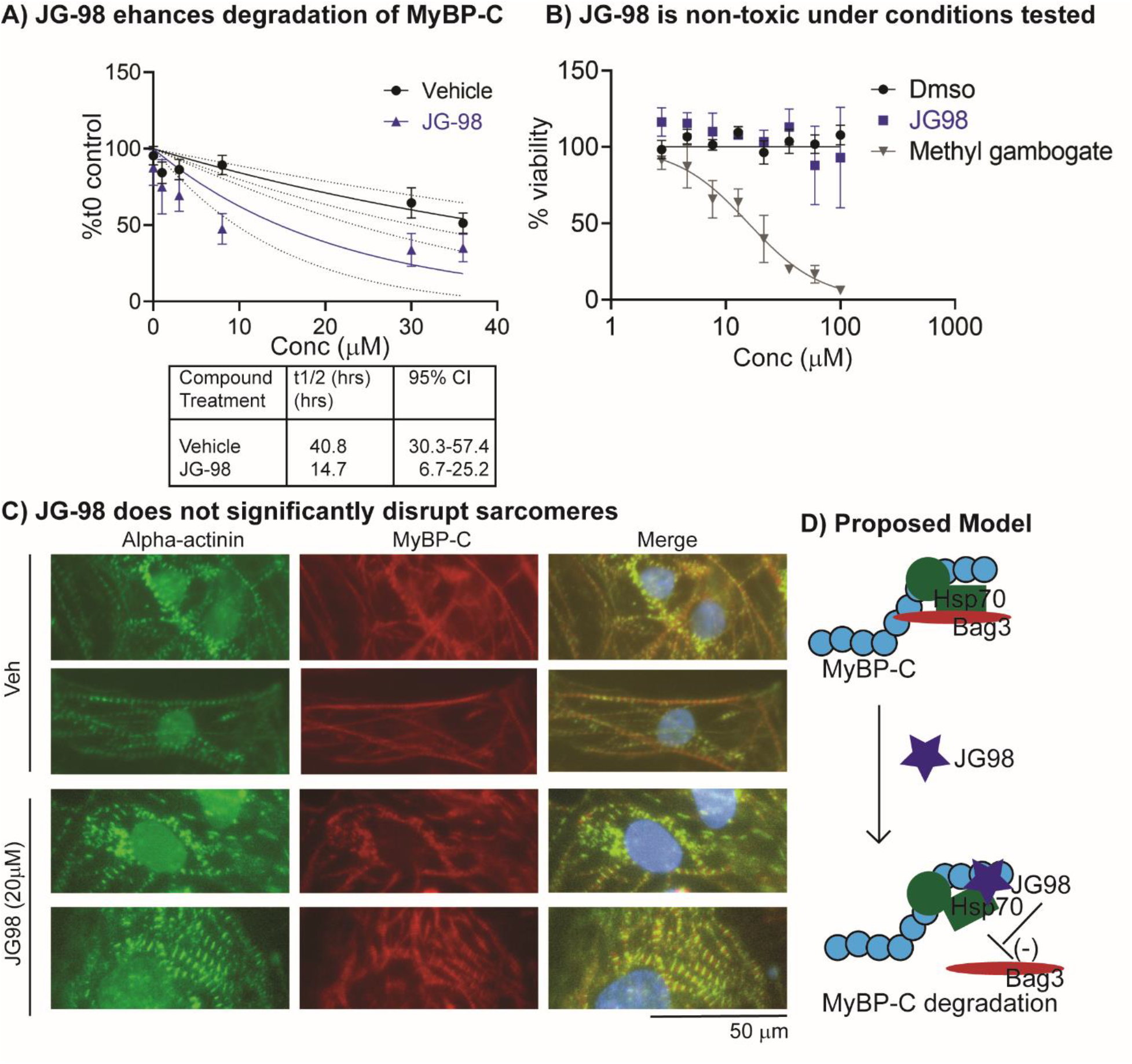
JG98 enhances MyBP-C degradation without inducing toxicity or disrupting sarcomere structure. A) Cycloheximide (CHX) chase assay demonstrates that treatment with JG98 results in more rapid clearance of MyBP-C from neonatal rat ventricular cardiomyocytes (NRVCMs) compared to vehicle control (veh, dmso). JG98 was tested at 1 μM for 24 hrs. [EC_50_ of JG98 in NRVCMs is 1.2 μM (95% CI-0.4-3.00)] NRVCMs then underwent treatment with CHX at time intervals indicated. Cells were tested in quadruplicate on independent biologic replicates (N=8). B) JG98 is non-toxic (EC_50_ >60 μM) within *MYBPC3* Patient1 iPSC-CMs. Positive control methyl gambogate with cellular toxicity, is shown (n = 4). (Also shown in Supplemental Figure 8 and Table 8). C) Sarcomere structure and MyBP-C localization is largely preserved after JG98 (20 μM for 24 hrs) treatment in *MYBP3* Patient1 iPSC-CMs. Representative images are shown. α-actinin (green) and MyBP-C (red). D) Proposed model: JG98 induces MyBP-C degradation by inhibiting Hsp70 and Bag3 binding.

**Figure 5:**
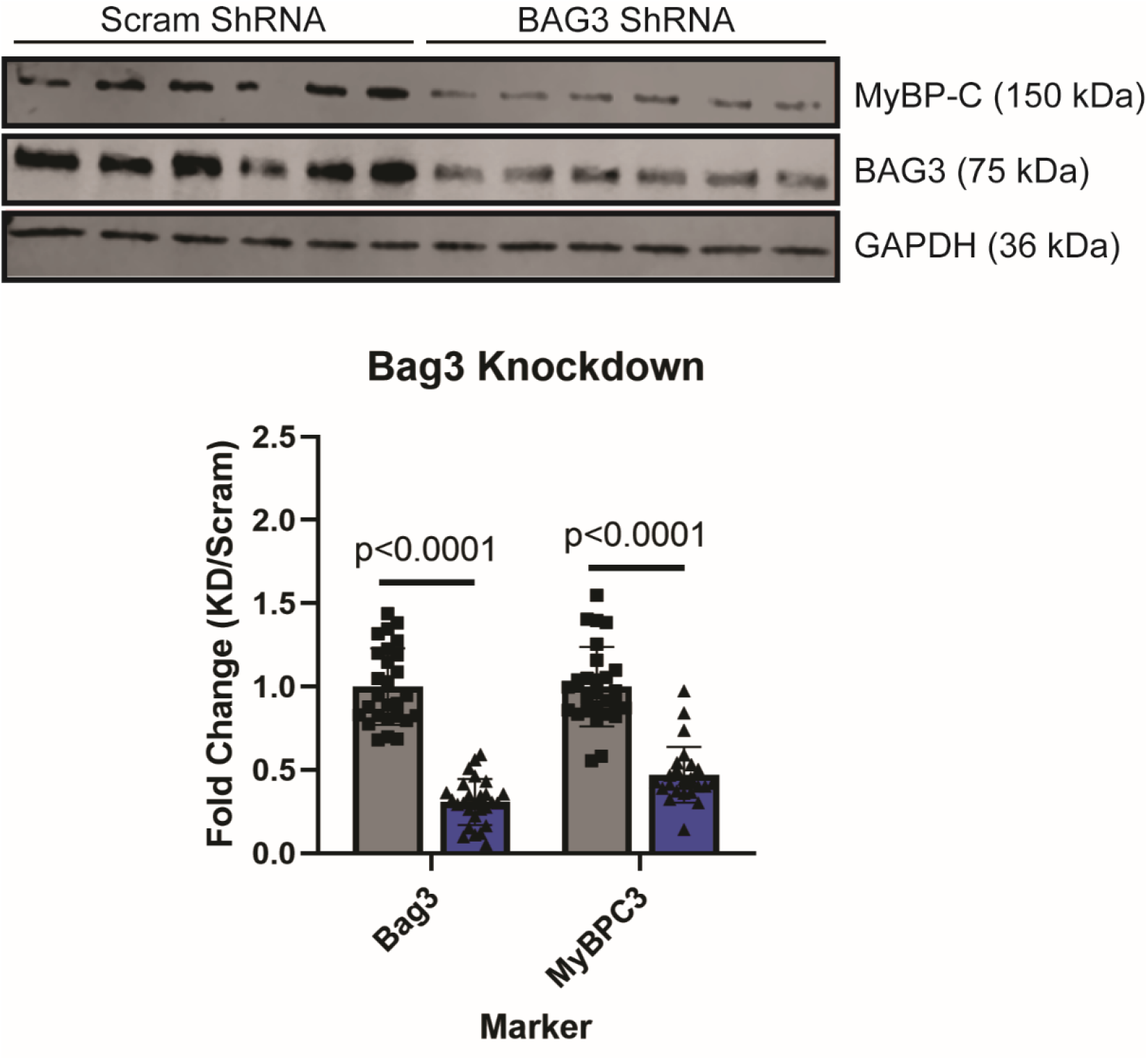
Reduction in BAG3 decreases MyBP-C protein levels. A representative western blot is shown. Ctrl-2 iPSC-CMs were cells treated with an adenovirus at a MOI 5 carrying scrambled shRNA (grey bars, black squares) or BAG3 shRNA (blue bars, black triangles) for 18 hours. Cellular lysates were analyzed 4 days after shRNA treatment (n =27, 5 biologic replicates). Quantification of western blots demonstrates significant reduction of BAG3 and MyBP-C, defining 1.0 as the average band intensity for the shRNA scramble control. Glyceraldehydre-3-phosphate dehydrogenase (GAPDH) was also used as a loading control.

## Discussion

Pathogenic truncating variants of *MYBPC3* lead to hypertrophic cardiomyopathy by causing haploinsufficiency.(7, 9) However, haploinsufficiency is not consistently observed in published mouse or iPSC-CM models with heterozygous truncating *MYBPC3* variants.(8, 15) Our group has shown previously that this compensatory response in iPSC-CMs occurs by slowing the degradation of the remaining wild-type protein.(15) Thus, identifying pathways which regulate MyBP-C protein levels is important to understanding the pathophysiology of HCM and developing therapeutic strategies.(3) Herein, we developed and executed a high throughput, small molecule screen, identified two small molecules that reduce MyBP-C protein levels, and pinpointed the Hsp70-BAG3 axis as a critical intervenable target to control MyBP-C abundance.

One exciting product of this work is the development of a robust high throughput screen for measuring MyBP-C levels. Although in this study we focused on a small library of bioactive compounds, this platform is amenable to screening larger, diverse collections. More immediately, we propose that the two compounds identified by our initial screen (JG98 and PTL), may serve as useful experimental probes for studying MyBP-C protein homeostasis. The activity of PTL and JG98 was consistent across multiple iPSC-CM cell lines, demonstrating it is not dependent on the genetic background of the cardiomyocyte cell line or influenced by batch-specific variables of cardiomyocyte differentiation. Understanding the molecular mechanism by which these compounds reduce MyBP-C may uncover pathways which contribute to the development of MyBP-C haploinsufficiency in HCM patients.

PTL is a natural product containing an epoxide moiety which enables it to make covalent modifications to cysteine amino acids.(21) In testing PTL analogs, we demonstrated that the ability to reduce MyBP-C protein levels is dependent on this epoxide. However, the target protein which leads to alterations in MyBP-C protein levels remains unknown. PTL is known to have multiple biologic cellular effects, including inhibition of HDAC1, modulation of NF-kB, and reduction in the amount of detyrosinated α-tubulin, among others (21, 26). Our results suggest that the activity of PTL on detyrosinated α-tubulin is not relevant to its activity on MyBP-C, suggesting that other targets must be involved. Given the pleiotropic cellular targets of PTL, identification of this target is expected be challenging and is outside the scope of this work.

In contrast, our initial results with JG98 and its analogs strongly suggest that the Hsp70-BAG complex is important for MyBP-C homeostasis. JG98 is known to bind an allosteric site on Hsp70s, trapping a conformation that has weak affinity for BAG domain proteins (25). Importantly, a structurally related control that lacks activity against HSP70, JG-258, did not affect MyBPC protein levels, supporting the idea that the target of JG98 is Hsp70 in cardiomyocytes. This conclusion is also consistent with prior studies from our group showing that heat shock protein 70 (Hsc70), an Hsp70-family member, targets MyBP-C for ubiquitin-dependent proteasomal degradation.(6) Further, expression of Hsp70 was noted to be downregulated in iPSC-CM models in which normal MYBP-C levels were maintained despite the presence of *MYBPC3* loss of function (truncating) variants.(15) Thus, this study adds further compelling evidence that converges on Hsp70 chaperones as critical and central regulators of MyBP-C homeostasis. Interestingly, our pilot screen also contained other Hsp70 modulators, which were not active. These other compounds, such as 115-7c, are known to bind other sites on hsp70 and to have distinct effects on other (non-BAG) co-chaperones.(32) Thus, these observations suggest that the Hsp70-BAG complex may be the most important for MyBP-C stability.

BAG domain proteins bind Hsp70 via their BAG domain and regulate nucleotide exchange. In addition, the six mammalian BAG proteins also contain additional functional domains which influence their cellular localization and function.(33) Among these, BAG3 was of particular interest to us because it is known to be localized to the sarcomere, and rare variants in BAG3 are associated with dilated cardiomyopathies.(34, 35) Further, GWAS studies have identified common *BAG3* variants as risk alleles for clinical phenotypes of DCM and protective alleles for HCM.(36–38) We hypothesized that Hsp70-BAG3 might stabilize MyBP-C, such that its disruption would reduce MyBP-C levels (Model Figure 4D). Consistent with this hypothesis, shRNA knock down of BAG3 also reduces MyBP-C. Thus, although multiple BAG proteins might be involved, disruption of BAG3 is sufficient to produce the phenotype. Yang and colleagues recently explored knock down of BAG3 in iPS-CMs, showing that reduced BAG3 leads to sarcomere damage after 10 days, along with a mild reduction in MyBP-C levels by immunofluorescence but not by western blot.(39) In our study, we evaluated a more acute disruption of BAG3 by examining cells after 24 hours of JG98 treatment or 96 hours of shRNA knock down. We chose this approach because chaperone effects on stability are typically rapid, and compensation by other factors (e.g., other BAG proteins) can occur at longer times. In our studies, we observed robust reductions of MyBP-C levels, including by cycloheximide pulse-chase studies. This activity appears to occur through direct binding of the Hsp70-BAG3 complex to MyBP-C, because proteomics experiments have previously confirmed that BAG-3, Hsp70, and MyBP-C interact.(30, 40) However, the mechanism by which this complex regulates MyBP-C stability is not yet clear. Some work has shown that the Hsp70-BAG3-MyBP-C protein complex facilitates clearance via chaperone assisted selective autophagy (CASA).(30), but for other client proteins (*i.e.,* not MyBP-C), BAG3 routinely binds to Hsp70 and, depending on the substrate and chaperone complex, can both protect against or assist UPS-mediated degradation of client proteins.(41, 42) Our results from cardiomyocyte experiments support a model in which the complex plays a primarily stabilizing role. Thus, we anticipate that strategies that enhance the function of Hsp70-BAG3 would be protective in HCM associated with truncating variants in *MYBPC3*, and potentially more broadly across a spectrum of cardiomyopathies, as this complex is likely to play a stabilizing role for other cardiomyocyte proteins.

### Study Limitations

Our results should be interpreted in the context of their limitations. First, we were unable to validate any compounds that increase MyBP-C/MYH ratios. Such compounds would clearly be interesting, as they could be the starting point for therapeutics with the desired effect. However, the absence of haploinsufficiency in iPSC-CM cellular models is an important limitation(15) (43), as overexpression of MyBP-C may not be achievable due to inherent maintenance of sarcomere stoichiometry (ref). Developing iPSC-CM models that exhibit haploinsufficiency, will be important to successfully identifying small molecules which increase this ratio back to wild-type levels.

Secondly, we acknowledge the limitations of compounds identified in our study. Both PTL and JG98 have poor potency, which limits their utility as chemical probes. Likewise, PTL has many known targets, and JG98 modulates the function of Hsp70, a chaperone that acts on multiple cellular client proteins. Future work will be needed to understand the broader effects of these compounds on the proteome. Moreover, while JG98 did not have significant toxicity or effects on sarcomere disruption under the conditions in this study, cardiotoxicity has been observed for JG98 under different conditions.(44)

## Conclusion

By screening known, biologically active compounds in an iPSC-CM model, we uncovered that the Hsp70-BAG3 complex is an intervenable target to directly modulate MyBP-C homeostasis independent of myosin. These results suggest that therapeutically targeting stabilization of the Hsp70-BAG3-MyBP-C interaction or increasing BAG3 protein levels may be beneficial in HCM caused by pathogenic *MYBPC3* variants.

## Clinical Perspectives

### Competency in medical knowledge

Hypertrophic cardiomyopathy is commonly caused by pathogenic *MYBPC3* variants that reduce total levels of wild-type MyBP-C protein. However, the pathways which regulate *MYBPC3* protein levels are poorly understood. By screening known, biologically active compounds in an induced pluripotent stem cell cardiomyocyte model this study identified the Hsp70-BAG3 complex as a key regulatory of MyBP-C protein levels.

### Translational Outlook

These results suggest that therapeutically targeting stabilization of the Hsp70-BAG3-MyBP-C interaction or increasing BAG3 protein levels may be beneficial in HCM caused by pathogenic *MYBPC3* variants.

## Acknowledgements

We would like to acknowledge Samantha Harris (University of Arizona) for providing the C5-C7 MyBP-C antibody. We acknowledge Christine L. Mummery (Leiden University) for providing the patient derived induced pluripotent stem cell line *MYBPC3* patient1. We acknowledge the center for chemical genomics staff, in particular director Andrew Alt, PhD and Aaron Robida, PhD and Nick Santoro PhD for guidance and technical support.

## Supplemental Information

### Expanded Methods

#### Human myocardial tissue

Interventricular septal tissue was obtained from gift of life donor hearts at time of explant (n = 3). Donor hearts were perfused with cardioplegia solution before removal. All tissues were snap-frozen in liquid nitrogen immediately after excision. These samples were utilized during assay development.

#### Cardiomyocyte (iPSC-CMs) production

iPSCs were cultured in STEM flex media (Gibco #A3349401) and passaged weekly using EDTA (Invitrogen #15575020) diluted to 0.5 mM in PBS (Gibco#10010-023). Stem cells were grown to confluency in 6-well plates for cardiac differentiation using wnt modulation as previously described.(1) Stem cell derived cardiomyocytes (CMs) underwent metabolic selection using glucose-deprived, lactate-containing media for five days as previously described.(2–4) Following lactate purification CMs were frozen in Cyrostor (Sigma) in liquid nitrogen.

#### Cardiomyocyte (iPSC-CMs) cell culture

Cell culture plates coated with Matrigel (Corning #354277) diluted 1:500 in DMEM/F12 (Gibco, #11330-032) at room temperature for 1 hour. Matrigel/DMEM is removed and cardiomyocytes are plated using replating media [RPMI+, 2% FBS, 0.2 μl/mL of 10 mM thiazovivin (Cayman Chemical, #14245) in sterile dmso (dimethyl sulfoxide, Sigma #D2650)] and maintained in RPMI+ P/S [500 ml-RPMI 1640 [+] L-glutamine [+] phenol red, Gibco #11875-093, 5ml of Penn/Strep 10,000 U/ml Gibco #15140122, 10 mL of B27 supplement, serum-free (Life technologies #17504-001]. Control-2 CMs (Ncyte cardiomyocytes, Ncardia) were cultured according to the manufactures recommendation-plated at 300,000 cells/well in 24 well plates coated with fibronectin (Sigma, #F1141),1:100 in PBS + CaCl and MgCl (Gibco, #1404-117), using iPSC-CM complete medium containing: 10% FBS (Gibco, #16000-044), 1% Penicillin/Streptomycin (Gibco, #15140-122), 2% B27 (Gibco, #17504-044) and supplemented with Y-276321 (Cayman Chemical, #10005583).

#### Alpha-LISA assay development

Alpha-LISA is a bead-based assay which uses luminescent oxygen-channeling chemistry. An analyte is bound in proximity by two antibodies linked to donor and acceptor beads, respectively. Upon photoexcitation at 680nm, the donor bead releases a singlet oxygen species (^1^O_2_) which can induce an acceptor bead to emit at 615 nm only if it is within 200nm of the donor bead (Figure 1A). This assay allows for detection of your protein of interest within a complex mixture of cellular lysate. To develop an Alpha-LISA assay targeting MyBP-C, a variety of MyBP-C antibodies were selected and screened using different antibody and Alpha-LISA bead combinations (Supplemental Table 10). These were tested using myocardial tissue lysate as a positive control and cellular lysate from *MYBPC3* (-/-) iPSC-CMs, which do not express MyBP-C, as a negative control. The antibody and bead combination that provided the strongest signal/noise ratio within human tissue in this initial screen were selected for use in the assay (Supplemental Figure 3). The MyBP-C Alpha-LISA assay demonstrated Z-factor of 0.89 and excellent plate uniformity when tested using human myocardial tissue lysate (Supplemental Figure 5).

A parallel assay was developed to control for sarcomere content, using antibodies targeting myosin heavy chain (MYH) (Supplemental Figure 4, Supplemental Table 10). 293T cellular lysate was utilized as a negative control, as 293T cells do not express MYH. The MyBP-C and MYH Alpha-LISA assays were optimized using the same approach, in which human myocardial tissue lysate was utilized to adapt this assay into a 384-plate format and the effect of antibody concentration and incubation time were evaluated (Supplemental Figure 3,4). The resulting final assay conditions are detailed below.

#### Alpha-LISA MyBP-C

Cellular lysate (2.5 μl) was transferred per well to MyBP-C assay plate (384-alphaplate, Perkin Elmer) using Biomek FX (Beckman Coulter). 2.5 μl of lysis buffer (1X PerkinElmer, AL003C) and 20 μl of assay mixture was added using Multidrop Combi (Thermo Scientific). This assay mixture included 1:2,000 dilution of mouse monoclonal MyBP-C antibody (C0, Santa Cruz, sc-137180), 1:2,000 dilution of rabbit monoclonal MyBP-C antibody (C5-C7, Samantha Harris, University of Arizona), 1:200 dilution of anti-mouse IgG donor bead (Perkin Elmer, AS104D) and a 1:200 dilution of anti-rabbit IgG acceptor bead (Perkin Elmer, AL104C) dissolved in 1X Alpha-LISA assay buffer (Perkin Elmer, AL000F). Plates were covered with foil seal protectant and incubated for 48 hours and then read by laser irradiation of donor beads at 680 nm generating chemiluminescent emission from the acceptor bead at 615 nm using EnVision Multimode Plate Reader (Perkin Elmer). Positive controls HMCL1 treated with vehicle (100% vehicle control) and negative controls *MYBPC3* (-/-) (0%) were tested within each plate.

#### Alpha-LISA Myosin Heavy Chain (MYH)

In a separate plate the MYH Alpha-LISA was performed. The following changes to Alpha-LISA assay protocol were made to adapt it for detection of MYH. Cellular lysate (1.25 μL) was transferred to MYH assay plate (384-Alpha-plate, Perkin Elmer) using Biomek FX (Beckman Coulter). This was chosen to avoid signal saturation typically was not observed until cellular lysate greater than 5 μL but on a few occasions was observed in the MYH assay at 5 μL. The final volume of lysis buffer remained 5 μl with 20 μl of assay mixture was added using Multidrop Combi (ThermoScientific). In the MYH assay a 1:2,000 dilution of mouse monoclonal αMYH antibody (abcam, ab50967), 1:2,000 dilution of rabbit polyclonal βMYH antibody (abcam, ab228353) were utilized. Positive controls HMCL1 treated with vehicle (100% vehicle control) and negative controls 293T cells (0%) were tested within each plate.

#### Compound selection for custom library

Compounds selected for the custom library were based on literature review for proteins and pathways identified to bind or affect MyBP-C. Compounds which target these proteins or pathways were tested. We began by reviewing literature regarding transcription, translation, and degradation of MYBP-C. MYBP-C is known to be degraded via the ubiquitin proteasome system in Hsp70 dependent manner.(5) Thus Hsp70 compounds and proteasome inhibitors were selected for our custom library (Supplemental table 2). Further, MyBP-C was shown to interact with several other molecular chaperones including Hsp90, small heat shock proteins, and VCP/p97(5) so modulators of these additional potential targets were also selected (Supplemental Table 1). Finally, differential expression of molecular chaperones and the potential protective role for heat shock response has been noted in a variety of models of cardiac injury, thus inducers of heat shock response were selected for evaluation (Supplemental Table 2). MyBP-C pathogenic variants which result in a premature stop codon, result in nonsense mediated decay of pathogenic variant mRNA product.(6) An inhibitor of nonsense mediated decay gentamicin(6) was already present within the MS2400 library. Both hypertrophic and adrenergic agonists have been demonstrated to influence expression of sarcomere genes including MyBP-C.(7–9) Adrenergic agonists phenylephrine, norepinephrine, isoproterenol(7,10,11) were present within the MS2400 library. Hypertrophic agonists TGF-beta and ET-1 were included in the custom library.(Supplemental Table 2) It is well established that MyBP-C is phosphorylated(12), thus inhibitors of both protein kinases and phosphatases known to regulate MyBP-C phosphorylation were selected (Supplemental Table 1). Finally, we tested the effect of compounds used to treat hypertrophic cardiomyopathy to evaluate their effect on MyBP-C homeostasis. These were largely present within the MS2400 library such as metoprolol tartrate, verapamil, valsartan. The Mavacamten was also tested in our custom library (Supplemental Table 2).

#### High-throughput Alpha-LISA screen, 384-well plate format

Prior to plating cardiomyocytes, 384-well flat polystyrene clear bottom plates (Greiner, Sigma Aldrich # M6936) were coated with Matrigel as described above (50 μl/well). Matrigel/DMEM media was removed just before plating using BioTek 405 Select microplate washer (BioTek). *MYBPC3* Patient1 and *MYBPC3* (-/-) CMs were thawed at room temperature and diluted to desired concentration (15,000 cells/well in our 384 well plates= 150,000 cells/cm^2^) in replating media. CMs were plated using a Multidrop Combi (Thermo Scientific), applying 50 μl/well. Cells were maintained in RPMI + P/S for 5 days in 384 well plates prior to compound treatment, media changes occurred 24 hours after plating and every 48 hours thereafter. Throughout the experiment BioTek 405 Select microplate washer (BioTek) and Multidrop Combin (Thermo Scientific) were utilized for buffer removal and delivery. Cells were then treated with screening compounds. At the time of compound treatment CMs were at 17-22 days of differentiation. Each well was treated with 20 μM of a library compound, unless otherwise noted for custom compounds (Supplemental Table 2), using Biomex FX (Beckman Coulter) with a 200nl head attachment. Cells were incubated with compound for 24 hours and the final concentration of dmso was 0.4 % in all wells. Columns 1 and 2 and 23 and 24 were treated with dmso alone and served as vehicle control samples [two columns-MYBPC3 Patient1 iPSC-CMs and two columns -*MYBPC3* (-/-) iPSC-CMs]. All compound treated wells contained *MYBPC3* Patient1 iPSC-CMs. 293T cells were grown in triplicate (3 wells) and treated under the same conditions. After compound treatment, cells were washed with PBS containing phosphatase inhibitor (PhosSTOP Roche #4906845001) and protease inhibitor (complete mini, EDTA free, Roche #1183670001) for 1 minute and lysed using 50 μl of Alpha-LISA lysis buffer (Perkin Elmer, AL003C) for 1 hour at room temperature, 200 rpm on plate shaker. After lysis samples were frozen at −80°C prior to performing Alpha-LISA assay as described below.

#### Selection of MyBP-C/MYH as primary endpoint

It is not uncommon to observe systematic experimental error associated with the culturing of cells within 384-well plates while performing phenotypic cellular screens. To evaluate for this, we next re-evaluated plate uniformity and z-factor within the 384-well format while culturing *MYBPC3* Patient1 iPSC-CMs under planned screening conditions. Indeed, we observed increased variability and a systematic error in which wells near the plate edge exhibited higher values than wells near the center of the plate (Supplemental Figure 5). We hypothesized part of this variability may be due to variations in sarcomere content within each well. To test this hypothesis, we were able to take advantage of the MYH alpha-LISA assay. We tested if utilizing MyBP-C/MYH as our primary endpoint, to account for MYH protein levels (as a surrogate for sarcomere content) would reduce variability. Indeed, we observed reduced variability [z-factor of 0.46 compared to 0.39 for MyBP-C alone]. Further, the observed systemic error, or plate effect, was also improved (Supplemental Figure 5). Thus, MyBP-C/MYH was selected as the primary endpoint for our screen. This optimization process resulted in the high-throughput Alpha-LISA screen within a 384-well plate format as detailed within our expanded methods section above.

#### Immunofluorescence (IF)

iPSC-CMs used in the HTS plated on coverslips cut from polydimethylsiloxane sheeting (Specialty Manufacturing, Inc) coated with Matrigel at 15,000 cells/cm^2^. The same media and culture conditions are maintained during the compound treatment step. Upon completion of the HTS, immunofluorescence was performed as previously described.(5) Antibodies were as follows; α-actinin mouse monoclonal 1:500 (Sigma Aldrich, A7811), Myosin light chain-2v (MLC-2v) rabbit polyclonal 1:200 (ProteinTech, 10906-1-APP), MyBP-C rabbit polyclonal 1:1,000 (custom Samantha Harris, University of Arizonia), goat anti-mouse IgG Alexa Fluor 594 1:500 (ThermoFisher Scientific, A11005); and goat anti-rabbit IgG Alexa Fluor 488 1:500 (ThermoFisher Scientific, A11008). Images of 3 random fields per coverslip were acquired using a Nikon Eclipse Ti-E inverted fluorescence microscope. Quantification of % cells positive for α-actinin and MLC-2v was obtained by manually counting using overlay of 4’,6-diamidino-2-phenylindole (DAPI) and protein of interest. Sarcomere structural integrity was evaluated qualitatively. In cells treated with JG-98 IF readings obtained on FITC channel were exposed for 30 minutes prior to imaging.

#### Cycloheximide (CHX) pulse-chase analysis of MyBP-C degradation rates

Neonatal rat ventricular cardiomyocytes (NRVCMs) were prepared as previously described(5). NVCMs were plated at a density of 4.0×10^4^ cells/well in 96-well white clear bottom plates (Costar #3610) in replating media and maintained in RPMI + P/S. Cells were tested with vehicle control (dmso) Jg-98 tested at EC_50_ (in NRVCMs this was 1 μM). After 24 hours of compound treatment. CHX (Sigma Aldrich) was freshly dissolved in dmso at 100 mg/mL), CHX is added to RPMI + P/S with dmso or JG-98 at CHX 300 μg/ml) for the designated lengths of 0, 1, 8, 30, 38 hours. These time intervals were performed in quadruplicate wells. Buffer control (no cells, which had previously been determined to give signal equivalent to *MYBPC3* (-/-) iPSC-CMs were utilized as negative controls within each 96-well plate. Upon completion of this experiment, samples were again washed with PBS three times and lysed using 50 ml/well Alpha-LISA lysis buffer (Perkin Elmer, AL003C) and incubating at room temperature for 1 hour. Samples were stored at −80°C sealed with an adhesive cover (microAmp optical adhesive film, Applied BioSystems) until MyBP-C Alpha-LISA could be performed as described above.

#### Expanded Statistics

Concentration response curve analysis was performed by normalizing data to positive and negative controls as defined above and fitting data to non-linear inhibitor vs response, variable slope curve.

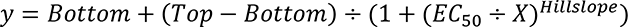

Where; Top and Bottom are the plateaus in the units of the Y-axis, EC50 is the concentration of the agonist that gives a response half way between Bottom and Top, Hillslope describes the steepness of the family of curves. If constraints were utilized this is indicated.

For all Alpha-LISA assays, the following vehicle controls (using the same dmso concentration as that utilized in compound treatment) were performed on the following cell lines; the iPSC-CMs of interest [for example, in the case of the high-throughput screen this was *MYBPC3* Patient1], *MYBPC3* (-/-) iPSC-CMs, and 293T. Two-fold serial dilutions starting at 5μl of cellular lysate from iPSC-CMs of interest were tested in parallel.

Results were normalized as follows.

MyBP-C & MyBP-C/MYH Alpha LISA:

100% iPSC-CMs of interest,

0% *MYBPC3* (-/-) iPSC-CMs

MYH Alpha LISA:

100% iPSC-CMs of interest,

0% 293T cells

For all screening plates, in plate Z-factor was calculated as using vehicle control treated cells as follows;

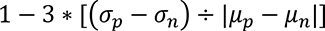

Where;

*σ_p_*= standard deviation of positive control (*MYBPC3* Patient1 CMs

*σ_n_*= standard deviation of negative control (*MYBPC3* (-/-) iPSC-CMs)

*μ_p_* = mean of positive control

*μ_n_* = mean of negative control

CHX chase data analysis was performed as previously described(5).

### Expanded Results

#### Validation of initial hits

For compounds that decreased MyBP-C/MYH we retested initial hits on biological independent cells, tested in triplicate. 52/241 initial hits which decreased MyBP-C/MYH, exhibited a value 3 standard deviations below the positive control in all 3 replicates (Supplemental Table 4). None of initial 70 hits which increased MyBP-C/MYH were able to increase MyBP-C/MYH in all 3 replicates by > 2 stdev (Figure 2). To identify pathways which regulate MyBP-C homeostasis, our ideal hit would decrease MyBP-C protein levels while demonstrating minimal effects on MYH. Thus, during the validation step we also evaluated the MyBP-C and MYH assays separately, using MYH assay as a counter screen. 41/52 compounds decreased MyBP-C > 3 stdev below vehicle control in all 3 replicates. Of these, 23/41 did not increase or decrease MYH by > 3 stdev of control. These 23 compounds were considered our validated hits (Supplemental Table 4), (Figure 2). Next, we re-tested our validated hits on biologically independent *MYBPC3* Patient1 iPSC-CMs at varying concentrations to evaluate concentration response curves (CRCs). Fourteen of these 23 compounds were identified as active (Supplemental Figure 7, Supplemental Table 5), warranting further investigation. Finally, we wanted to ensure we were not selecting compounds with significant cellular toxicity. Thus, CRCs of the 14 active compounds were tested using CellTiter-Fluor^TM^ cell viability assay. 6/14 compounds were identified as non-toxic (EC_50_ > 60 μM) (Supplemental Figure 7, Supplemental Table 7).

---

**Supplemental Figure 1:**
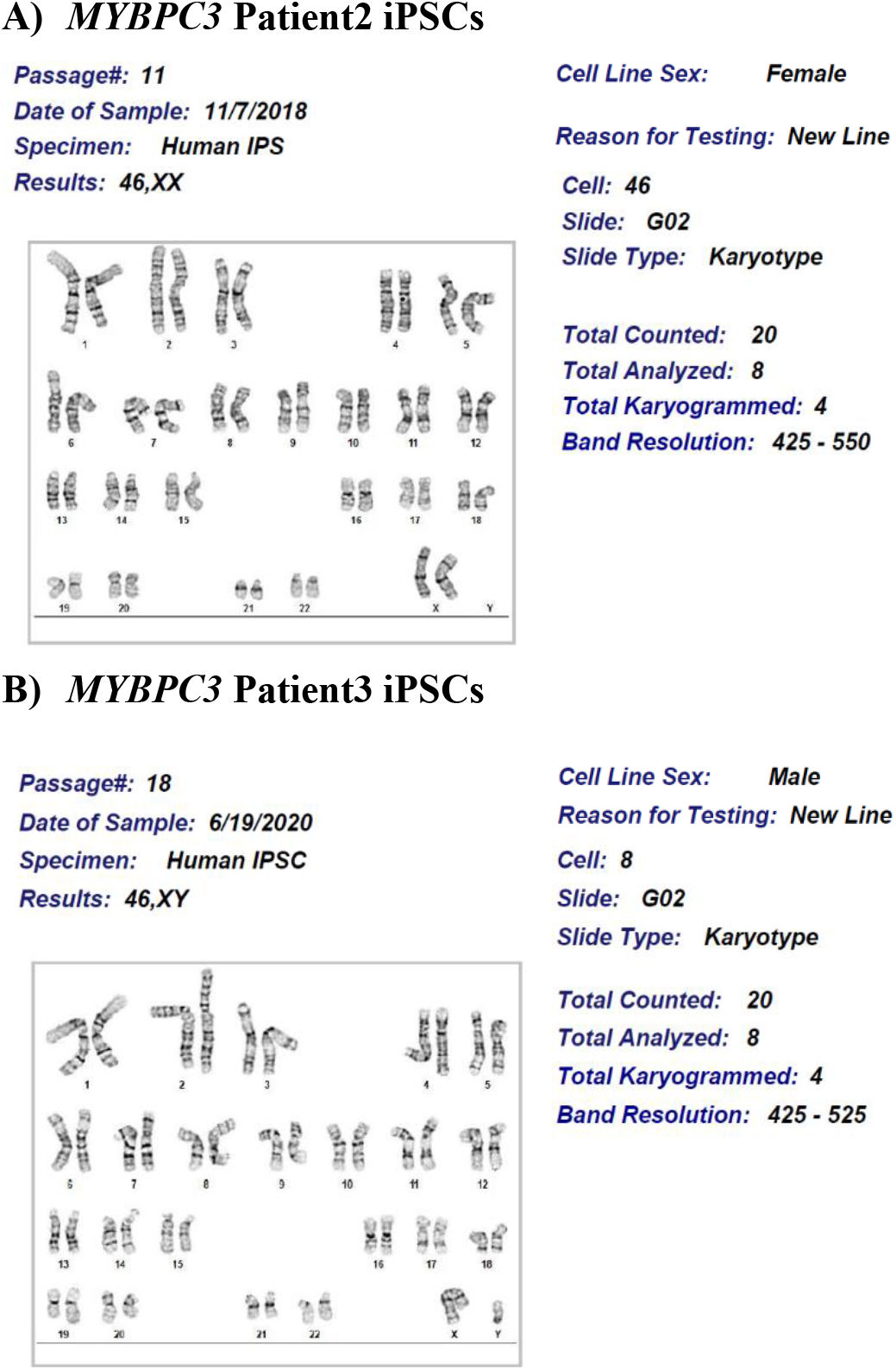
Karyotypes performed by WiCell Institute on newly generated induced pluripotent stem cell (iPSC) lines. A) *MYBPC3* Patient2 and B) *MYBPC3* Patient3 (Supplemental Table 1).

**Supplemental Figure 2:**
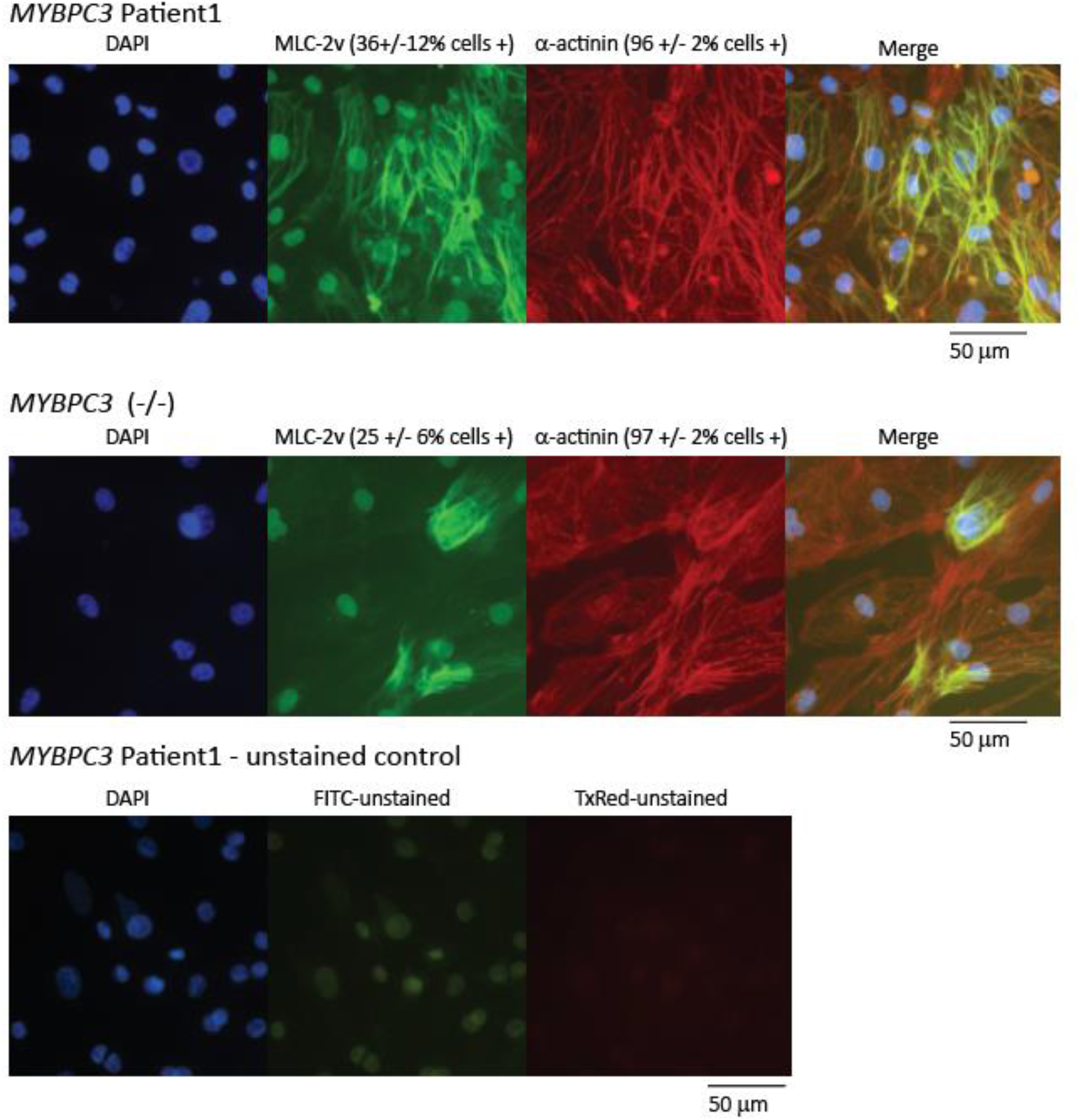
Immunofluorescence on cardiomyocytes (iPSC-CMs) utilized in high-throughput screen. The screen was performed using *MYBPC3* Patient1 iPSC-CMs. Further, *MYBPC3* (-/-) iPSC-CMs were utilized as a negative control (Supplemental Table 1). Both cell lines demonstrated > 90% purity (> 90% of cells positive for cardiac marker α-actinin, red). Cells were also stained for MLC-2v (green). To evaluate for auto-fluorescence, *MYBPC3* Patient1 cells were imaged after undergoing protocol lacking exposure to primary antibodies.

**Supplemental Figure 3:**
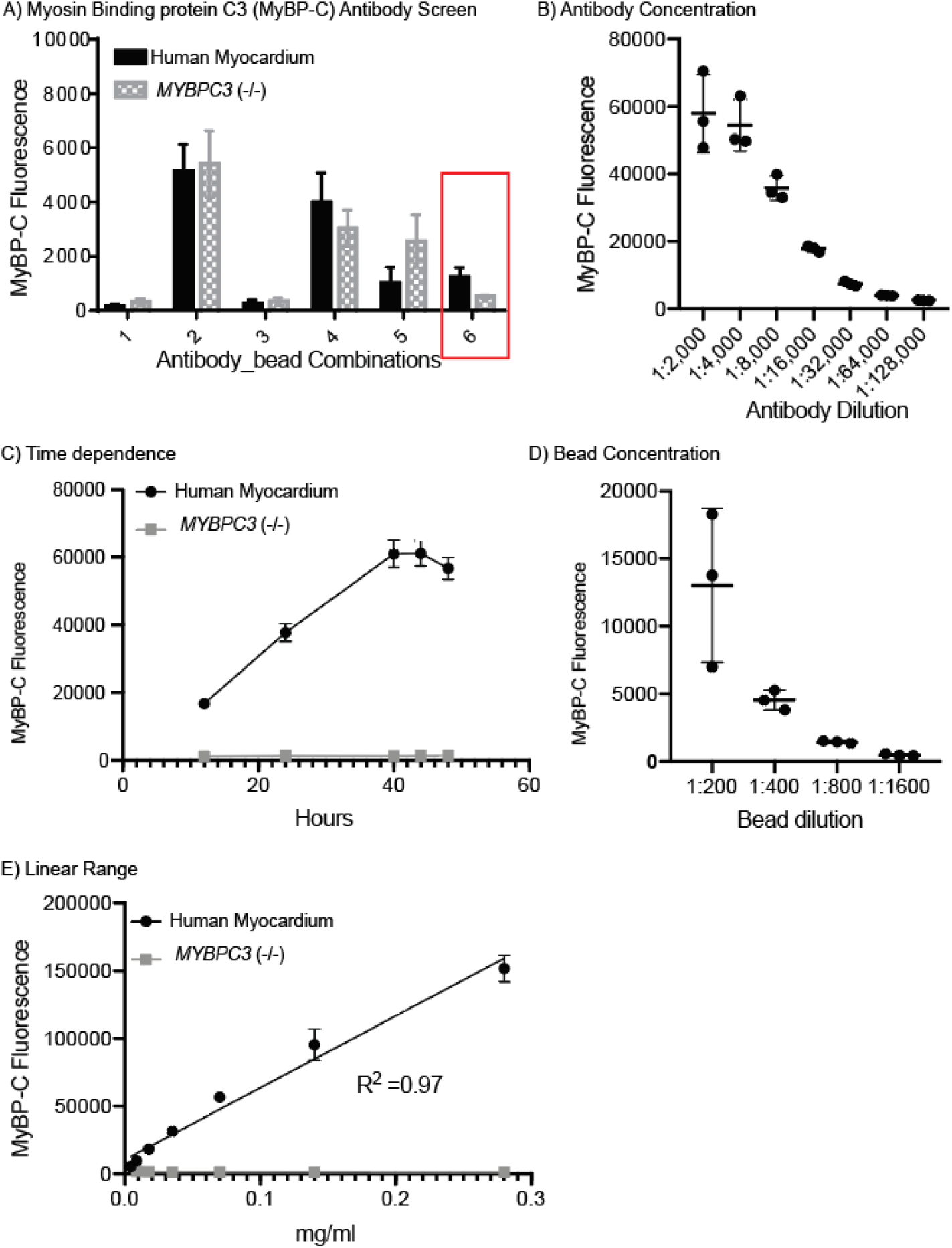
Optimization of MYBP-C Alpha-LISA Assay Conditions. A) Screening various antibody combinations for detection of MyBP-C within human myectomy tissue and *MYBPC3* (-/-) iPSC-CMS (negative control known to lack expression of MyBP-C). Both samples were tested at a total protein concentration 0.3 mg/ml. Antibody combination with best signal to noise ratio was selected for further optimization (red box). B) The relationship of antibody titer and MyBP-C fluorescence was evaluated. C) Time dependence of MyBP-C Fluorescence was tested. D) The relationship of bead concentration and MyBP-C fluorescence was evaluated. E) The relationship of total protein concentration in MyBP-C Fluorescence was evaluated in human myocardial tissue. *MYBPC3* (-/-) iPSC-CMs was evaluated as a negative control. Average and standard error of mean for samples tested in triplicate are shown in each graph.

**Supplemental Figure 4:**
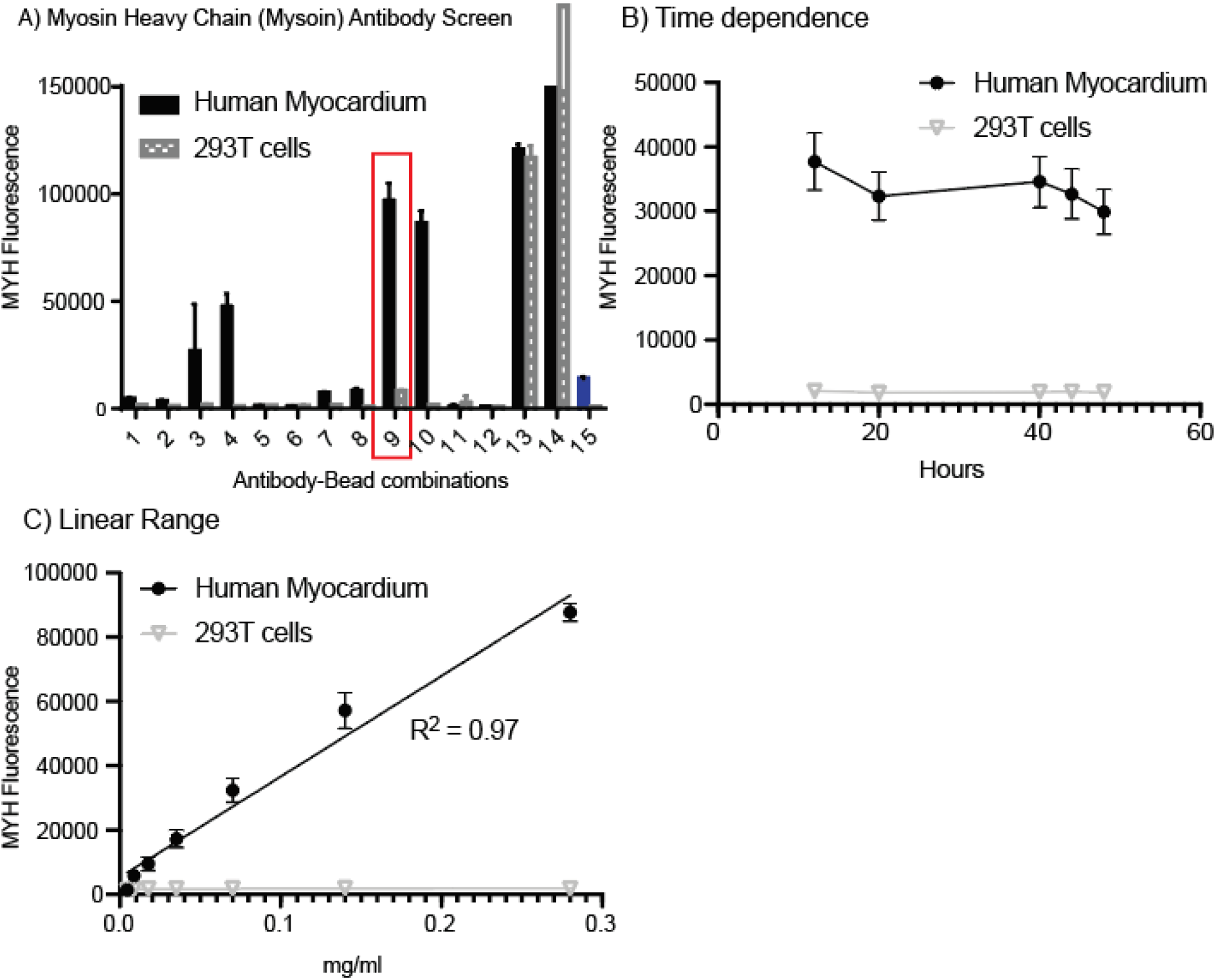
Optimization of Myosin Heavy Chain AlphaLISA Assay Conditions. A) Screening various antibody combinations for detection of myosin heavy chain (MYH), including antibodies for both αMYH and βMYH7, were tested within human myectomy tissue and 293T cells (negative control known to lack aMYH and bMYH expression). Both samples were tested at a total protein concentration 0.3 mg/ml. Antibody combination 9 was selected (red box) and includes antibodies targeting both MYH6 and MYH7. As a positive control optimized MyBP-C antibody and bead combination was tested in 15 (blue bar graph). B) Time dependence of MYH Fluorescence was tested. C) The relationship of total protein concentration in MYH Fluorescence was evaluated in human myocardial tissue and *MYBPC3* (-/-) iPSC-CMs was evaluated. Average and standard error of mean for samples tested in triplicate are shown in each graph.

**Supplemental Figure 5:**
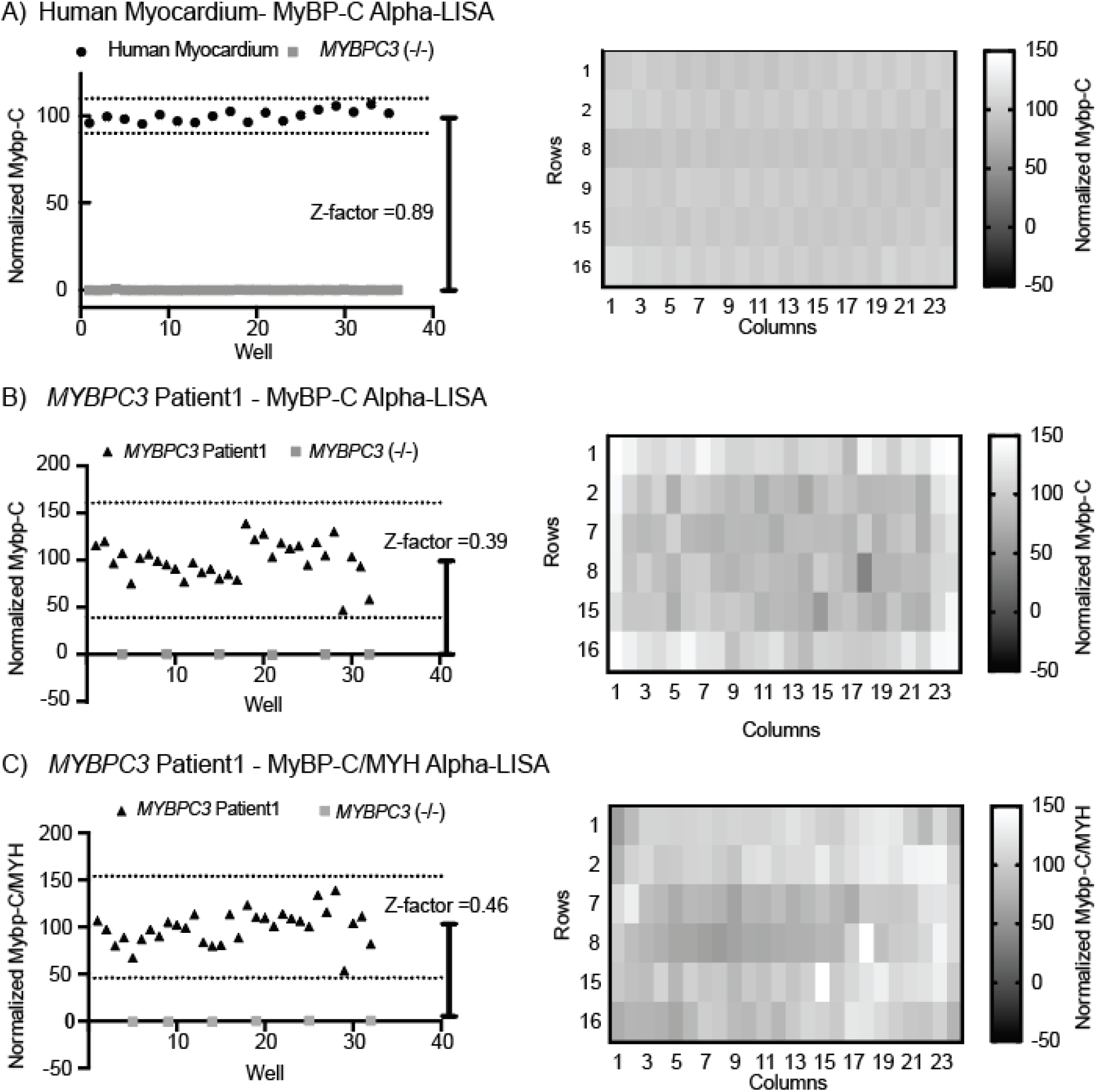
Plate Uniformity. For (A) Human myocardium-MyBP-C Alpha-LISA and (B) *MYBPC3* Patient1 iPSC-CMs grown in 384 well plates an MyBP-C Alpha-LISA plate uniformity test was performed as shown on the right. Further, a Z-factor analysis was performed using *MYBPC3* (-/-) as a negative control (0%) as shown on the left. Culturing *MYBPC3* Patient1 iPSC-CMs does result in increased variability within the MyBP-C Alpha-LISA assay with reduced signal observed within outer edges of the plate. This is improved when MyBP-C Alpha-LISA assay and MYH Alpha-LISA assay are performed in parallel on cellular lysate from *MYBPC3* patient 1 iPSC-CMs grown in 384 well plates using MyBP-C/MYH as the primary assay endpoint (C).

**Supplemental Figure 6:**
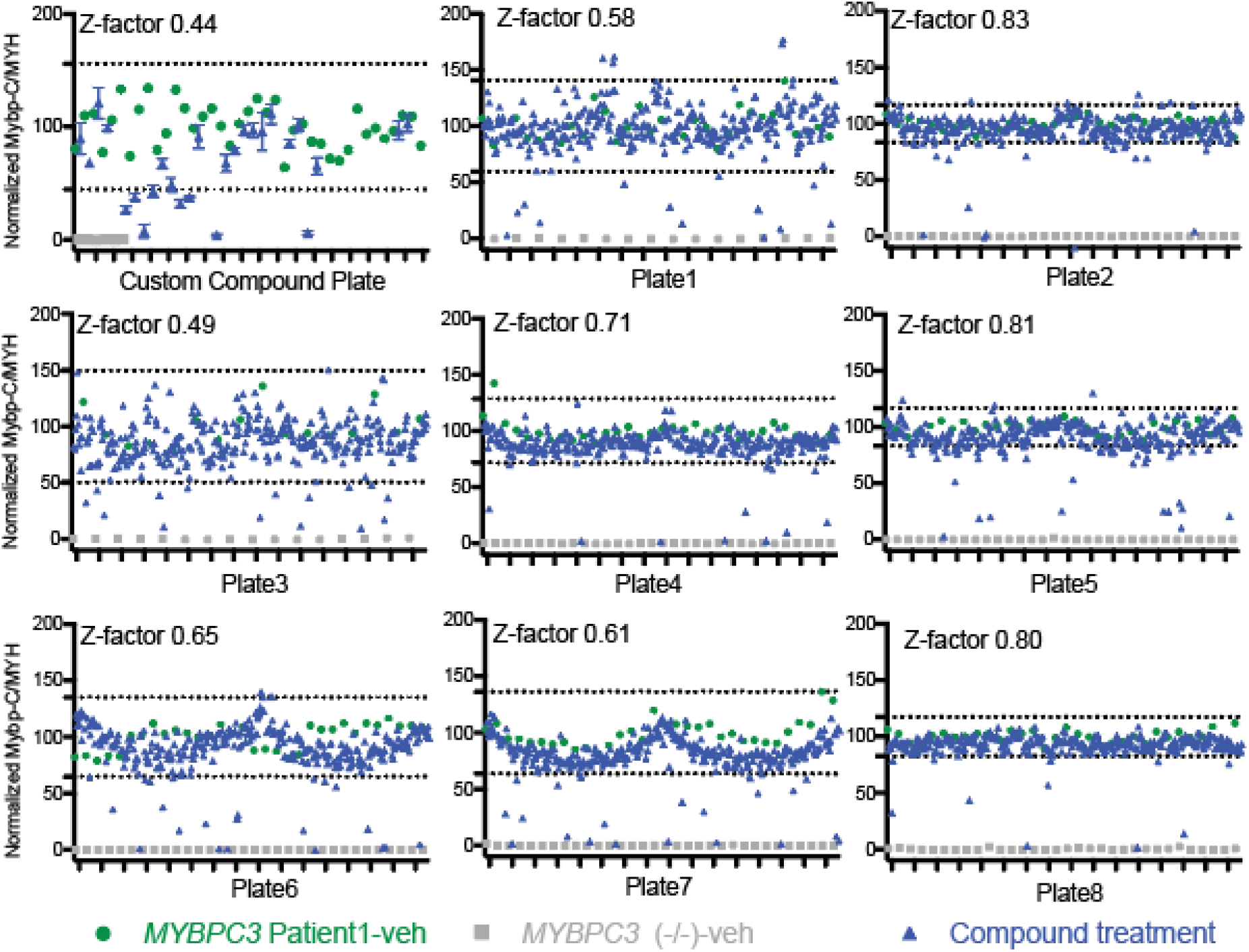
Full Screen results. Full screen results for every plate tested is shown. Results are normalized with positive control set at 100%-*MYBPC3* Patient1-veh (iPSC-CMs treated with vehicle control dmso), shown in green circles. The negative controls are set at 0%-*MYBPC3* (-/-)-veh, shown in grey squares. *MYBPC3* Patient1 iPSC-CMs treated with compound are shown in blue triangles. A Z-factor is calculated for each plate and hits are defined as 3 standard deviations above or below the mean of the positive control. This cut-off is marked by a dotted line for each plate.

**Supplemental Figure 7:**
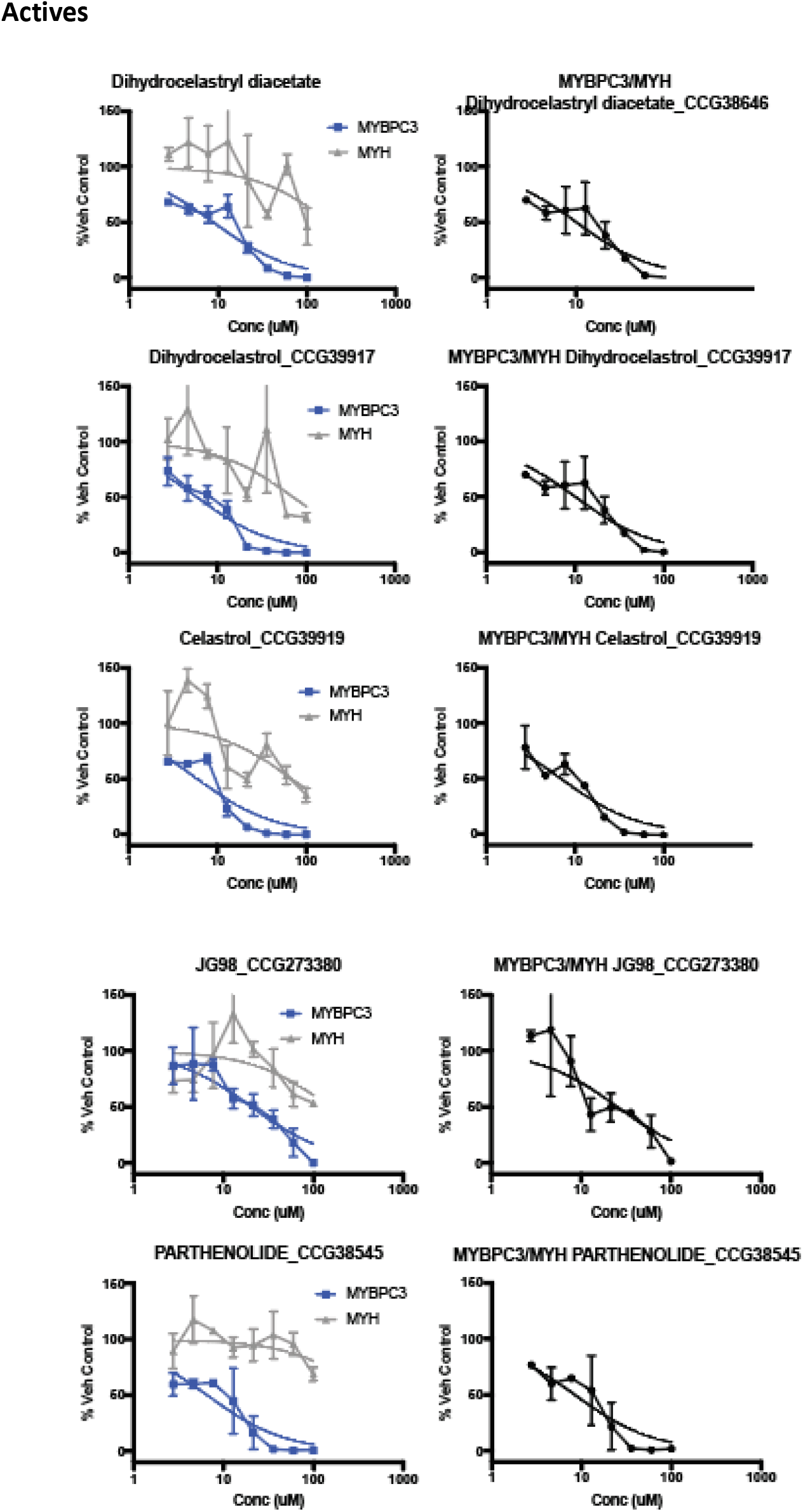
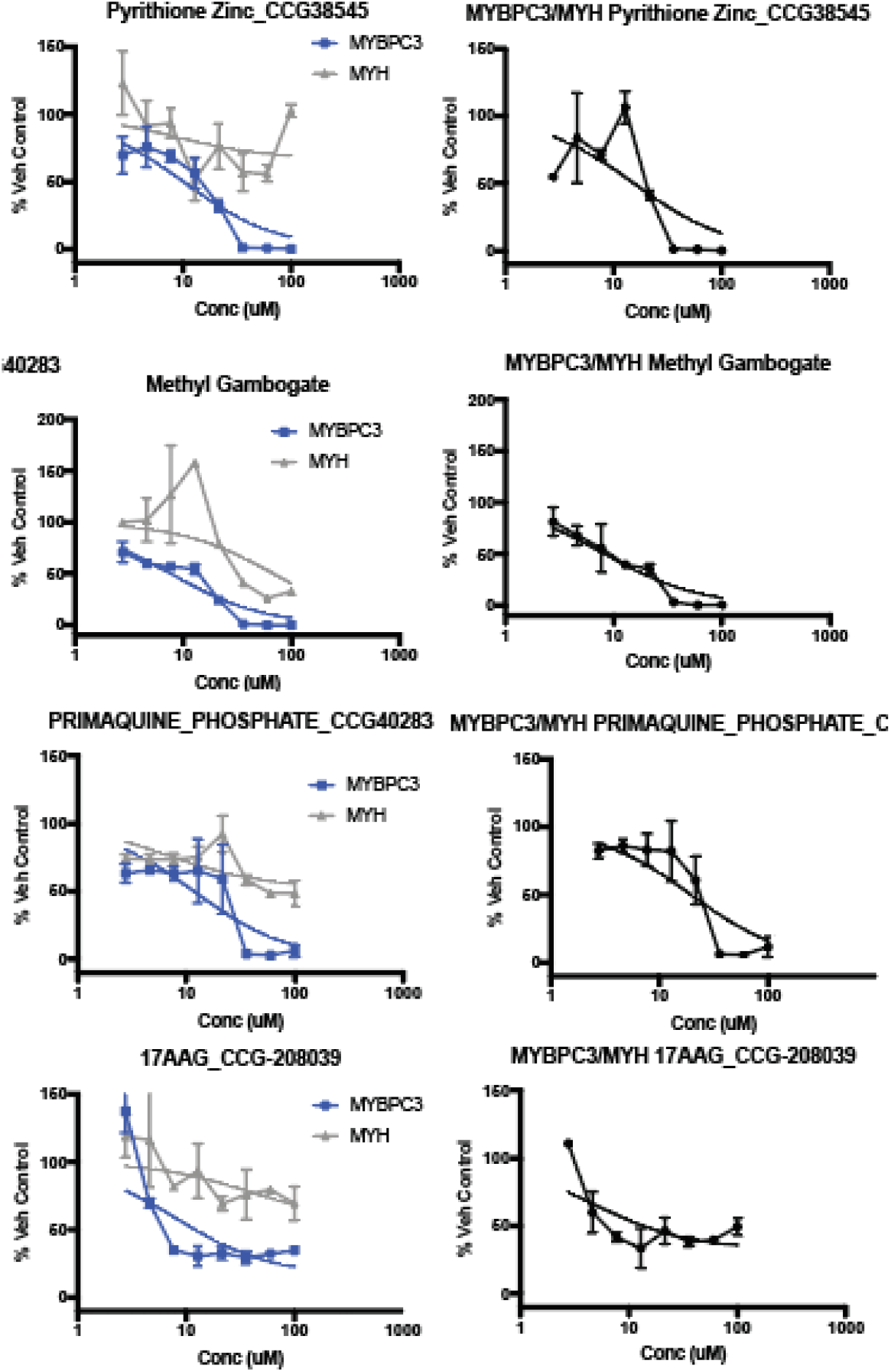
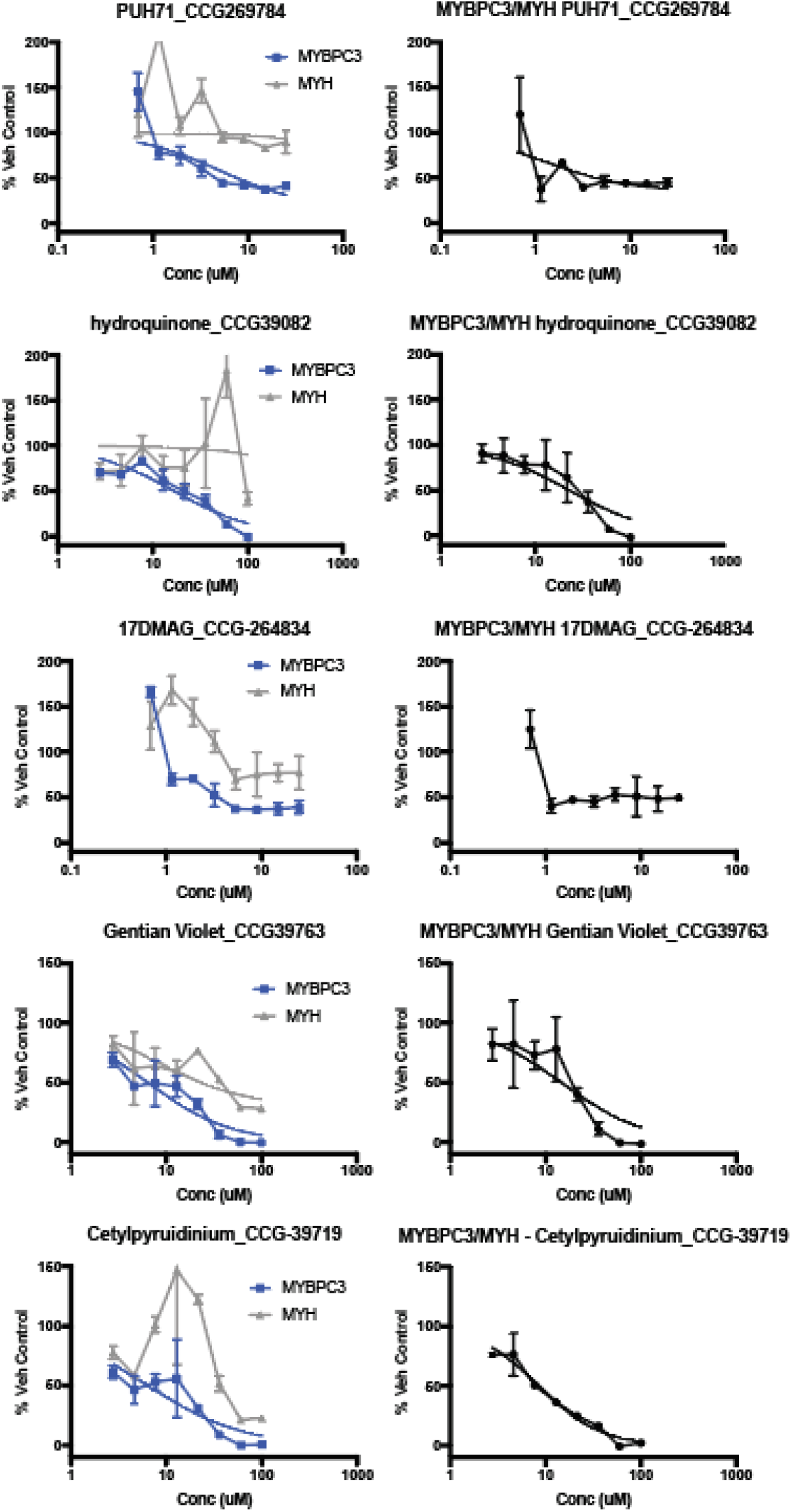
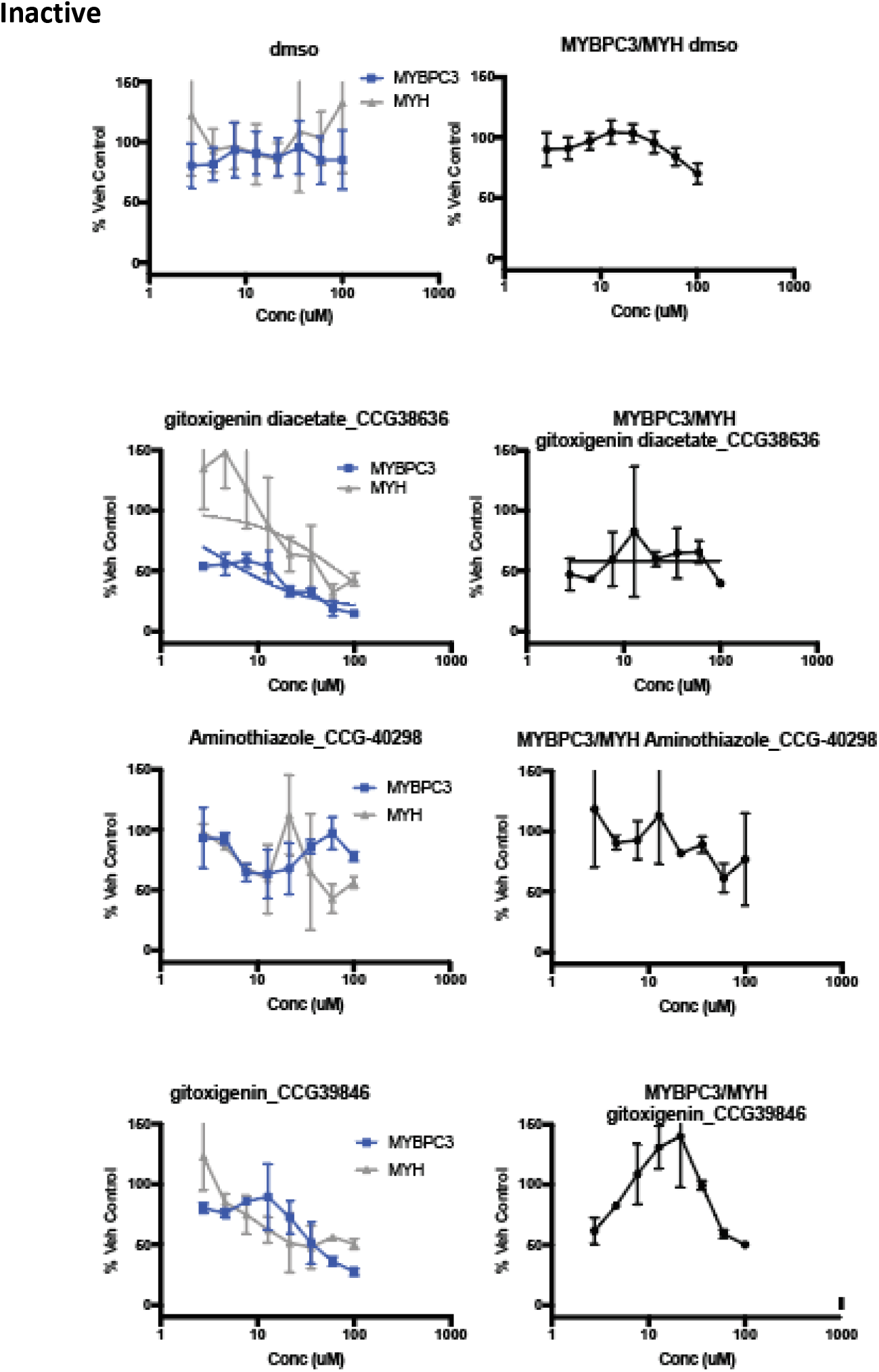
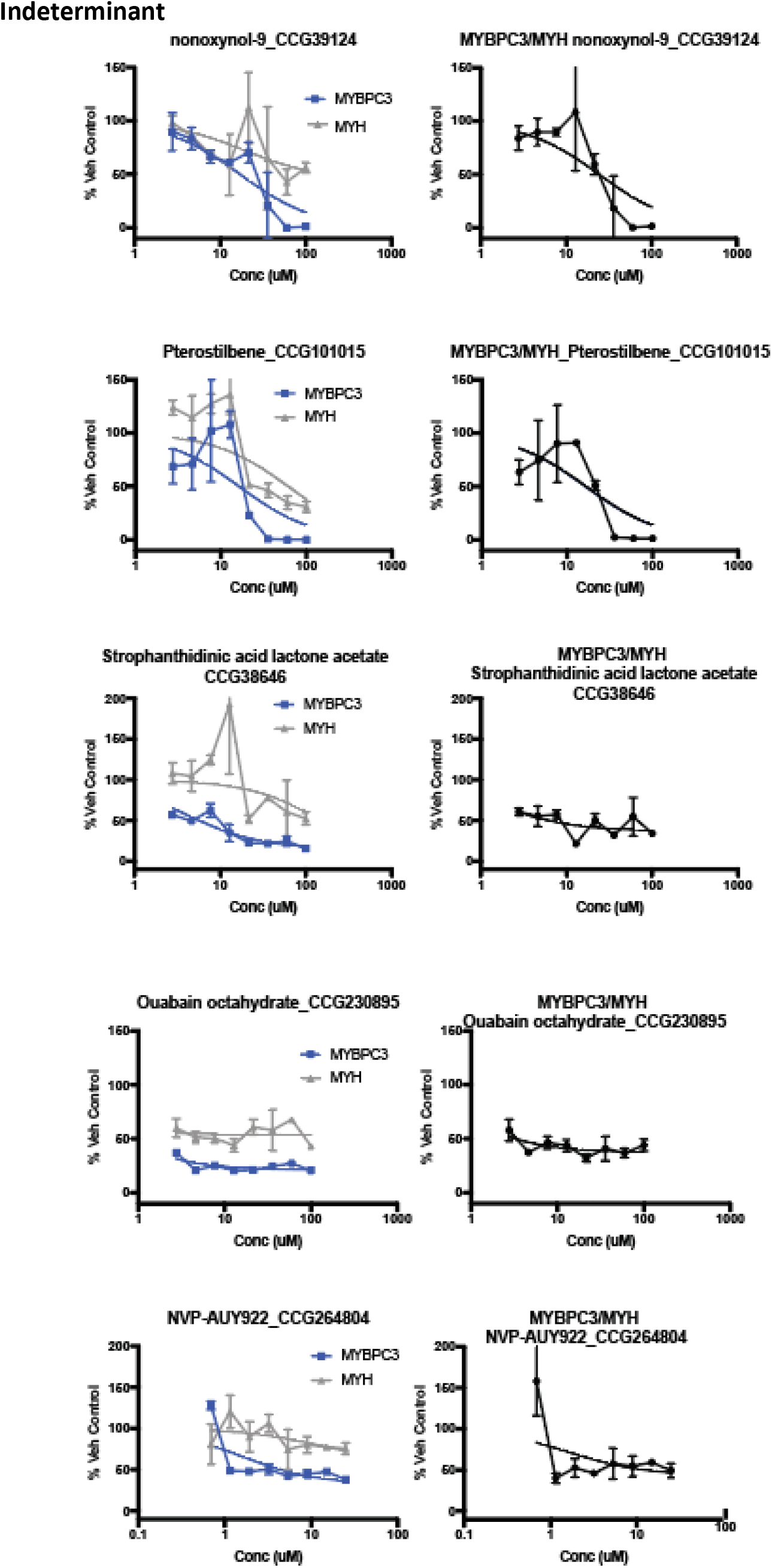
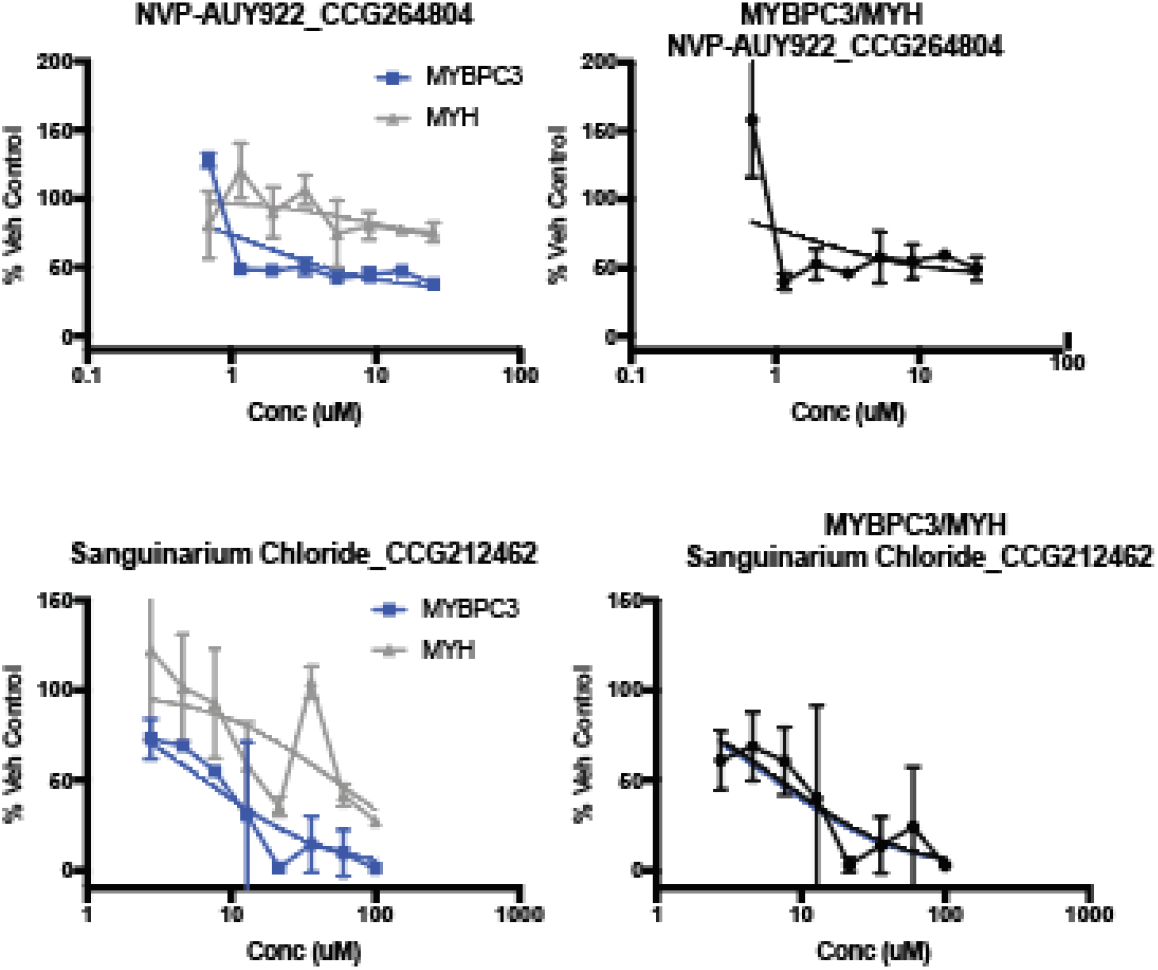
Full Concentration Response Curve (CRC) results-. *MYBPC3* Patient1 iPSC-CMs pooled from 3 independent differentiates were treated with compounds at the indicated concentrations (μM) tested in duplicate for 24 hours. MyBP-C and MYH Alpha-LISA was performed. Results were normalized to 100% *MYBPC3* Patient1 treated with vehicle control (dmso). 0% is set by the negative control. For MyBP-C and MyBP-C/MYH this was *MYBPC3* (-/-) treated with vehicle control (dmso). For MYH 0% is defined by a negative control of 293T cells treated with vehicle control (dmso). The mean and standard error of the mean for MyBP-C, MYH and the ratio of MyBP-C/MYH is shown for each compound. CRC results are shown for each of the 23 validated hit (Decreases MyBP-C/MYH, Decreases MyBP-C, does not decrease MYH). 14 compounds were identified as active. Concentration response curves were fit using GraphPad prism inhibitor vs response curves assuming bottom is greater than 0, top = 100, and hill slope −1. Results are further summarized in Supplemental Table 7.

**Supplemental Figure 8:**
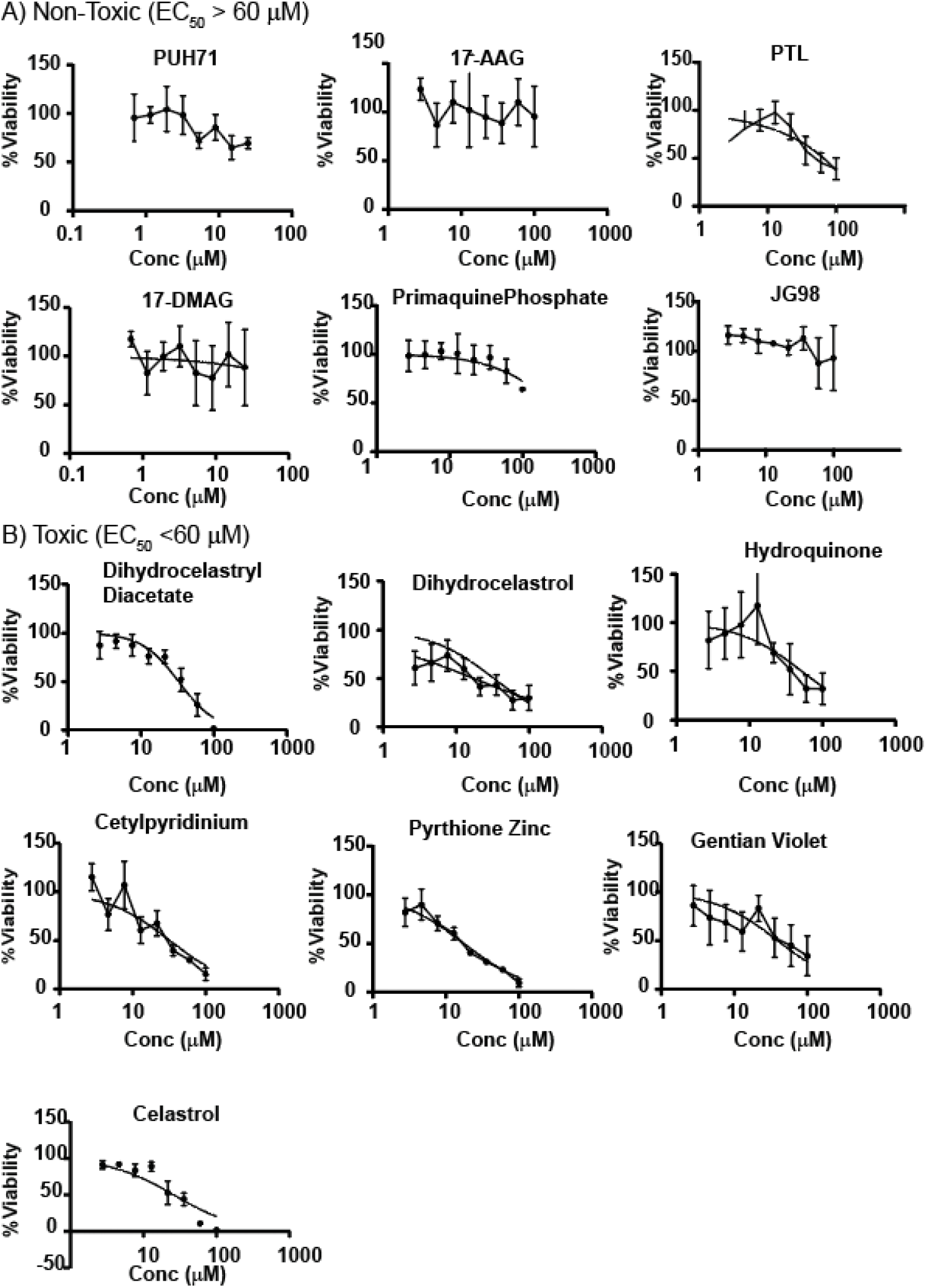
Full Cell Toxicity Results-. Tested in duplication on 2 biologic replicates of *MYBPC3* Patient1 iPSC-CMs (N=4). Results are normalized to %Viability, where 100% = *MYBPC3* Patient1 iPSC-CMs treated with vehicle control (dmso) and 0% = buffer only control (no live cells). Average and standard error of mean shown and fitted using GraphPad Prism inhibitor vs response non-linear fit assuming Top = 100, Bottom = 0 and hill-slope = −1. A) Non-toxic compounds have EC_50_ of > 60 μM. B) Toxic compounds have EC_50_ of <60 μM.

**Supplemental Figure 9:**
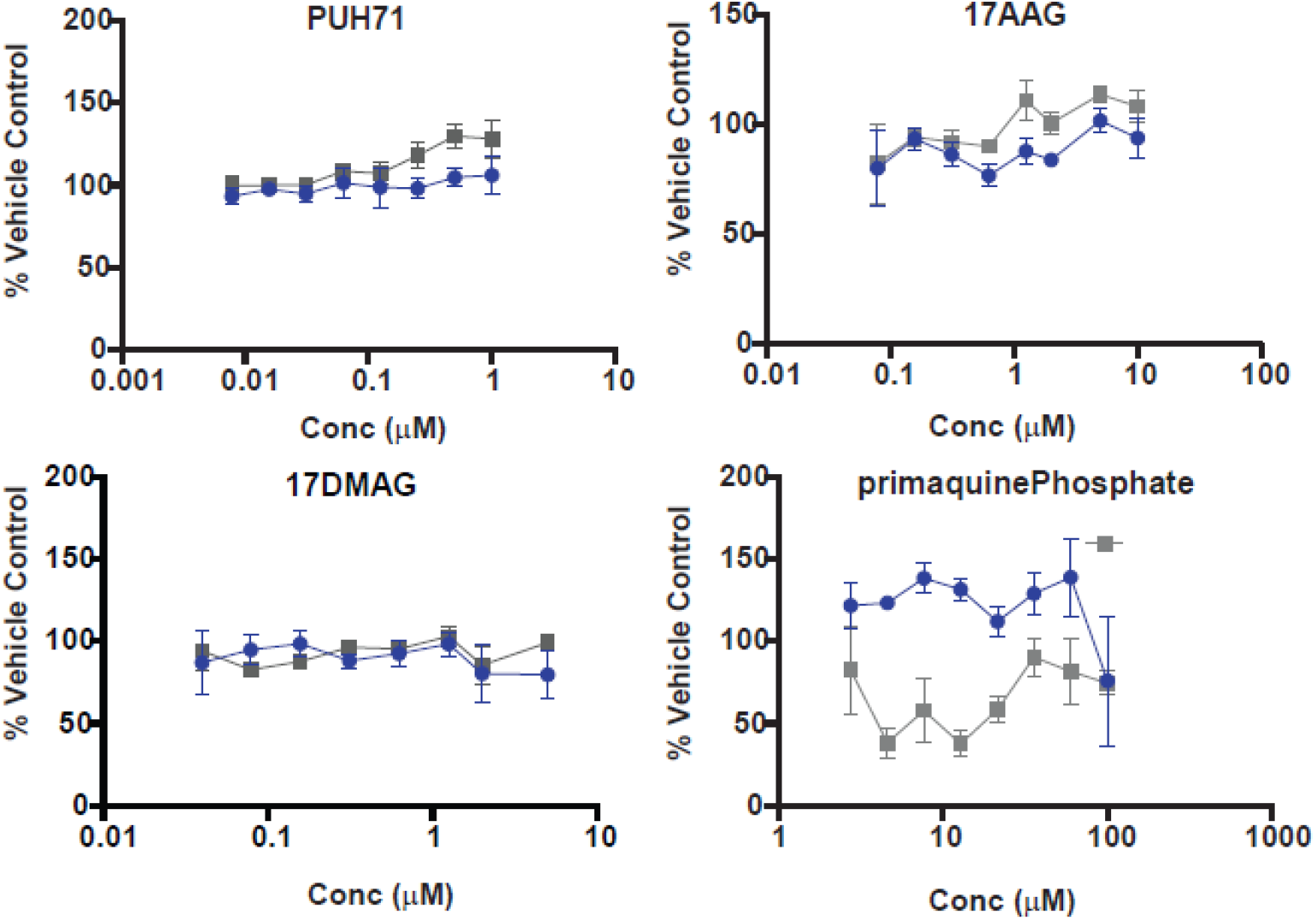
Inactive Hits from powder retest. 4/6 compounds lost activity observed during screening when fresh powder was tested (PUH71, 17-AAG, 17-DMAG, primaquine phosphate). The results of compound that were inactive during powder retest are shown above. All compounds were tested in duplicate on independently differentiated and pooled MYBPC3 Patient1 iPSC-CMs (n =4). Average and standard error of mean is depicted for each concentration tested. MyBP-C Alpha-LISA results are shown in blue circles and MYH Alpha-LISA result are shown in grey squares. Of note, the Hsp90 inhibitors did not exhibit typical dose dependent activity in CRC performed, but rather a rapid decrease in MyBP-C at lower concentrations (Supplemental Figure 7). As such a lower maximum concentration based on their known Hsp90 inhibitor EC_50_ was selected for their powder retest.

**Supplemental Figure 10:**
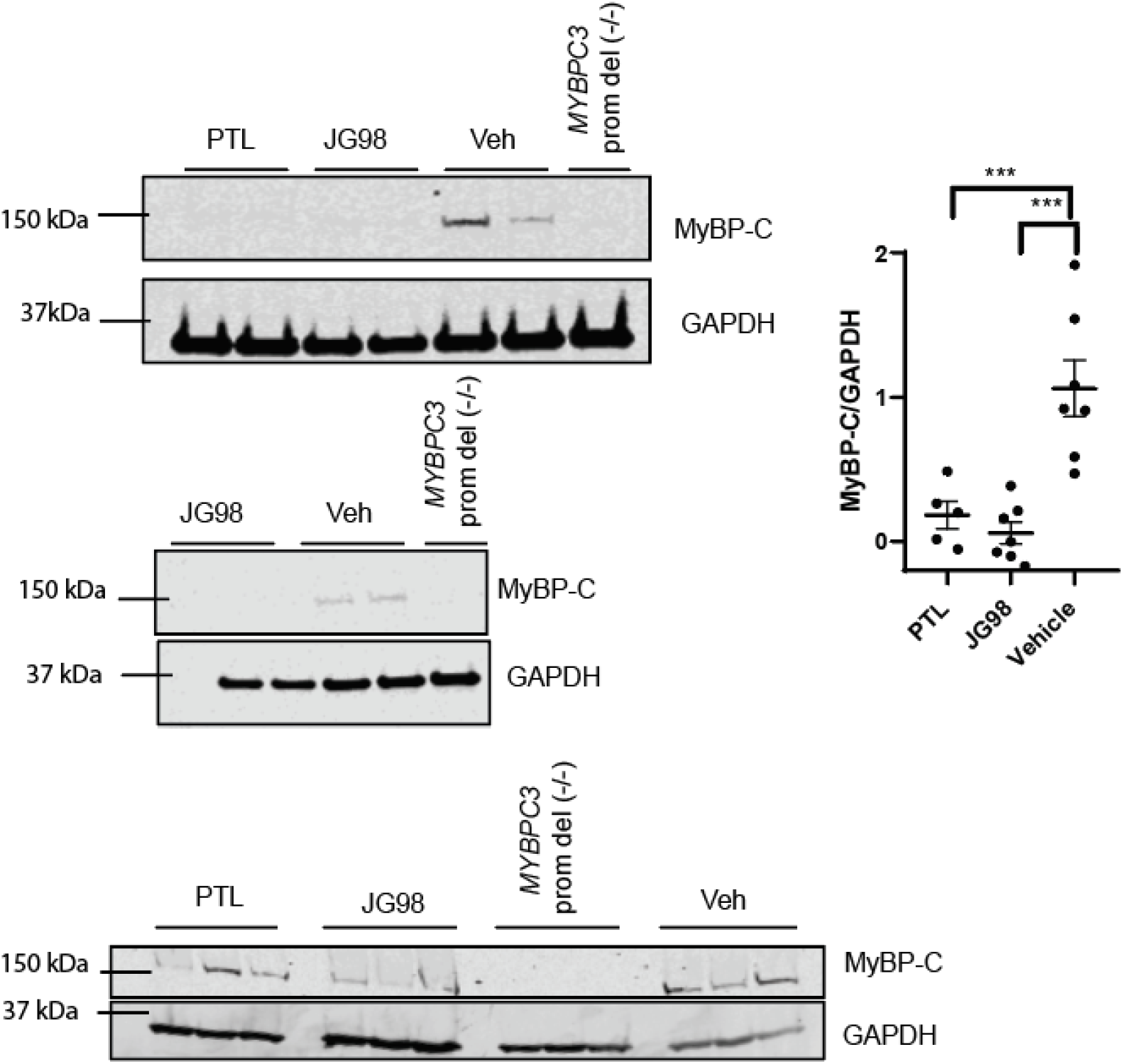
Western Blot confirmation of PTL and JG-98 Activity. Western blot using Mouse monoclonal *MYBPC3* (Santa Cruz sc-137180) on denaturing SDS-page gel were performed on cellular lysate from Ctrl-1 iPSC-CMs treated with JG-98, PTL at 20 mM or vehicle control are shown. was utilized. MyBP-C levels are normalized using GAPDH loading control. In all graphs, mean and standard error of the mean are shown, *** indicates p-value <0.0001.

**Supplemental Figure 11:**
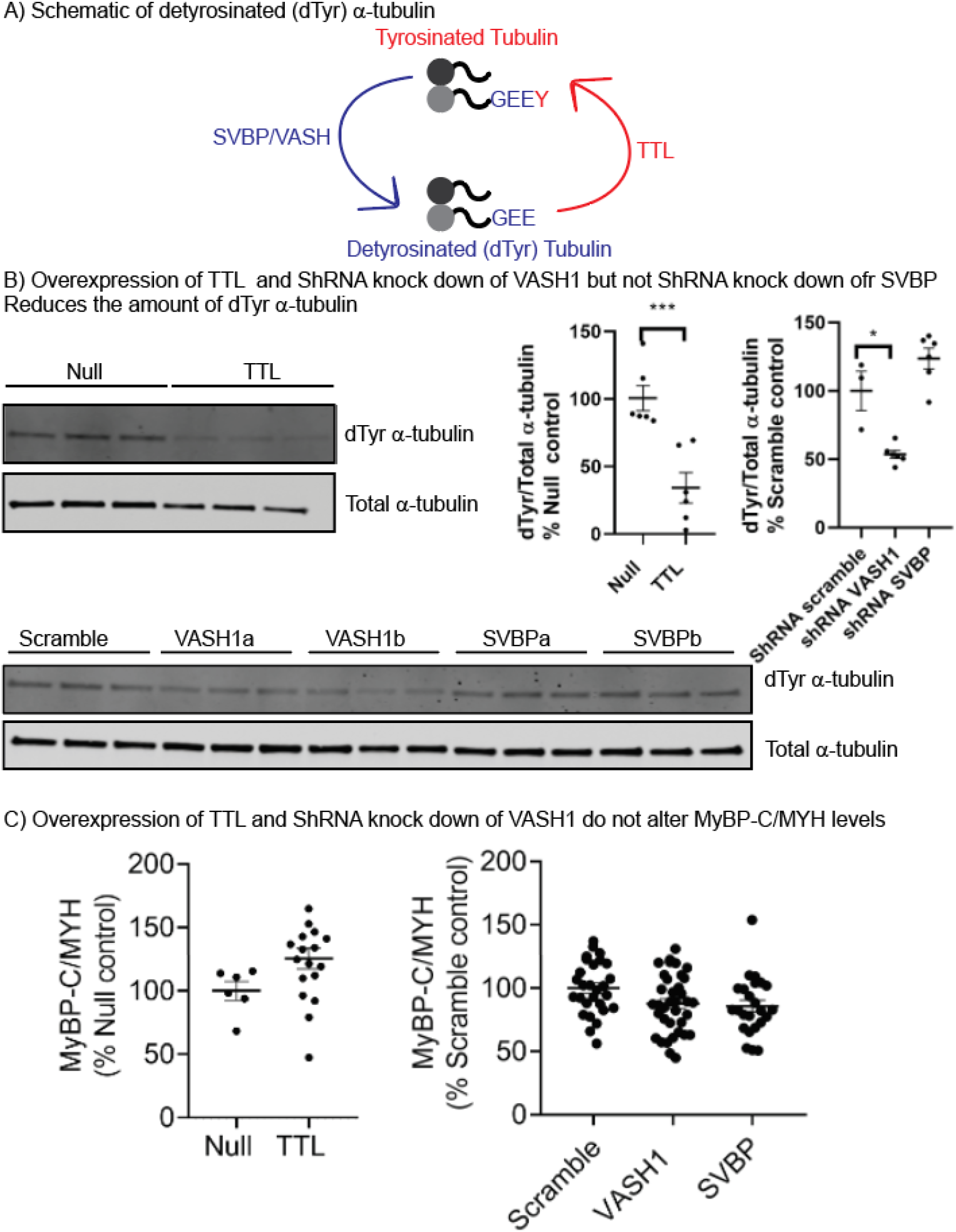
Genetic Experiments to evaluate if detyrosinated α-tubulin (dTyr) regulates MyBP-C protein homeostasis. (A) The proteins that drive tyrosination and detyrosination of α-tubulin are depicted. B) *MYBPC3* Patient3 iPSC-CMs were treated with adenovirus MOI 10 carrying a null control, the enzyme TTL, or shRNA (Scrambled, or targeting-VASH1, SVBP). Two shRNA constructs were used for both VASH1 and SVBP (a,b).(14) Treated cells were lysed and analyzed by (B) western blot to evaluate their effect on dTyr-α-tubulin or (C) Alpha-LISA to evaluate their effect on MyBP-C/MYH protein levels. Graphs show mean and standard error of the mean, p-value is inducated by; * <0.05, ** < 0.005, *** <0.0005.

**Supplemental Table 1:**
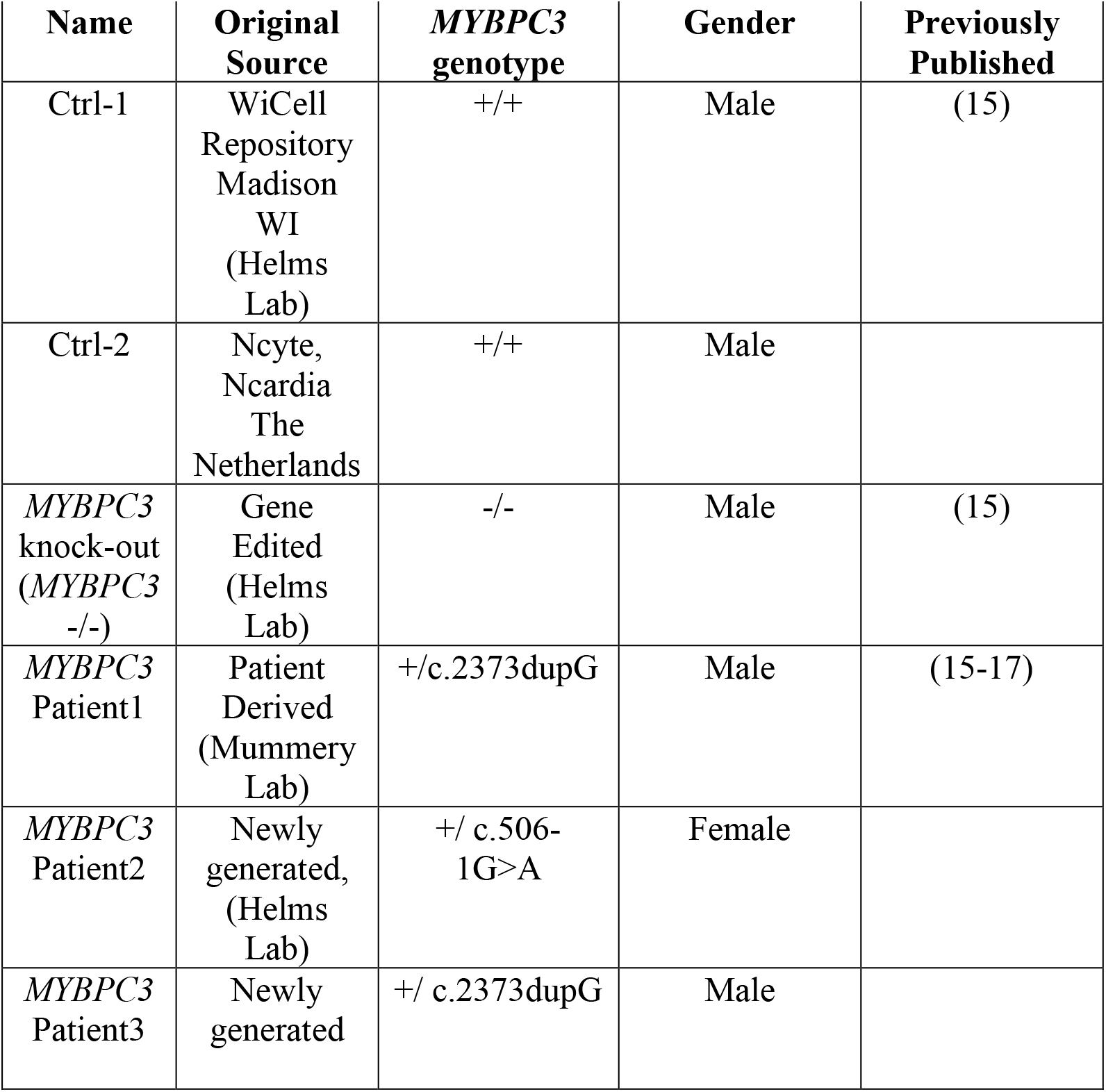
Induced pluripotent stem cell (iPSC lines)

**Supplemental Table 2:**
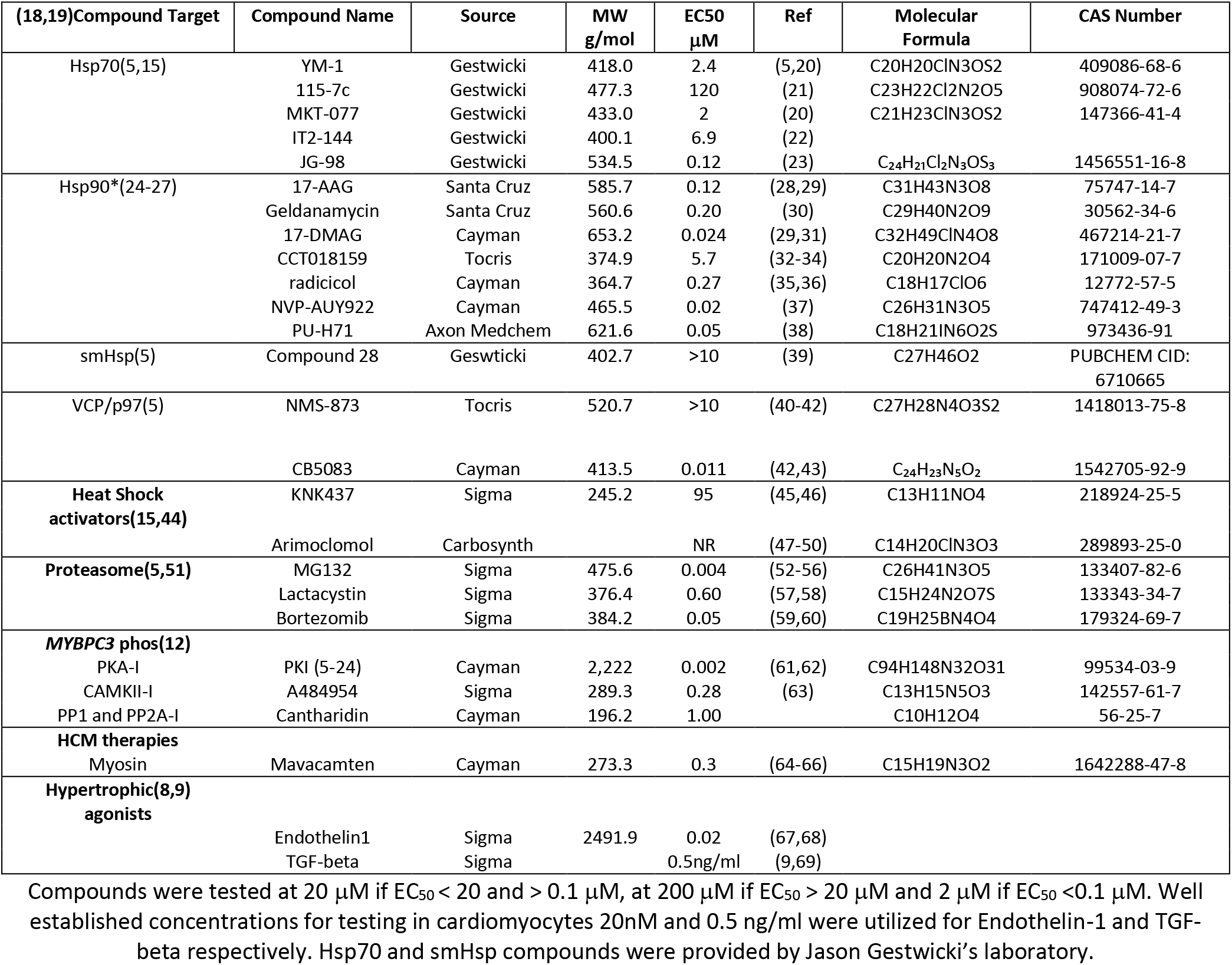
Custom Compounds-MyBP-C Homeostasis Literature Review.

**Supplemental Table 3:**
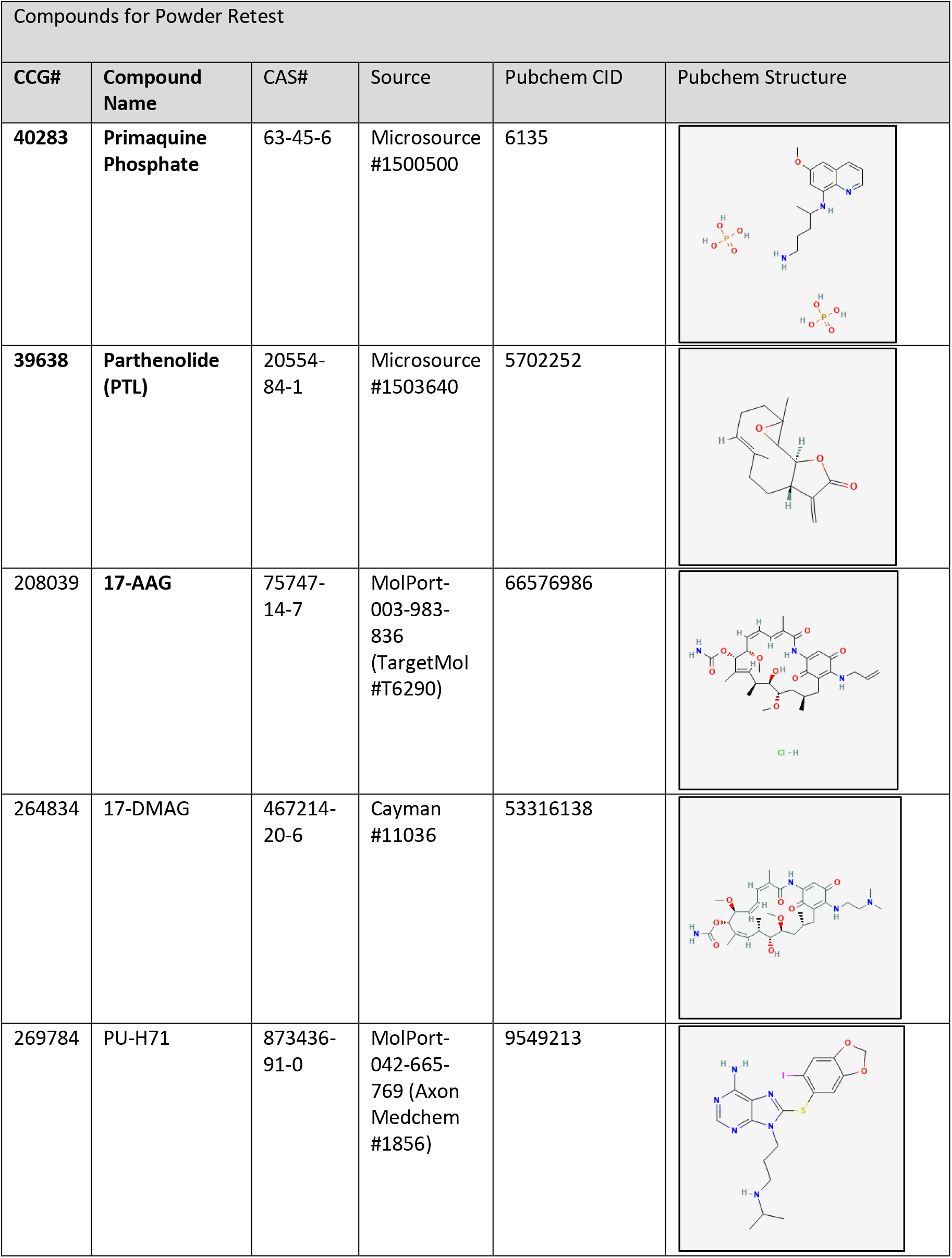
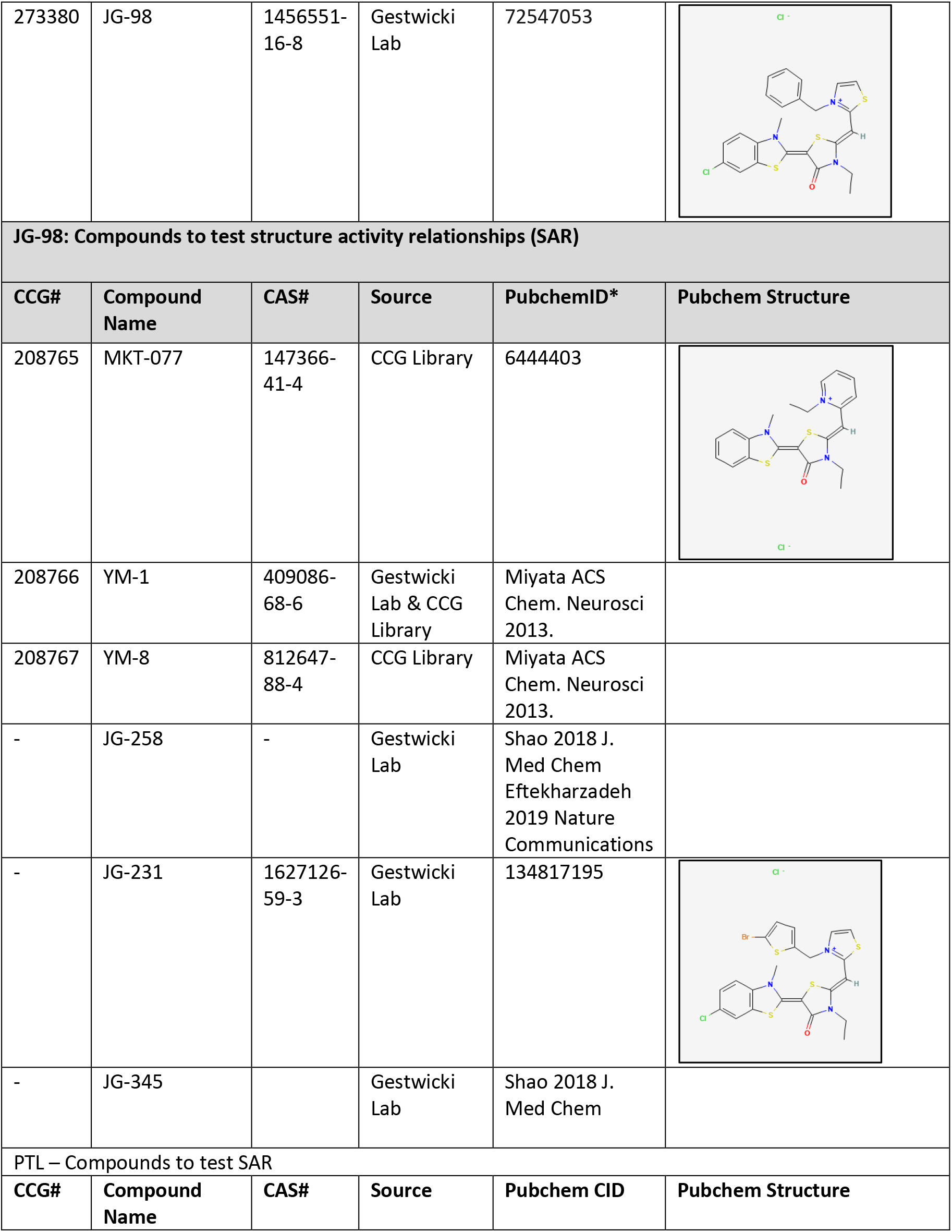
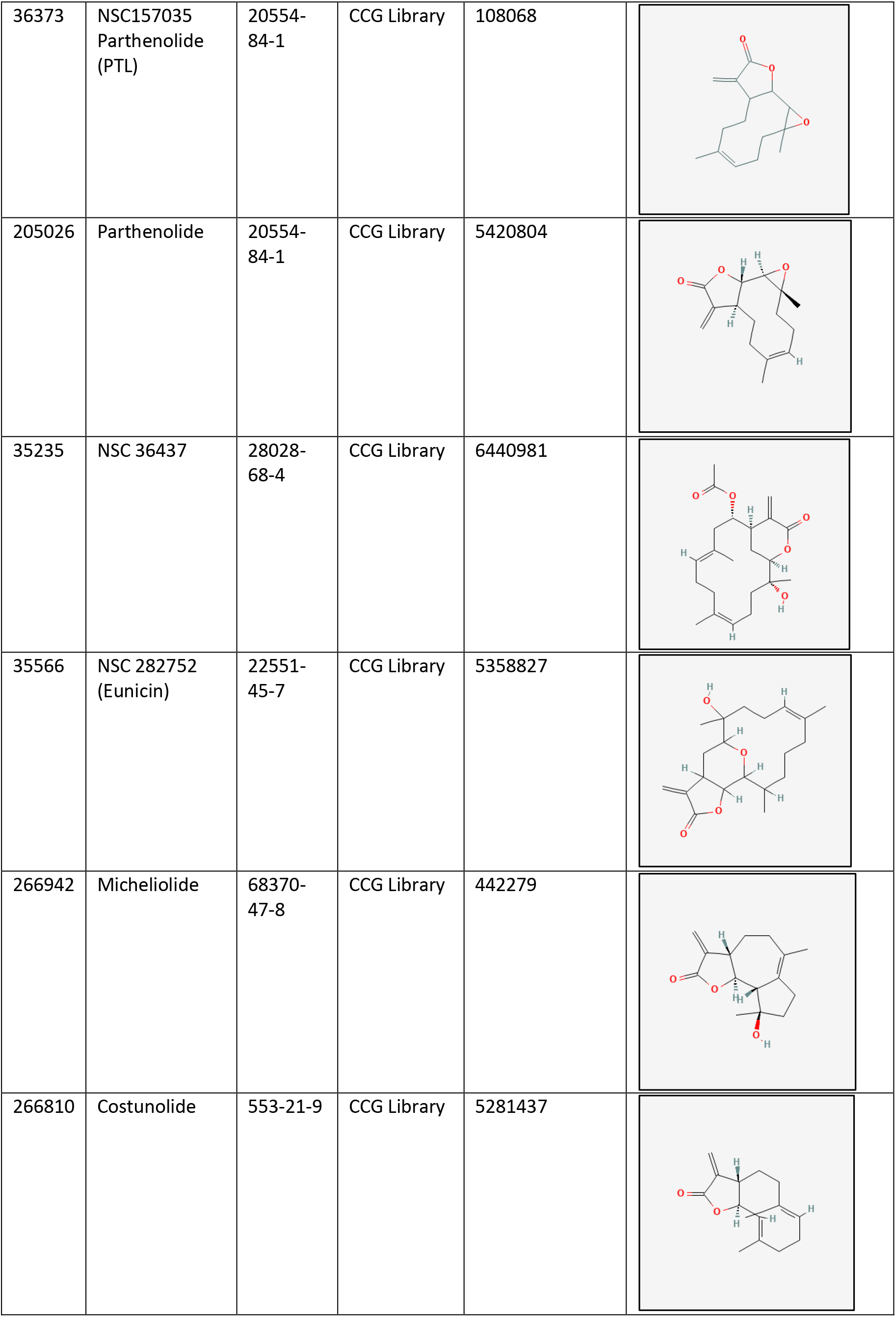
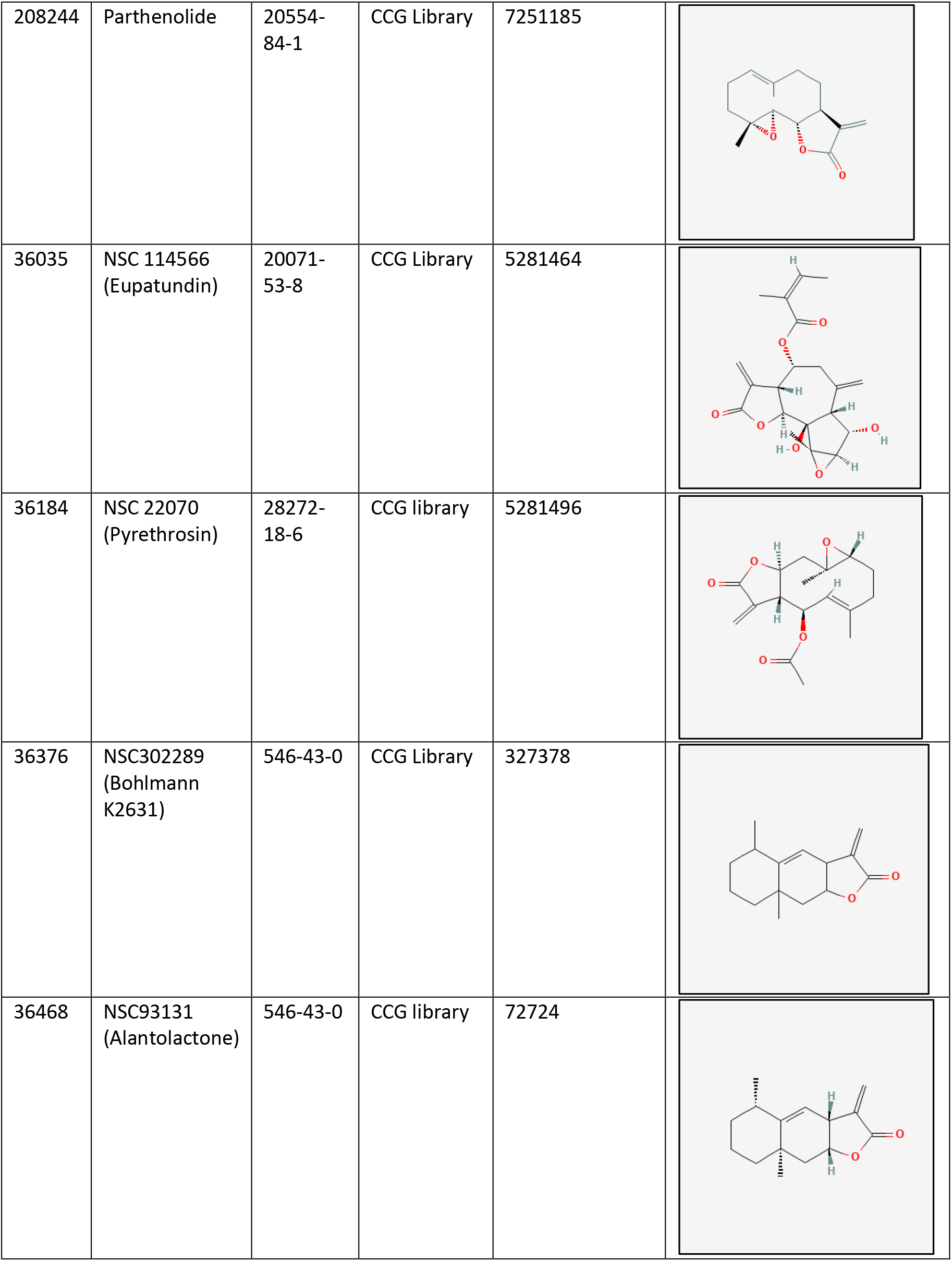
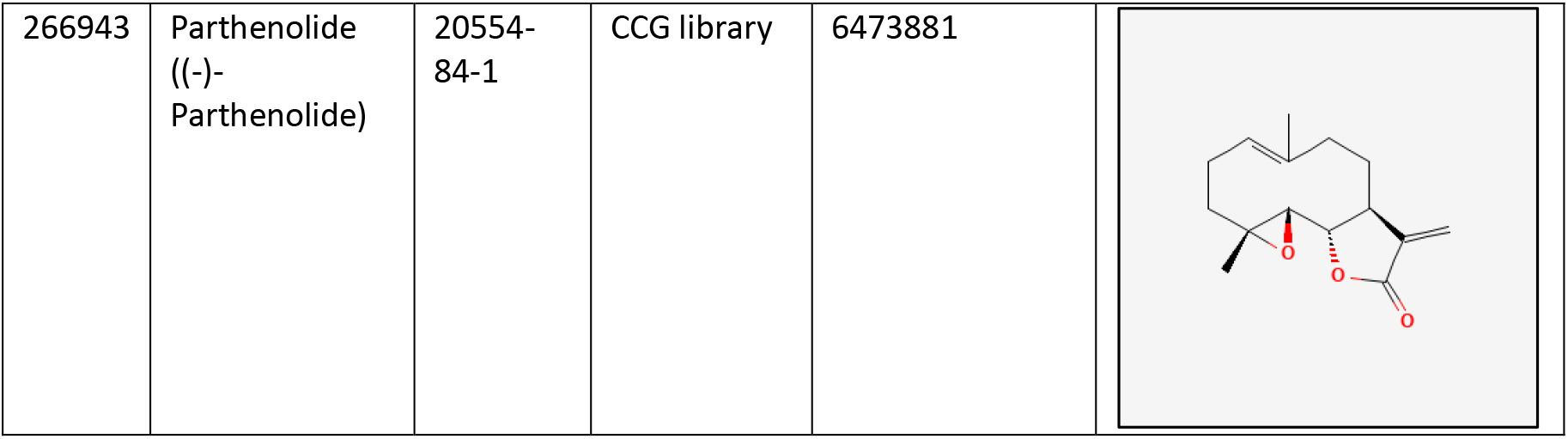
Compounds for Powder Retest and Structure activity relationships.

**Supplemental Table 4:**
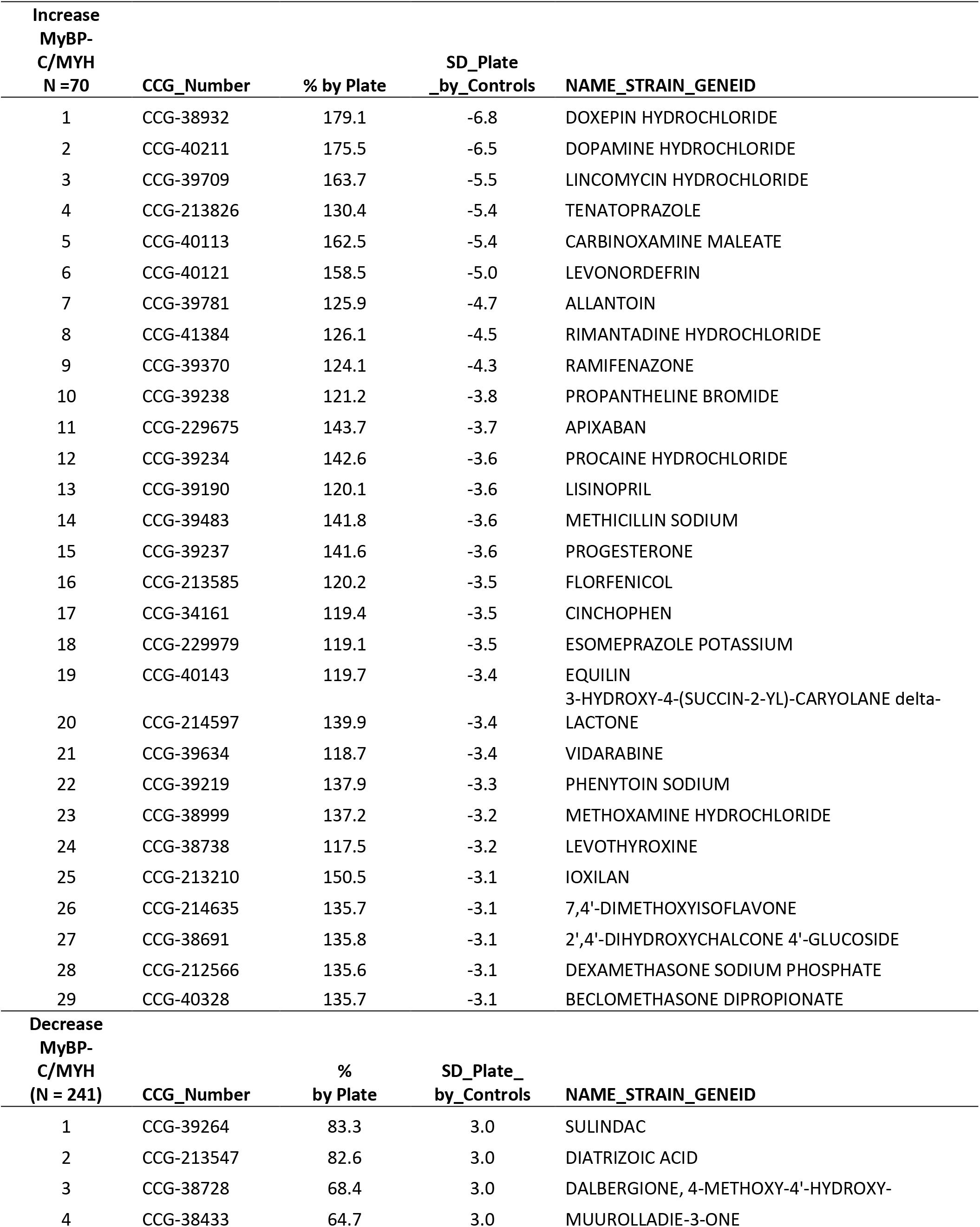
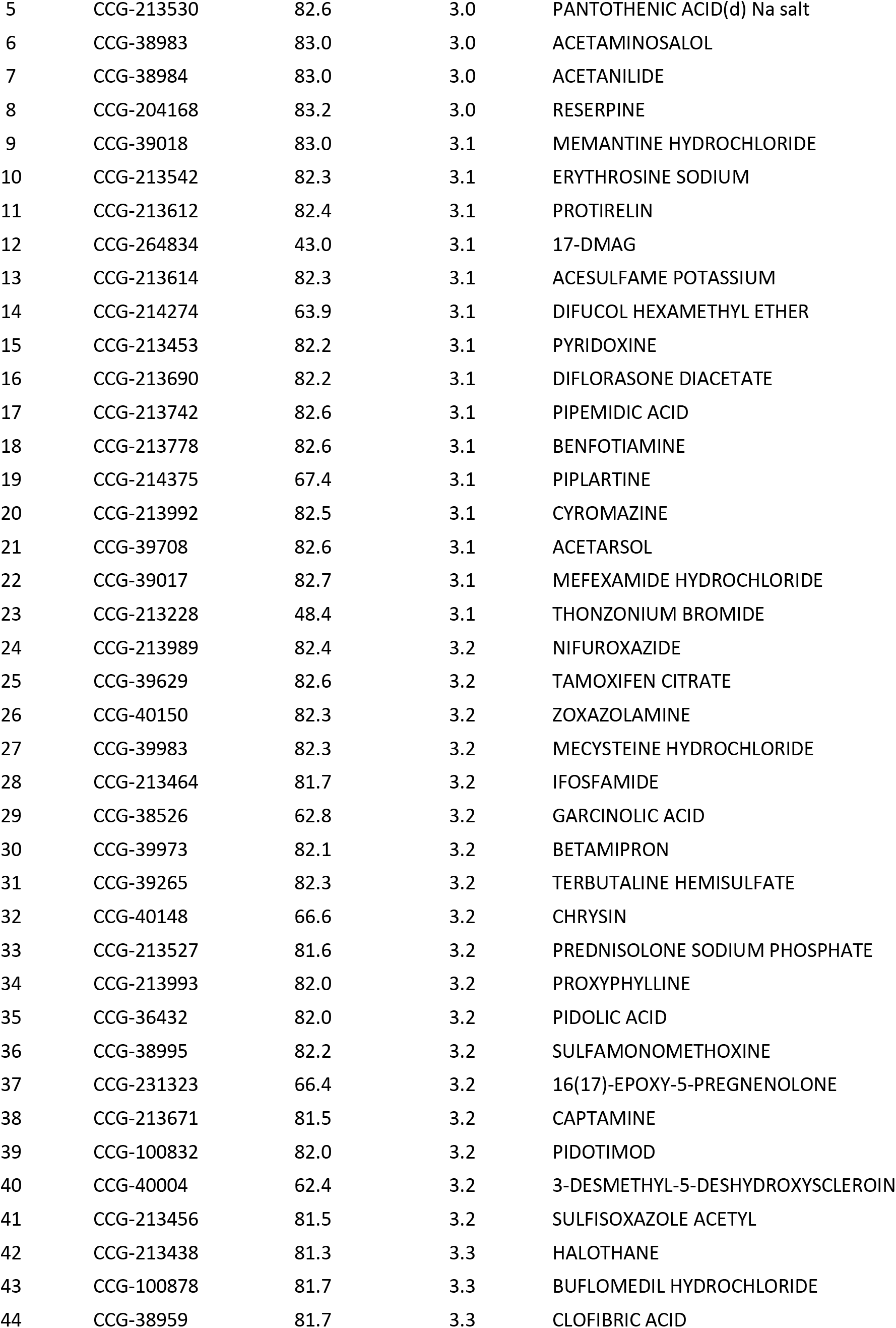
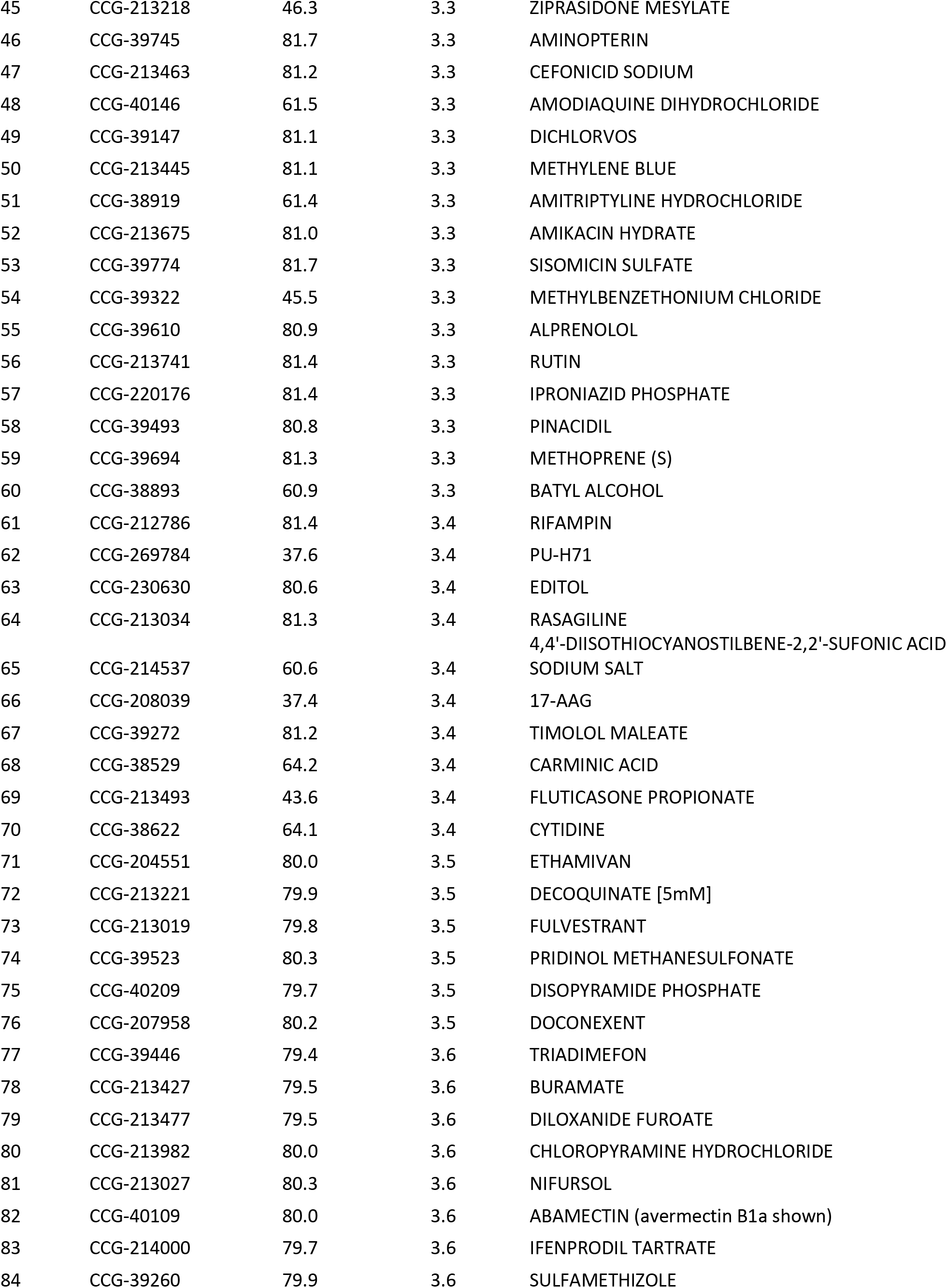
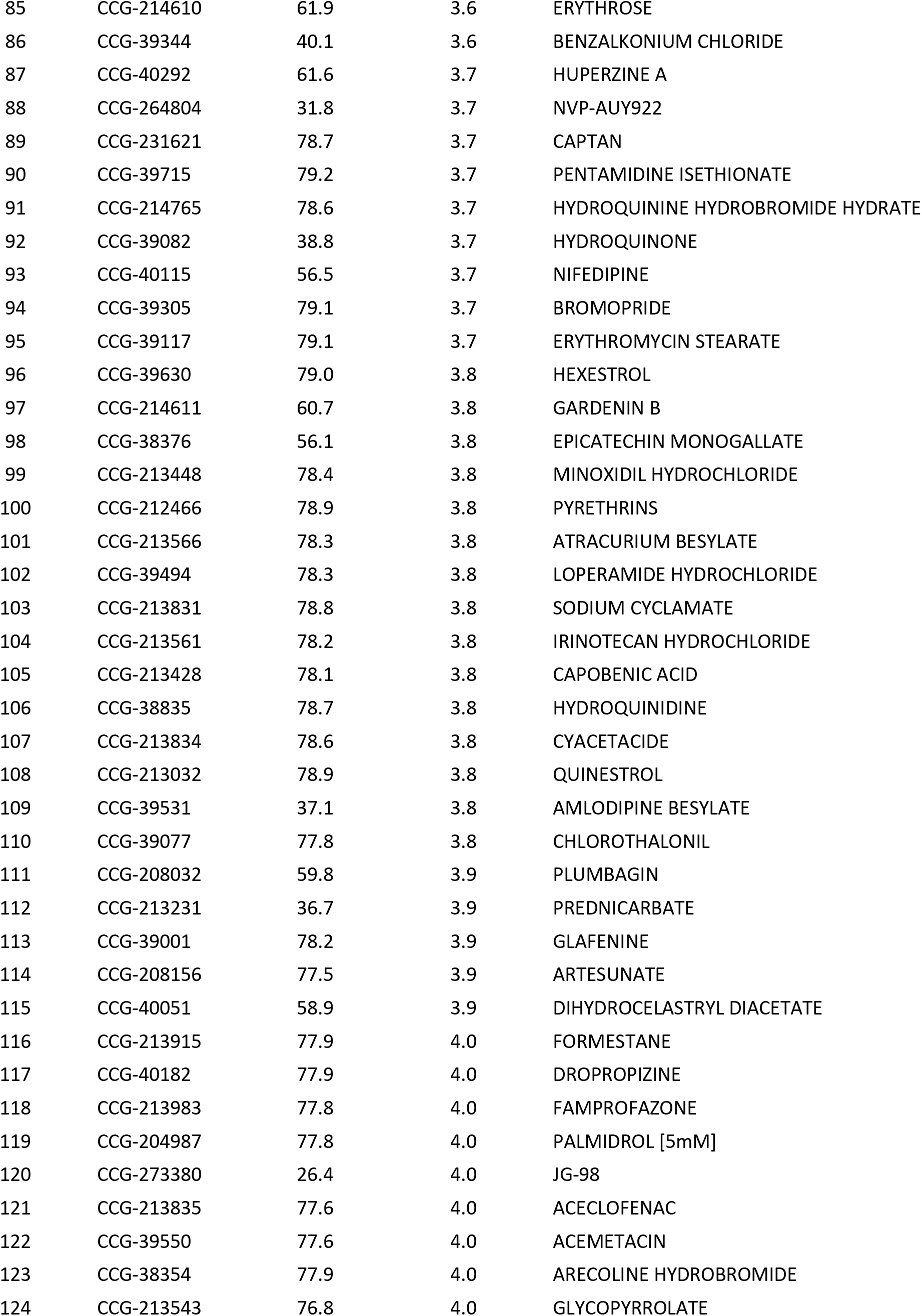
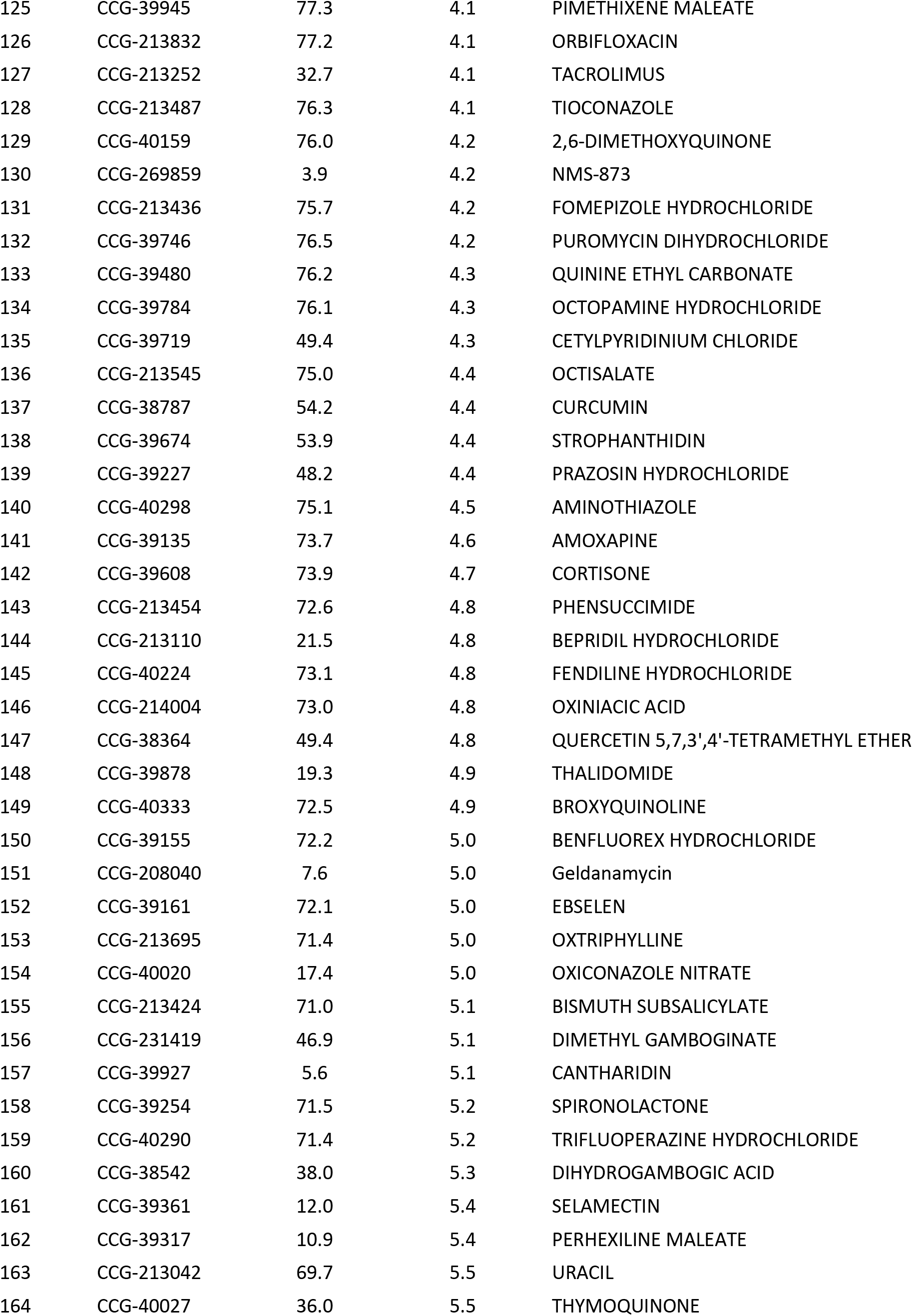
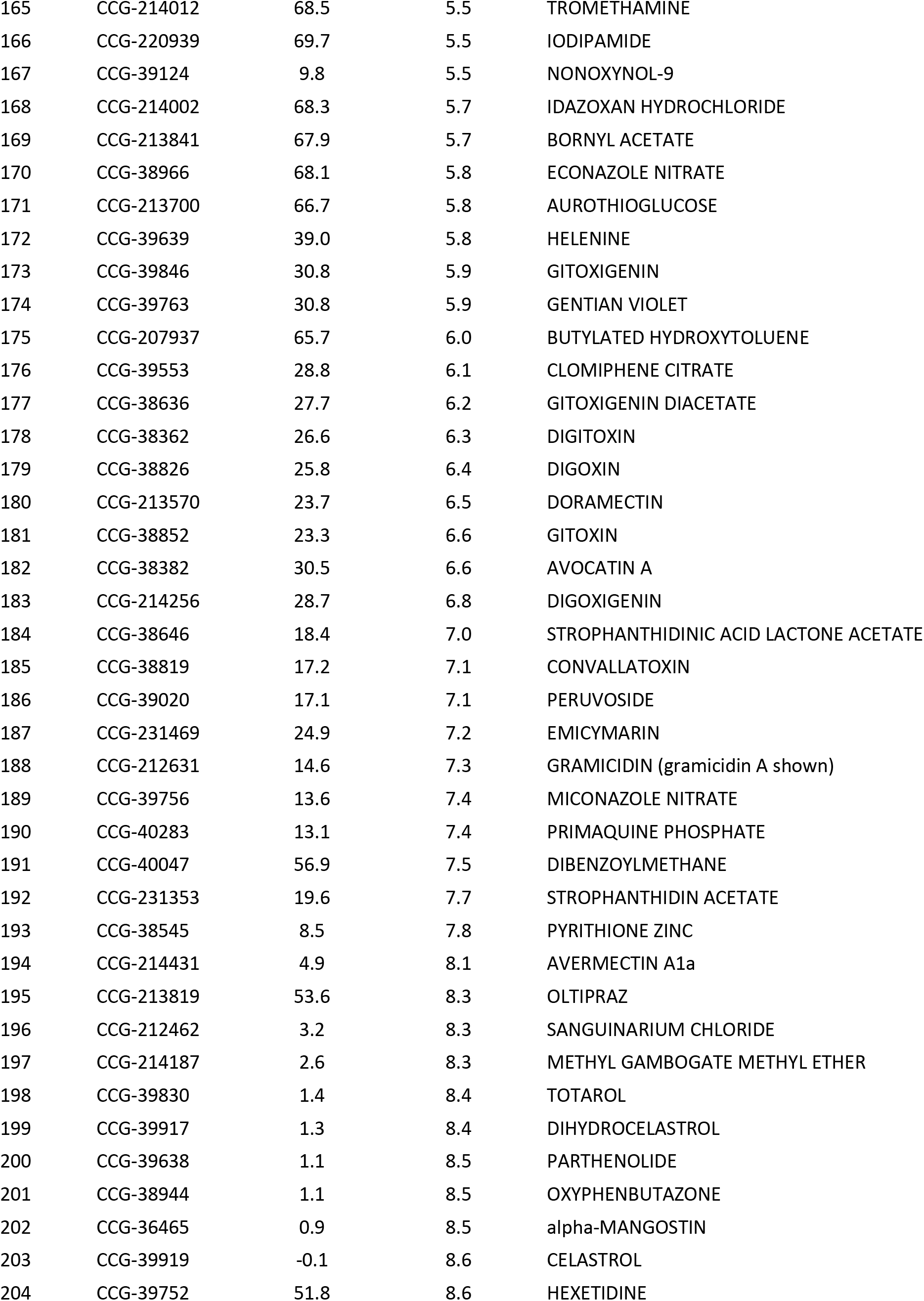
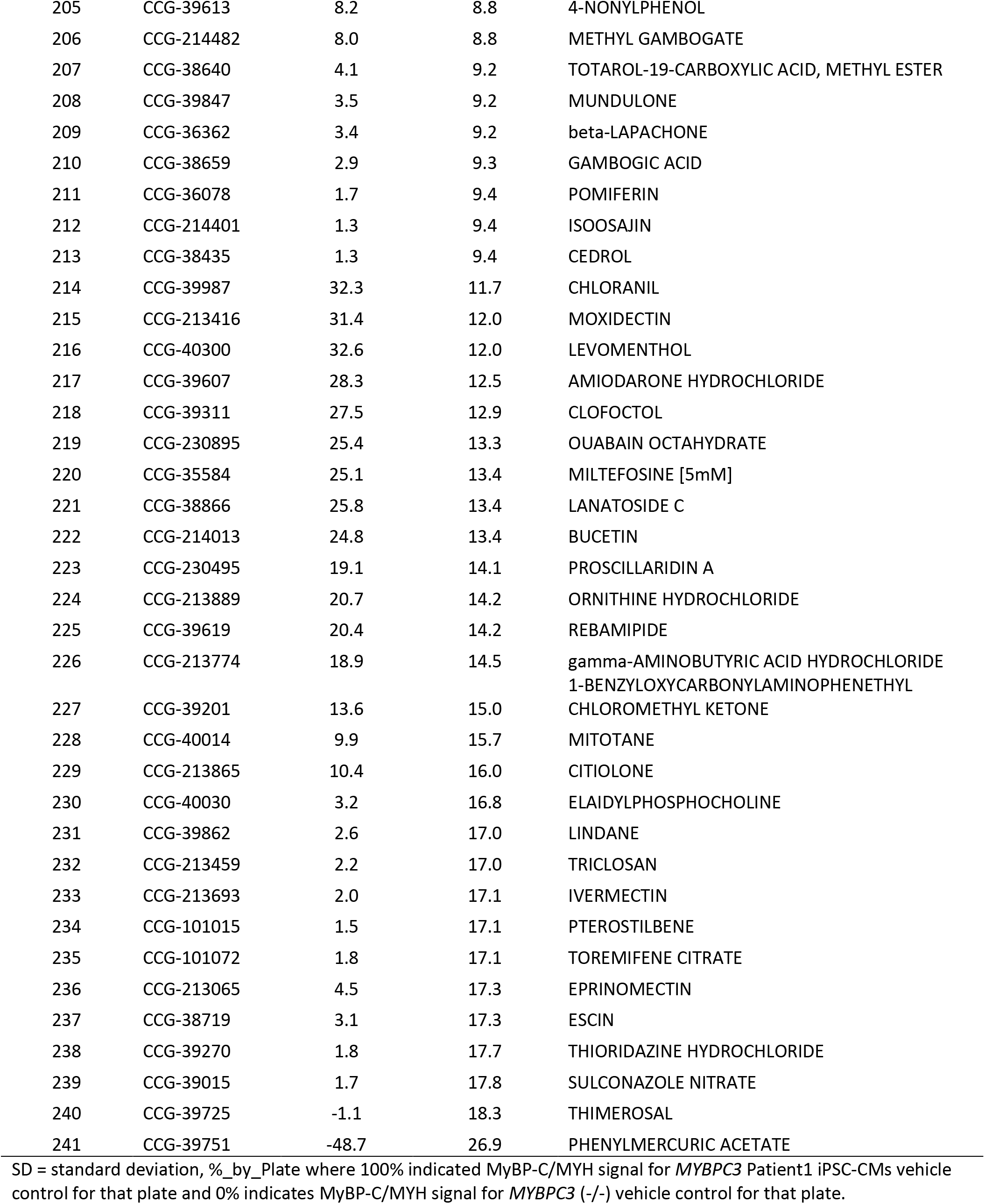
Initial Hits.

**Supplemental Table 5:**
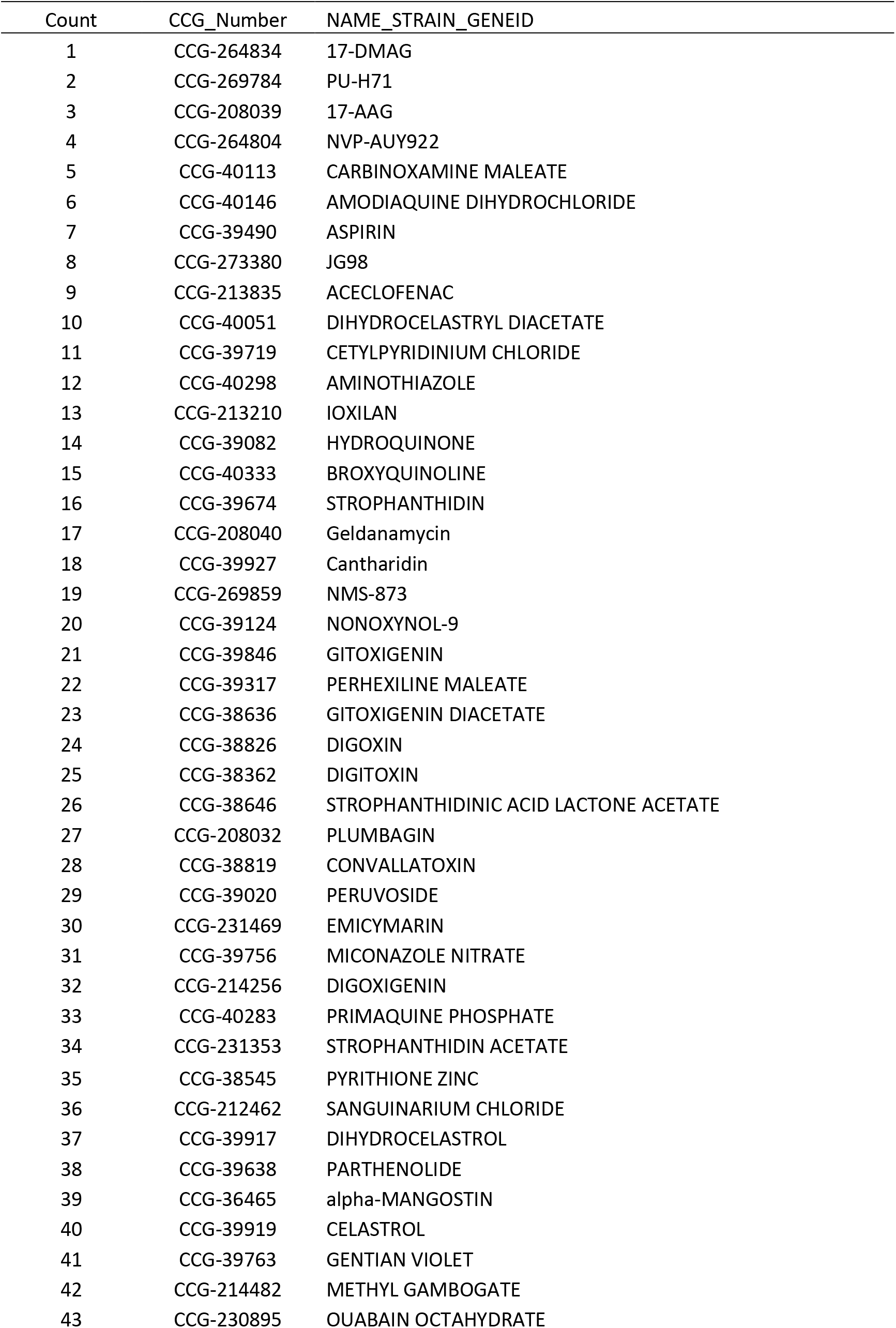
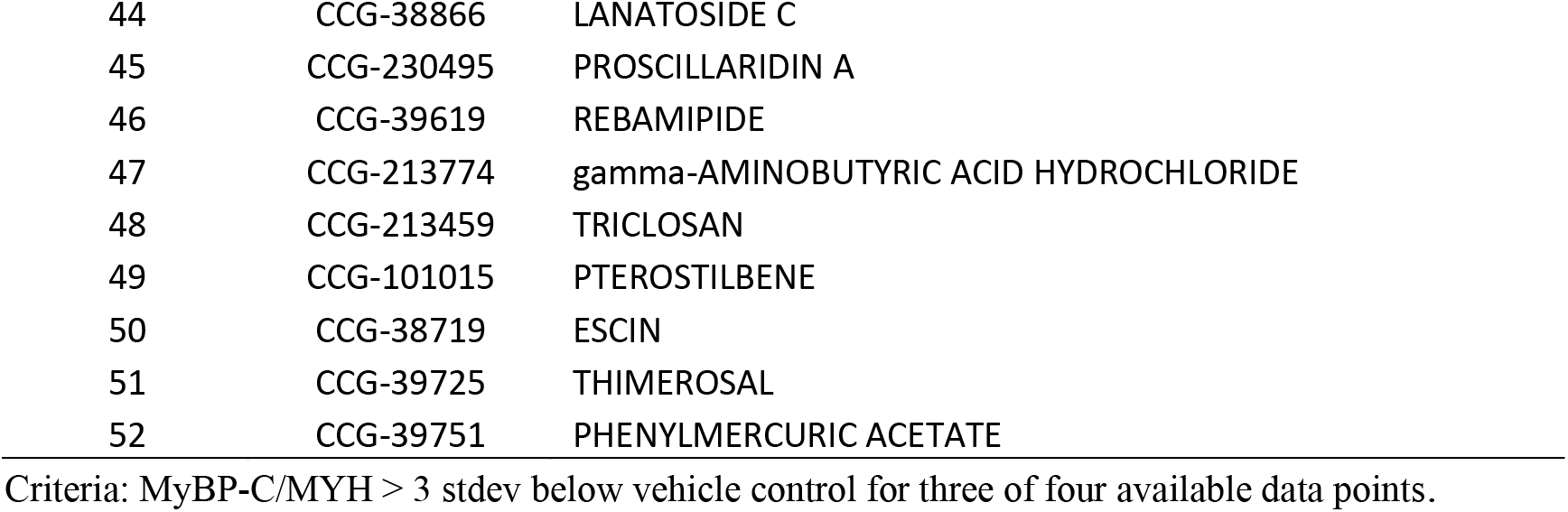
Validated by retesting – Compounds that decrease MyBP-C/MYH.

**Supplemental Table 6:**
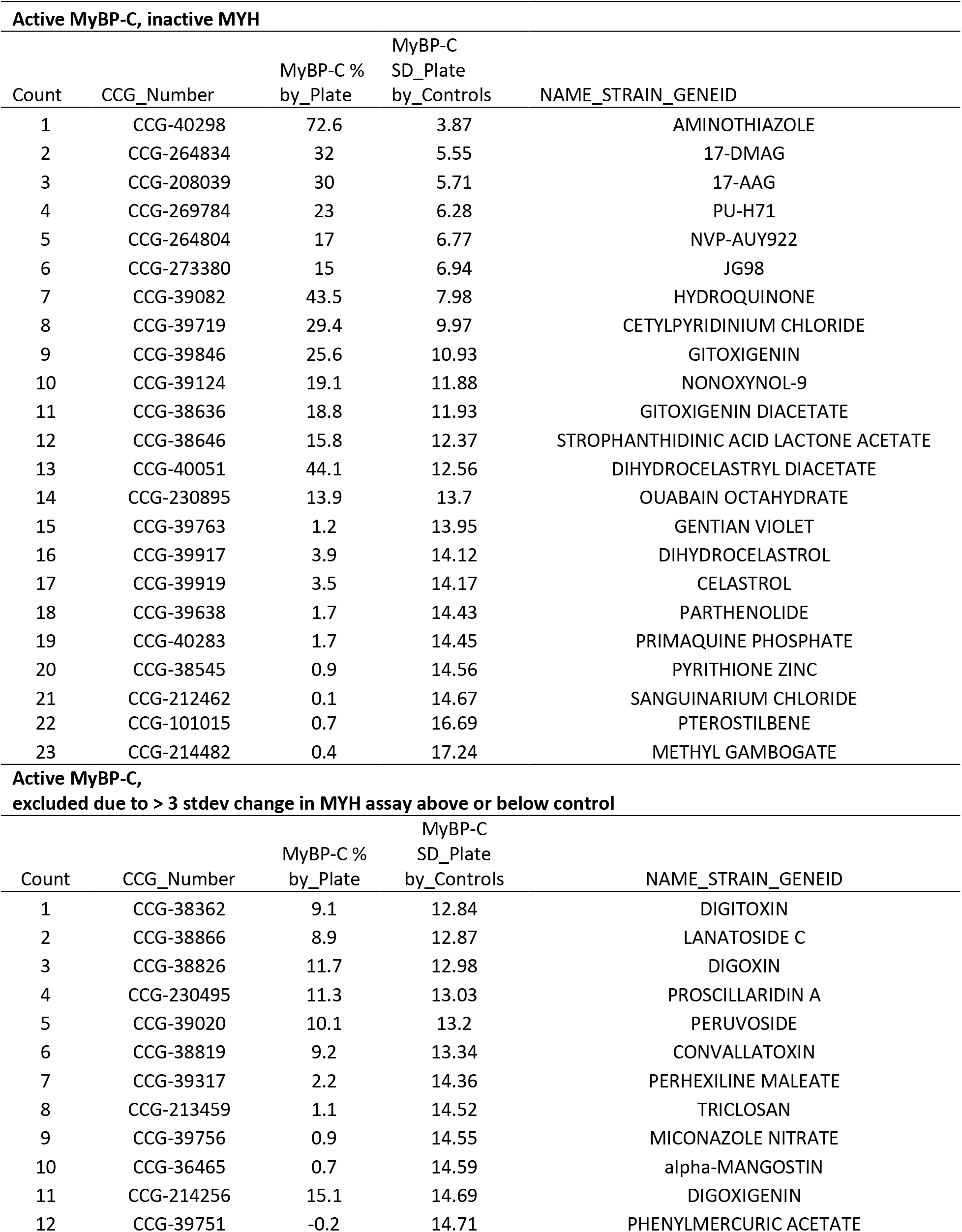
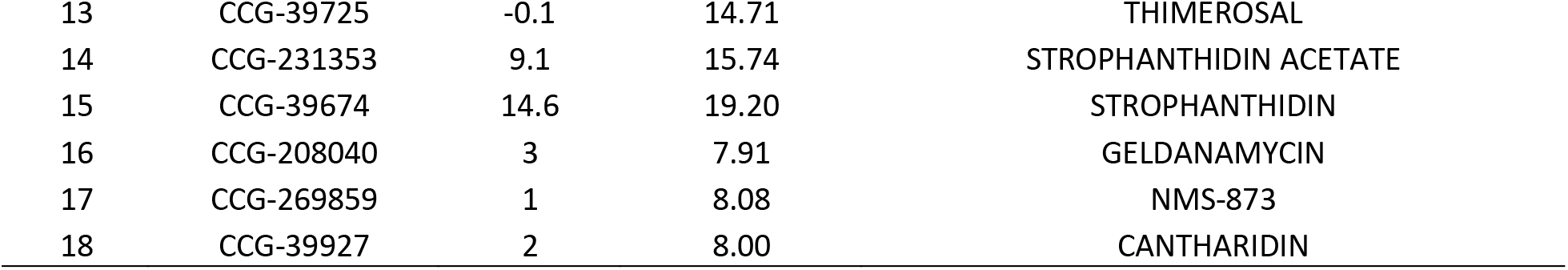
Evaluating MyBP-C and MYH assay results for validated MyBP-C/MYH compounds.

**Supplemental Table 7:**
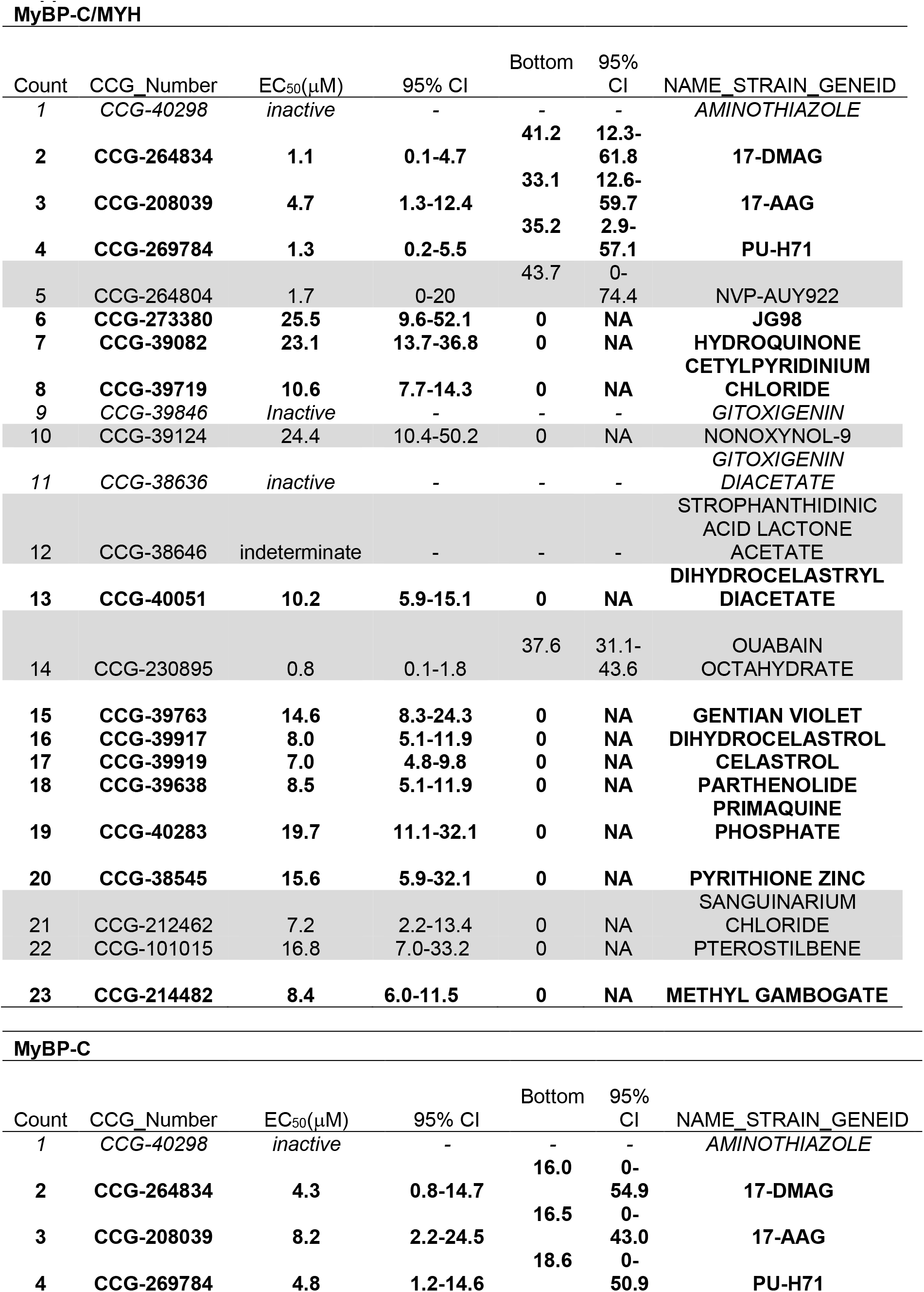
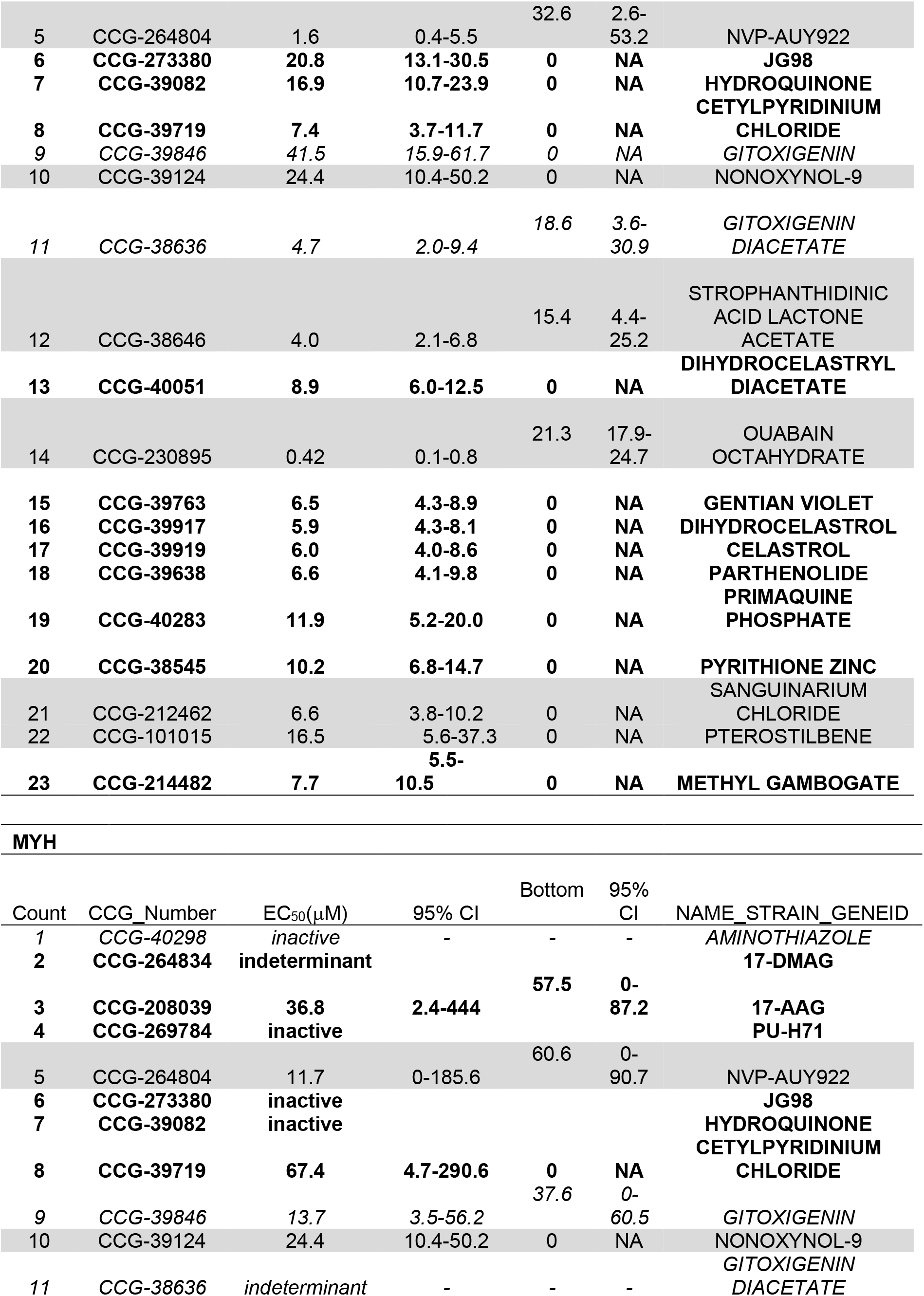
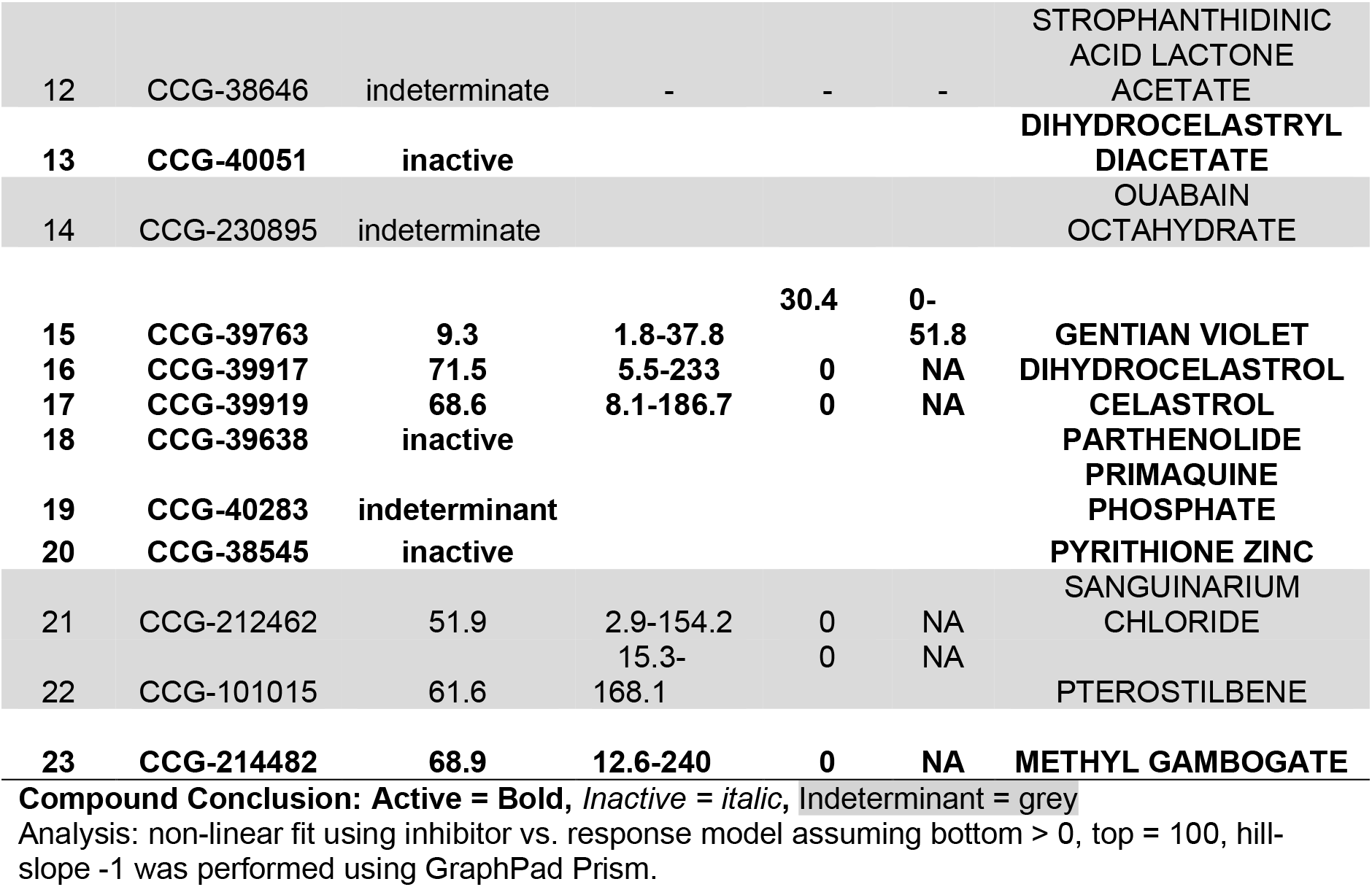
CRC results.

**Supplemental Table 8:**
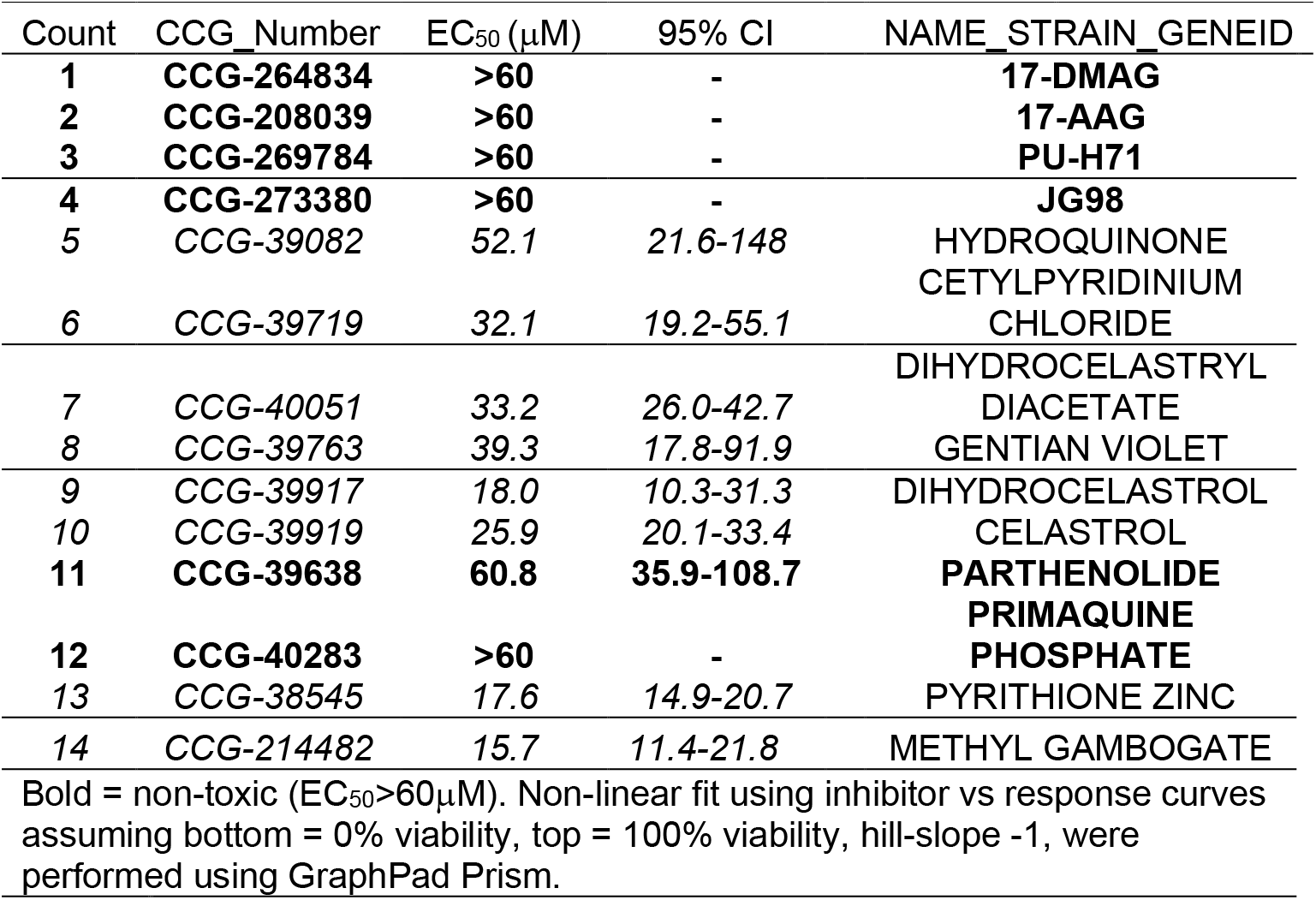
Cell Toxicity Results.

**Supplemental Table 9.**
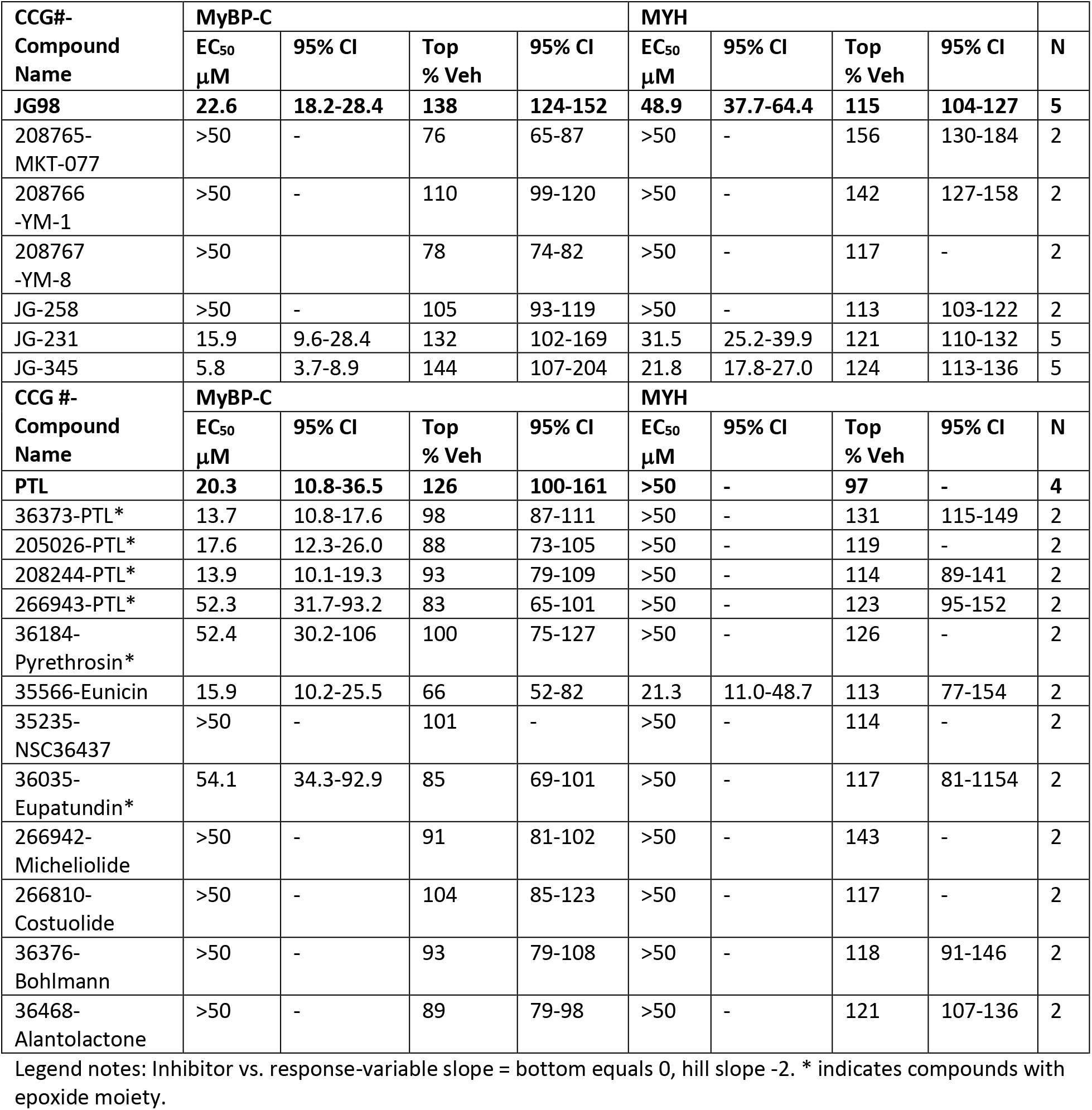
Structure Activity Relationships (SAR) for JG98 and PTL.

**Supplemental Table 10.**
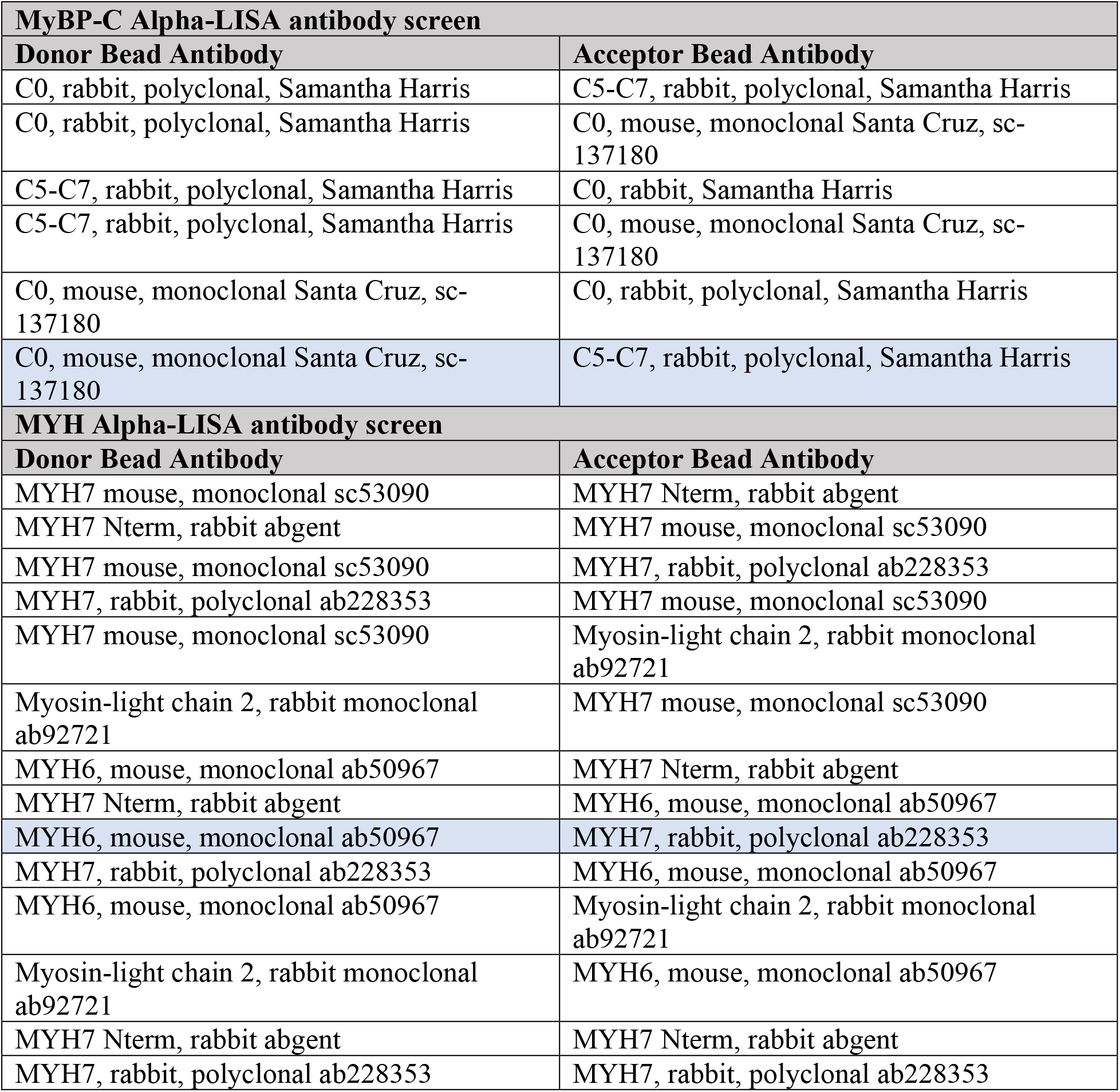
Antibodies tested in the development of MyBP-C and MYH Alpha-LISA assays.

## References

1. Ho CY, Day SM, Ashley EA et al. Genotype and Lifetime Burden of Disease in Hypertrophic Cardiomyopathy: Insights from the Sarcomeric Human Cardiomyopathy Registry (SHaRe). Circulation 2018;138:1387–1398.

2. Walsh R, Thomson KL, Ware JS et al. Reassessment of Mendelian gene pathogenicity using 7,855 cardiomyopathy cases and 60,706 reference samples. Genet Med 2017;19:192–203.

3. Helms AS, Thompson AD, Day SM. Translation of New and Emerging Therapies for Genetic Cardiomyopathies. JACC Basic Transl Sci 2022;7:70–83.

4. Helms AS, Thompson AD, Glazier AA et al. Spatial and Functional Distribution of MYBPC3 Pathogenic Variants and Clinical Outcomes in Patients With Hypertrophic Cardiomyopathy. Circ Genom Precis Med 2020;13:396–405.

5. Vignier N, Schlossarek S, Fraysse B et al. Nonsense-mediated mRNA decay and ubiquitin-proteasome system regulate cardiac myosin-binding protein C mutant levels in cardiomyopathic mice. Circ Res 2009;105:239–48.

6. Glazier AA, Hafeez N, Mellacheruvu D et al. HSC70 is a chaperone for wild-type and mutant cardiac myosin binding protein C. JCI Insight 2018;3.

7. O’Leary TS, Snyder J, Sadayappan S, Day SM, Previs MJ. MYBPC3 truncation mutations enhance actomyosin contractile mechanics in human hypertrophic cardiomyopathy. J Mol Cell Cardiol 2019;127:165–173.

8. Glazier AA, Thompson A, Day SM. Allelic imbalance and haploinsufficiency in MYBPC3-linked hypertrophic cardiomyopathy. Pflugers Arch 2019;471:781–793.

9. Marston S, Copeland O, Jacques A et al. Evidence from human myectomy samples that MYBPC3 mutations cause hypertrophic cardiomyopathy through haploinsufficiency. Circ Res 2009;105:219–22.

10. Barefield D, Kumar M, Gorham J et al. Haploinsufficiency of MYBPC3 exacerbates the development of hypertrophic cardiomyopathy in heterozygous mice. J Mol Cell Cardiol 2015;79:234–43.

11. Monteiro da Rocha A, Guerrero-Serna G, Helms A et al. Deficient cMyBP-C protein expression during cardiomyocyte differentiation underlies human hypertrophic cardiomyopathy cellular phenotypes in disease specific human ES cell derived cardiomyocytes. J Mol Cell Cardiol 2016;99:197–206.

12. Prondzynski M, Kramer E, Laufer SD et al. Evaluation of MYBPC3 trans-Splicing and Gene Replacement as Therapeutic Options in Human iPSC-Derived Cardiomyocytes. Mol Ther Nucleic Acids 2017;7:475–486.

13. Prondzynski M, Mearini G, Carrier L. Gene therapy strategies in the treatment of hypertrophic cardiomyopathy. Pflugers Arch 2019;471:807–815.

14. Mearini G, Stimpel D, Geertz B et al. Mybpc3 gene therapy for neonatal cardiomyopathy enables long-term disease prevention in mice. Nat Commun 2014;5:5515.

15. Helms AS, Tang VT, O’Leary TS et al. Effects of MYBPC3 loss-of-function mutations preceding hypertrophic cardiomyopathy. JCI Insight 2020;5.

16. Dambrot C, Braam SR, Tertoolen LG, Birket M, Atsma DE, Mummery CL. Serum supplemented culture medium masks hypertrophic phenotypes in human pluripotent stem cell derived cardiomyocytes. J Cell Mol Med 2014;18:1509–18.

17. Birket MJ, Ribeiro MC, Kosmidis G et al. Contractile Defect Caused by Mutation in MYBPC3 Revealed under Conditions Optimized for Human PSC-Cardiomyocyte Function. Cell Rep 2015;13:733–745.

18. Chen CY, Salomon AK, Caporizzo MA et al. Depletion of Vasohibin 1 Speeds Contraction and Relaxation in Failing Human Cardiomyocytes. Circ Res 2020;127:e14–e27.

19. Li X, Colvin T, Rauch JN et al. Validation of the Hsp70-Bag3 protein-protein interaction as a potential therapeutic target in cancer. Mol Cancer Ther 2015;14:642–8.

20. Li X, Shao H, Taylor IR, Gestwicki JE. Targeting Allosteric Control Mechanisms in Heat Shock Protein 70 (Hsp70). Curr Top Med Chem 2016;16:2729–40.

21. Freund RRA, Gobrecht P, Fischer D, Arndt HD. Advances in chemistry and bioactivity of parthenolide. Nat Prod Rep 2020;37:541–565.

22. Shao H, Li X, Moses MA et al. Exploration of Benzothiazole Rhodacyanines as Allosteric Inhibitors of Protein-Protein Interactions with Heat Shock Protein 70 (Hsp70). J Med Chem 2018;61:6163–6177.

23. Ouimet CM, Shao H, Rauch JN et al. Protein Cross-Linking Capillary Electrophoresis for Protein-Protein Interaction Analysis. Anal Chem 2016;88:8272–8.

24. Miyata Y, Li X, Lee HF et al. Synthesis and initial evaluation of YM-08, a blood-brain barrier permeable derivative of the heat shock protein 70 (Hsp70) inhibitor MKT-077, which reduces tau levels. ACS Chem Neurosci 2013;4:930–9.

25. Li X, Srinivasan SR, Connarn J et al. Analogs of the Allosteric Heat Shock Protein 70 (Hsp70) Inhibitor, MKT-077, as Anti-Cancer Agents. ACS Med Chem Lett 2013;4.

26. Robison P, Caporizzo MA, Ahmadzadeh H et al. Detyrosinated microtubules buckle and bear load in contracting cardiomyocytes. Science 2016;352:aaf0659.

27. Aillaud C, Bosc C, Peris L et al. Vasohibins/SVBP are tubulin carboxypeptidases (TCPs) that regulate neuron differentiation. Science 2017;358:1448–1453.

28. Nieuwenhuis J, Adamopoulos A, Bleijerveld OB et al. Vasohibins encode tubulin detyrosinating activity. Science 2017;358:1453–1456.

29. Janke C, Bulinski JC. Post-translational regulation of the microtubule cytoskeleton: mechanisms and functions. Nat Rev Mol Cell Biol 2011;12:773–86.

30. Martin TG, Myers VD, Dubey P et al. Cardiomyocyte contractile impairment in heart failure results from reduced BAG3-mediated sarcomeric protein turnover. Nat Commun 2021;12:2942.

31. Kirk JA, Cheung JY, Feldman AM. Therapeutic targeting of BAG3: considering its complexity in cancer and heart disease. J Clin Invest 2021;131.

32. Rousaki A, Miyata Y, Jinwal UK, Dickey CA, Gestwicki JE, Zuiderweg ER. Allosteric drugs: the interaction of antitumor compound MKT-077 with human Hsp70 chaperones. J Mol Biol 2011;411:614–32.

33. Liu L, Sun K, Zhang X, Tang Y, Xu D. Advances in the role and mechanism of BAG3 in dilated cardiomyopathy. Heart Fail Rev 2021;26:183–194.

34. Feldman AM, Begay RL, Knezevic T et al. Decreased levels of BAG3 in a family with a rare variant and in idiopathic dilated cardiomyopathy. J Cell Physiol 2014;229:1697–702.

35. Ellinor PT, Sasse-Klaassen S, Probst S et al. A novel locus for dilated cardiomyopathy, diffuse myocardial fibrosis, and sudden death on chromosome 10q25-26. J Am Coll Cardiol 2006;48:106–11.

36. Villard E, Perret C, Gary F et al. A genome-wide association study identifies two loci associated with heart failure due to dilated cardiomyopathy. Eur Heart J 2011;32:1065–76.

37. Harper AR, Goel A, Grace C et al. Common genetic variants and modifiable risk factors underpin hypertrophic cardiomyopathy susceptibility and expressivity. Nat Genet 2021;53:135–142.

38. Tadros R, Francis C, Xu X et al. Shared genetic pathways contribute to risk of hypertrophic and dilated cardiomyopathies with opposite directions of effect. Nat Genet 2021;53:128–134.

39. Yang J, Grafton F, Ranjbarvaziri S et al. Phenotypic screening with deep learning identifies HDAC6 inhibitors as cardioprotective in a BAG3 mouse model of dilated cardiomyopathy. Sci Transl Med 2022;14:eabl5654.

40. Judge LM, Perez-Bermejo JA, Truong A et al. A BAG3 chaperone complex maintains cardiomyocyte function during proteotoxic stress. JCI Insight 2017;2.

41. Rauch JN, Gestwicki JE. Binding of human nucleotide exchange factors to heat shock protein 70 (Hsp70) generates functionally distinct complexes in vitro. J Biol Chem 2014;289:1402–14.

42. Doong H, Rizzo K, Fang S, Kulpa V, Weissman AM, Kohn EC. CAIR-1/BAG-3 abrogates heat shock protein-70 chaperone complex-mediated protein degradation: accumulation of poly-ubiquitinated Hsp90 client proteins. J Biol Chem 2003;278:28490–500.

43. Seeger T, Shrestha R, Lam CK et al. A Premature Termination Codon Mutation in MYBPC3 Causes Hypertrophic Cardiomyopathy via Chronic Activation of Nonsense-Mediated Decay. Circulation 2019;139:799–811.

44. Martin TG, Delligatti CE, Muntu NA, Stachowski-Doll MJ, Kirk JA. Pharmacological inhibition of BAG3-HSP70 with the proposed cancer therapeutic JG-98 is toxic for cardiomyocytes. J Cell Biochem 2022;123:128–141.

## References

1. Lian X, Zhang J, Azarin SM et al. Directed cardiomyocyte differentiation from human pluripotent stem cells by modulating Wnt/beta-catenin signaling under fully defined conditions. Nat Protoc 2013;8:162–75.

2. Burridge PW, Holmstrom A, Wu JC. Chemically Defined Culture and Cardiomyocyte Differentiation of Human Pluripotent Stem Cells. Curr Protoc Hum Genet 2015;87:21 3 1-21 3 15.

3. Tohyama S, Hattori F, Sano M et al. Distinct metabolic flow enables large-scale purification of mouse and human pluripotent stem cell-derived cardiomyocytes. Cell Stem Cell 2013;12:127–37.

4. Ufford K, Friedline S, Tong Z et al. Myofibrillar Structural Variability Underlies Contractile Function in Stem Cell-Derived Cardiomyocytes. Stem Cell Reports 2021;16:470–477.

5. Glazier AA, Hafeez N, Mellacheruvu D et al. HSC70 is a chaperone for wild-type and mutant cardiac myosin binding protein C. JCI Insight 2018;3.

6. Carrier L, Schlossarek S, Willis MS, Eschenhagen T. The ubiquitin-proteasome system and nonsense-mediated mRNA decay in hypertrophic cardiomyopathy. Cardiovasc Res 2010;85:330–8.

7. Schlossarek S, Schuermann F, Geertz B, Mearini G, Eschenhagen T, Carrier L. Adrenergic stress reveals septal hypertrophy and proteasome impairment in heterozygous Mybpc3-targeted knock-in mice. J Muscle Res Cell Motil 2012;33:5–15.

8. Tanaka A, Yuasa S, Mearini G et al. Endothelin-1 induces myofibrillar disarray and contractile vector variability in hypertrophic cardiomyopathy-induced pluripotent stem cell-derived cardiomyocytes. J Am Heart Assoc 2014;3:e001263.

9. Gray MO, Long CS, Kalinyak JE, Li HT, Karliner JS. Angiotensin II stimulates cardiac myocyte hypertrophy via paracrine release of TGF-beta 1 and endothelin-1 from fibroblasts. Cardiovasc Res 1998;40:352–63.

10. Najafi A, Sequeira V, Helmes M et al. Selective phosphorylation of PKA targets after beta-adrenergic receptor stimulation impairs myofilament function in Mybpc3-targeted HCM mouse model. Cardiovasc Res 2016;110:200–14.

11. Barefield D, Kumar M, Gorham J et al. Haploinsufficiency of MYBPC3 exacerbates the development of hypertrophic cardiomyopathy in heterozygous mice. J Mol Cell Cardiol 2015;79:234–43.

12. Jacques AM, Copeland O, Messer AE et al. Myosin binding protein C phosphorylation in normal, hypertrophic and failing human heart muscle. J Mol Cell Cardiol 2008;45:209–16.

13. Judge LM, Perez-Bermejo JA, Truong A et al. A BAG3 chaperone complex maintains cardiomyocyte function during proteotoxic stress. JCI Insight 2017;2.

14. Chen CY, Salomon AK, Caporizzo MA et al. Depletion of Vasohibin 1 Speeds Contraction and Relaxation in Failing Human Cardiomyocytes. Circ Res 2020;127:e14–e27.

18. Li YM, Casida JE. Cantharidin-binding protein: identification as protein phosphatase 2A. Proc Natl Acad Sci U S A 1992;89:11867–70.

19. Li YM, Mackintosh C, Casida JE. Protein phosphatase 2A and its [3H]cantharidin/[3H]endothall thioanhydride binding site. Inhibitor specificity of cantharidin and ATP analogues. Biochem Pharmacol 1993;46:1435–43.

20. Li X, Srinivasan SR, Connarn J et al. Analogs of the Allosteric Heat Shock Protein 70 (Hsp70) Inhibitor, MKT-077, as Anti-Cancer Agents. ACS Med Chem Lett 2013;4.

21. Jinwal UK, Miyata Y, Koren J, 3rd et al. Chemical manipulation of hsp70 ATPase activity regulates tau stability. J Neurosci 2009;29:12079-88.

22. Taylor IR, Dunyak BM, Komiyama T et al. High-throughput screen for inhibitors of protein-protein interactions in a reconstituted heat shock protein 70 (Hsp70) complex. J Biol Chem 2018;293:4014–4025.

23. Srinivasan SR, Cesa LC, Li X et al. Heat Shock Protein 70 (Hsp70) Suppresses RIP1-Dependent Apoptotic and Necroptotic Cascades. Mol Cancer Res 2018;16:58–68.

24. Henning RH, Brundel B. Proteostasis in cardiac health and disease. Nat Rev Cardiol 2017;14:637–653.

25. Wiersma M, Henning RH, Brundel BJ. Derailed Proteostasis as a Determinant of Cardiac Aging. Can J Cardiol 2016;32:1166 e11–20.

26. Lee WC, Lin KY, Chiu YT et al. Substantial decrease of heat shock protein 90 in ventricular tissues of two sudden-death pigs with hypertrophic cardiomyopathy. FASEB J 1996;10:1198–204.

27. Sharma S, Mishra R, Walker BL et al. Celastrol, an oral heat shock activator, ameliorates multiple animal disease models of cell death. Cell Stress Chaperones 2015;20:185–201.

28. Sydor JR, Normant E, Pien CS et al. Development of 17-allylamino-17-demethoxygeldanamycin hydroquinone hydrochloride (IPI-504), an anti-cancer agent directed against Hsp90. Proc Natl Acad Sci U S A 2006;103:17408–13.

29. Smith V, Sausville EA, Camalier RF, Fiebig HH, Burger AM. Comparison of 17-dimethylaminoethylamino-17-demethoxy-geldanamycin (17DMAG) and 17-allylamino-17-demethoxygeldanamycin (17AAG) in vitro: effects on Hsp90 and client proteins in melanoma models. Cancer Chemother Pharmacol 2005;56:126–37.

30. Schulte TW, Neckers LM. The benzoquinone ansamycin 17-allylamino-17-demethoxygeldanamycin binds to HSP90 and shares important biologic activities with geldanamycin. Cancer Chemother Pharmacol 1998;42:273–9.

31. Egorin MJ, Lagattuta TF, Hamburger DR et al. Pharmacokinetics, tissue distribution, and metabolism of 17-(dimethylaminoethylamino)-17-demethoxygeldanamycin (NSC 707545) in CD2F1 mice and Fischer 344 rats. Cancer Chemother Pharmacol 2002;49:7–19.

32. Cheung KM, Matthews TP, James K et al. The identification, synthesis, protein crystal structure and in vitro biochemical evaluation of a new 3,4-diarylpyrazole class of Hsp90 inhibitors. Bioorg Med Chem Lett 2005;15:3338–43.

33. Dymock BW, Barril X, Brough PA et al. Novel, potent small-molecule inhibitors of the molecular chaperone Hsp90 discovered through structure-based design. J Med Chem 2005;48:4212–5.

34. Sharp SY, Boxall K, Rowlands M et al. In vitro biological characterization of a novel, synthetic diaryl pyrazole resorcinol class of heat shock protein 90 inhibitors. Cancer Res 2007;67:2206–16.

35. Sharma SV, Agatsuma T, Nakano H. Targeting of the protein chaperone, HSP90, by the transformation suppressing agent, radicicol. Oncogene 1998;16:2639-45.

36. Kwon HJ, Yoshida M, Muroya K et al. Morphology of ras-transformed cells becomes apparently normal again with tyrosine kinase inhibitors without a decrease in the ras-GTP complex. J Biochem 1995;118:221–8.

37. Kaiser M, Lamottke B, Mieth M et al. Synergistic action of the novel HSP90 inhibitor NVP-AUY922 with histone deacetylase inhibitors, melphalan, or doxorubicin in multiple myeloma. Eur J Haematol 2010;84:337–44.

38. He H, Zatorska D, Kim J et al. Identification of potent water soluble purine-scaffold inhibitors of the heat shock protein 90. J Med Chem 2006;49:381–90.

39. Makley LN, McMenimen KA, DeVree BT et al. Pharmacological chaperone for alpha-crystallin partially restores transparency in cataract models. Science 2015;350:674–7.

40. Magnaghi P, D’Alessio R, Valsasina B et al. Covalent and allosteric inhibitors of the ATPase VCP/p97 induce cancer cell death. Nat Chem Biol 2013;9:548–56.

41. Anderson DJ, Le Moigne R, Djakovic S et al. Targeting the AAA ATPase p97 as an Approach to Treat Cancer through Disruption of Protein Homeostasis. Cancer Cell 2015;28:653–665.

42. Polucci P, Magnaghi P, Angiolini M et al. Alkylsulfanyl-1,2,4-triazoles, a new class of allosteric valosine containing protein inhibitors. Synthesis and structure-activity relationships. J Med Chem 2013;56:437-50.

43. Zhou HJ, Wang J, Yao B et al. Discovery of a First-in-Class, Potent, Selective, and Orally Bioavailable Inhibitor of the p97 AAA ATPase (CB-5083). J Med Chem 2015;58:9480-97.

44. Brundel BJ, Shiroshita-Takeshita A, Qi X et al. Induction of heat shock response protects the heart against atrial fibrillation. Circ Res 2006;99:1394–402.

45. Koishi M, Yokota S, Mae T et al. The effects of KNK437, a novel inhibitor of heat shock protein synthesis, on the acquisition of thermotolerance in a murine transplantable tumor in vivo. Clin Cancer Res 2001;7:215–9.

46. Yokota S, Kitahara M, Nagata K. Benzylidene lactam compound, KNK437, a novel inhibitor of acquisition of thermotolerance and heat shock protein induction in human colon carcinoma cells. Cancer Res 2000;60:2942–8.

47. Lanka V, Wieland S, Barber J, Cudkowicz M. Arimoclomol: a potential therapy under development for ALS. Expert Opin Investig Drugs 2009;18:1907–18.

48. Ahmed M, Machado PM, Miller A et al. Targeting protein homeostasis in sporadic inclusion body myositis. Sci Transl Med 2016;8:331ra41.

49. Kieran D, Kalmar B, Dick JR, Riddoch-Contreras J, Burnstock G, Greensmith L. Treatment with arimoclomol, a coinducer of heat shock proteins, delays disease progression in ALS mice. Nat Med 2004;10:402–5.

50. Atkinson BN, Woodward HL, Sipthorp J, Fish PV. Regioselective and enantiospecific synthesis of the HSP co-inducer arimoclomol from chiral glycidyl derivatives. Org Biomol Chem 2017;15:9794–9799.

51. Ranek MJ, Zheng H, Huang W et al. Genetically induced moderate inhibition of 20S proteasomes in cardiomyocytes facilitates heart failure in mice during systolic overload. J Mol Cell Cardiol 2015;85:273–81.

52. Doll D, Sarikas A, Krajcik R, Zolk O. Proteomic expression analysis of cardiomyocytes subjected to proteasome inhibition. Biochem Biophys Res Commun 2007;353:436–42.

53. Dong X, Liu J, Zheng H et al. In situ dynamically monitoring the proteolytic function of the ubiquitin-proteasome system in cultured cardiac myocytes. Am J Physiol Heart Circ Physiol 2004;287:H1417–25.

54. Rock KL, Gramm C, Rothstein L et al. Inhibitors of the proteasome block the degradation of most cell proteins and the generation of peptides presented on MHC class I molecules. Cell 1994;78:761–71.

55. Read MA, Neish AS, Luscinskas FW, Palombella VJ, Maniatis T, Collins T. The proteasome pathway is required for cytokine-induced endothelial-leukocyte adhesion molecule expression. Immunity 1995;2:493–506.

56. Jensen TJ, Loo MA, Pind S, Williams DB, Goldberg AL, Riordan JR. Multiple proteolytic systems, including the proteasome, contribute to CFTR processing. Cell 1995;83:129–35.

57. Dick LR, Cruikshank AA, Grenier L, Melandri FD, Nunes SL, Stein RL. Mechanistic studies on the inactivation of the proteasome by lactacystin: a central role for clasto-lactacystin beta-lactone. J Biol Chem 1996;271:7273–6.

58. Fenteany G, Schreiber SL. Lactacystin, proteasome function, and cell fate. J Biol Chem 1998;273:8545–8.

59. Adams J, Kauffman M. Development of the proteasome inhibitor Velcade (Bortezomib). Cancer Invest 2004;22:304–11.

60. Chauhan D, Li G, Podar K et al. The bortezomib/proteasome inhibitor PS-341 and triterpenoid CDDO-Im induce synergistic anti-multiple myeloma (MM) activity and overcome bortezomib resistance. Blood 2004;103:3158–66.

61. Narayana N, Cox S, Shaltiel S, Taylor SS, Xuong N. Crystal structure of a polyhistidine-tagged recombinant catalytic subunit of cAMP-dependent protein kinase complexed with the peptide inhibitor PKI(5-24) and adenosine. Biochemistry 1997;36:4438–48.

62. Cheng HC, Kemp BE, Pearson RB et al. A potent synthetic peptide inhibitor of the cAMP-dependent protein kinase. J Biol Chem 1986;261:989–92.

63. Chen Z, Gopalakrishnan SM, Bui MH et al. 1-Benzyl-3-cetyl-2-methylimidazolium iodide (NH125) induces phosphorylation of eukaryotic elongation factor-2 (eEF2): a cautionary note on the anticancer mechanism of an eEF2 kinase inhibitor. J Biol Chem 2011;286:43951–8.

64. Green EM, Wakimoto H, Anderson RL et al. A small-molecule inhibitor of sarcomere contractility suppresses hypertrophic cardiomyopathy in mice. Science 2016;351:617–21.

65. Rohde JA, Roopnarine O, Thomas DD, Muretta JM. Mavacamten stabilizes an autoinhibited state of two-headed cardiac myosin. Proc Natl Acad Sci U S A 2018;115:E7486–E7494.

66. Heitner SB, Jacoby D, Lester SJ et al. Mavacamten Treatment for Obstructive Hypertrophic Cardiomyopathy: A Clinical Trial. Ann Intern Med 2019;170:741–748.

67. Bogoyevitch MA, Glennon PE, Sugden PH. Endothelin-1, phorbol esters and phenylephrine stimulate MAP kinase activities in ventricular cardiomyocytes. FEBS Lett 1993;317:271–5.

68. Bogoyevitch MA, Glennon PE, Andersson MB et al. Endothelin-1 and fibroblast growth factors stimulate the mitogen-activated protein kinase signaling cascade in cardiac myocytes. The potential role of the cascade in the integration of two signaling pathways leading to myocyte hypertrophy. J Biol Chem 1994;269:1110–9.

69. Takahashi N, Calderone A, Izzo NJ, Jr., Maki TM, Marsh JD, Colucci WS. Hypertrophic stimuli induce transforming growth factor-beta 1 expression in rat ventricular myocytes. J Clin Invest 1994;94:1470–6.

