## Supplemental Information for "An unbiased screen identified the Hsp70-BAG3 complex as a regulator of myosin binding protein C3"

**Outline:**

**Expanded Methods** 3-11

**Expanded Results** 12

**Supplemental Figure 1:** Karyotypes performed by WiCell Institute on newly generated induced pluripotent stem cell (iPSC) lines. **13**

**Supplemental Figure 2:** Immunofluorescence on cardiomyocytes (iPSC-CMs) utilized in high-throughput screen. **14**

**Supplemental Figure 3:** Optimization of MYBP-C Alpha-LISA Assay Conditions. **15**

**Supplemental Figure 4:** Optimization of Myosin Heavy Chain Alpha-LISA Assay Conditions. **16**

**Supplemental Figure 5:** Plate Uniformity. **17**

**Supplemental Figure 6:** Full Screen results. **18**

**Supplemental Figure 7:** Full Concentration Response Curve (CRC) results **19-24**

**Supplemental Figure 8:** Full Cell Toxicity Results **25**

**Supplemental Figure 9:** Inactive Hits from powder retest. **26**

**Supplemental Figure 10:** Western Blot confirmation of PTL and JG-98 Activity **27**

**Supplemental Figure 11:** Genetic Experiments to evaluate if detyrosinated $\alpha$-tubulin (dTyr) regulates MyBP-C protein homeostasis. **28**

**Supplemental Table 1:** Induced pluripotent stem cell (iPSC) lines **29**

**Supplemental Table 2:** Custom Compounds- MyBP-C Homeostasis Literature Review **30**

**Supplemental Table 3:** Compounds for Powder Retest and Structure activity relationships **31-35**

**Supplemental Table 4:** Initial Hits **36-43**

**Supplemental Table 5:** Validated by retesting - MyBP-C/MYH **44-45**

**Supplemental Table 6:** Evaluating MyBP-C and MYH assay results for validated MyBP-C/MYH compounds **46-47**

**Supplemental Table 7:** CRC results **48-50**

**Supplemental Table 8:** Cell Toxicity Results **51**

**Supplemental Table 9:** Structure Activity Relationships (SAR) for JG98 and PTL **52**

**Supplemental Table 10:** Antibodies tested in the development of MyBP-C and MYH Alpha-LISA assays **53**

Where;

$\sigma_p$= standard deviation of positive control (*MYBPC3* Patient1 CMs

$\sigma_n$= standard deviation of negative control (*MYBPC3* (-/-) iPSC-CMs)

$\mu_p$ = mean of positive control

$\mu_n$ = mean of negative control

CHX chase data analysis was performed as previously described(5).

**Expanded Results**

**Validation of initial hits.** For compounds that decreased MyBP-C/MYH we retested initial hits on biological independent cells, tested in triplicate. 52/241 initial hits which decreased MyBP-C/MYH, exhibited a value 3 standard deviations below the positive control in all 3 replicates (Supplemental Table 4). None of initial 70 hits which increased MyBP-C/MYH were able to increase MyBP-C/MYH in all 3 replicates by > 2 stdev (Figure 2). To identify pathways which regulate MyBP-C homeostasis, our ideal hit would decrease MyBP-C protein levels while demonstrating minimal effects on MYH. Thus, during the validation step we also evaluated the MyBP-C and MYH assays separately, using MYH assay as a counter screen. 41/52 compounds decreased MyBP-C > 3 stdev below vehicle control in all 3 replicates. Of these, 23/41 did not increase or decrease MYH by > 3 stdev of control. These 23 compounds were considered our validated hits (Supplemental Table 4), (Figure 2). Next, we re-tested our validated hits on biologically independent *MYBPC3* Patient1 iPSC-CMs at varying concentrations to evaluate concentration response curves (CRCs). Fourteen of these 23 compounds were identified as active (Supplemental Figure 7, Supplemental Table 5), warranting further investigation. Finally, we wanted to ensure we were not selecting compounds with significant cellular toxicity. Thus, CRCs of the 14 active compounds were tested using CellTiter-Fluor$^{TM}$ cell viability assay. 6/14 compounds were identified as non-toxic ($EC_{50}$ > 60 μM) (Supplemental Figure 7, Supplemental Table 7).

**A) *MYBPC3* Patient2 iPSCs**

*Passage#:* 11
*Date of Sample:* 11/7/2018
*Specimen:* Human IPS
*Results:* 46,XX

*Cell Line Sex:* Female

*Reason for Testing:* New Line

*Cell:* 46
*Slide:* G02
*Slide Type:* Karyotype

*Total Counted:* 20
*Total Analyzed:* 8
*Total Karyogrammed:* 4
*Band Resolution:* 425 - 550

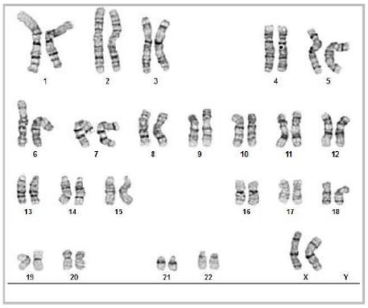

**B) *MYBPC3* Patient3 iPSCs**

*Passage#:* 18
*Date of Sample:* 6/19/2020
*Specimen:* Human IPSC
*Results:* 46,XY

*Cell Line Sex:* Male
*Reason for Testing:* New Line
*Cell:* 8
*Slide:* G02
*Slide Type:* Karyotype

*Total Counted:* 20
*Total Analyzed:* 8
*Total Karyogrammed:* 4
*Band Resolution:* 425 - 525

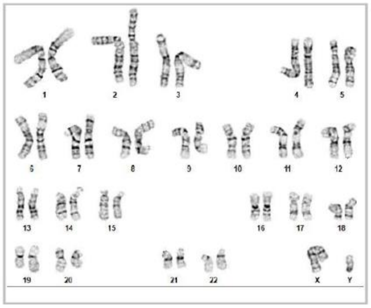

**Supplemental Figure 1: Karyotypes performed by WiCell Institute on newly generated induced pluripotent stem cell (iPSC) lines.** A) *MYBPC3* Patient2 and B) *MYBPC3* Patient3 (Supplemental Table 1).

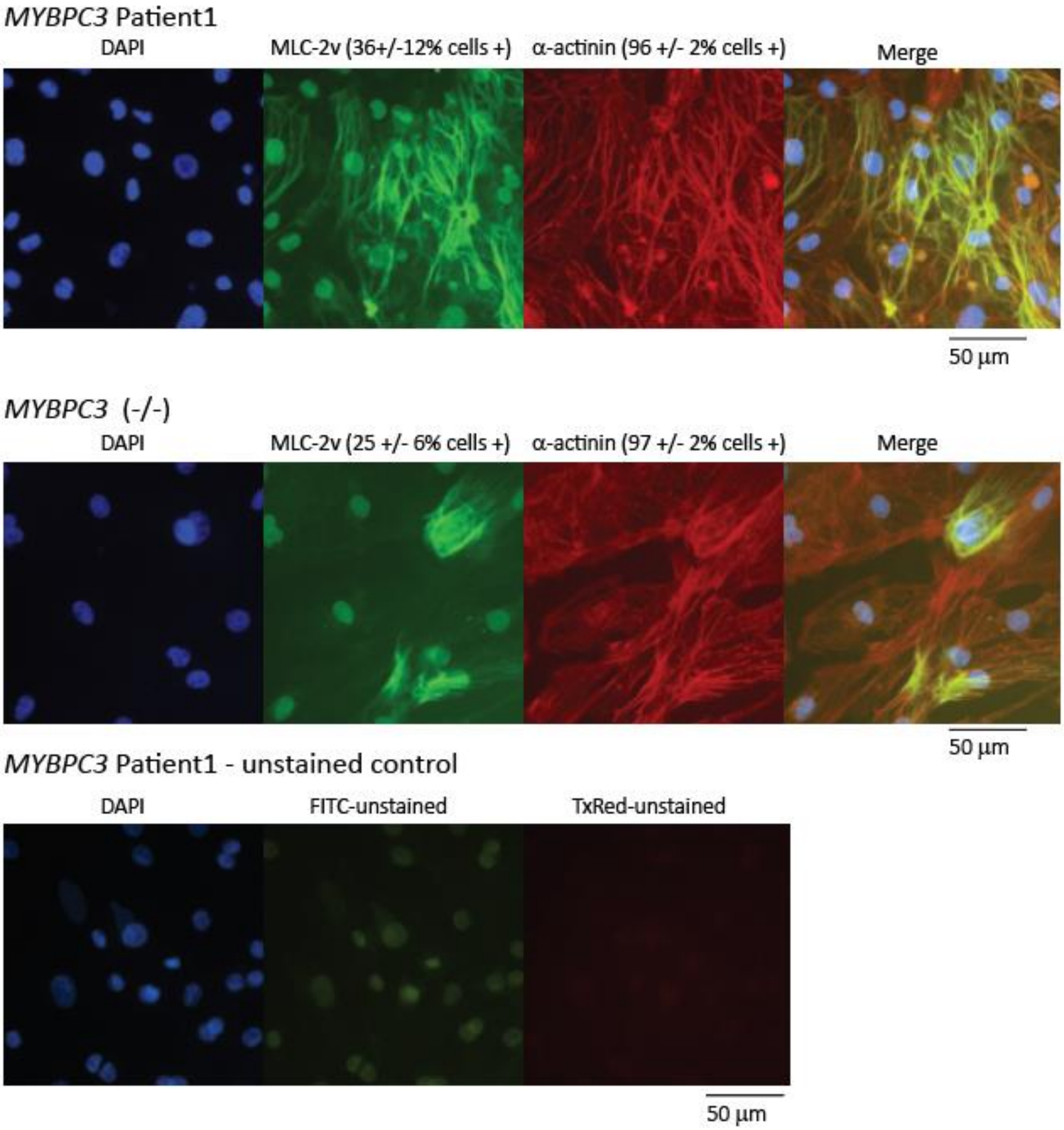


**Supplemental Figure 2: Immunofluorescence on cardiomyocytes (iPSC-CMs) utilized in high-throughput screen.** The screen was performed using *MYBPC3* Patient1 iPSC-CMs. Further, *MYBPC3* (-/-) iPSC-CMs were utilized as a negative control (Supplemental Table 1). Both cell lines demonstrated > 90% purity (> 90% of cells positive for cardiac marker α-actinin, red). Cells were also stained for MLC-2v (green). To evaluate for auto-fluorescence, *MYBPC3* Patient1 cells were imaged after undergoing protocol lacking exposure to primary antibodies.

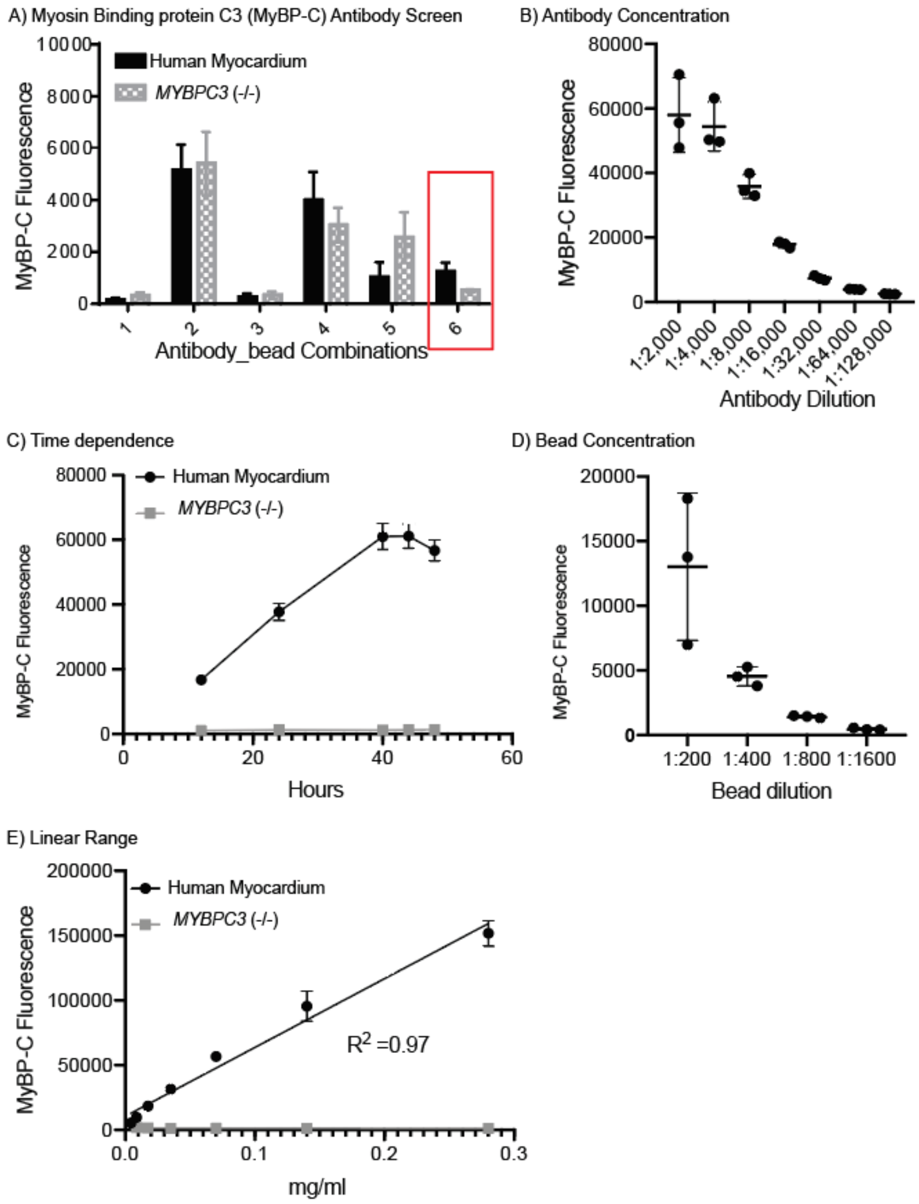


**Supplemental Figure 3: Optimization of MYBP-C Alpha-LISA Assay Conditions.** A) Screening various antibody combinations for detection of MyBP-C within human myectomy tissue and *MYBPC3* (-/-) iPSC-CMS (negative control known to lack expression of MyBP-C). Both samples were tested at a total protein concentration 0.3 mg/ml. Antibody combination with best signal to noise ratio was selected for further optimization (red box). B) The relationship of antibody titer and MyBP-C fluorescence was evaluated. C) Time dependence of MyBP-C Fluorescence was tested. D) The relationship of bead concentration and MyBP-C fluorescence was evaluated. E) The relationship of total protein concentration in MyBP-C Fluorescence was evaluated in human myocardial tissue. *MYBPC3* (-/-) iPSC-CMs was evaluated as a negative control. Average and standard error of mean for samples tested in triplicate are shown in each graph.

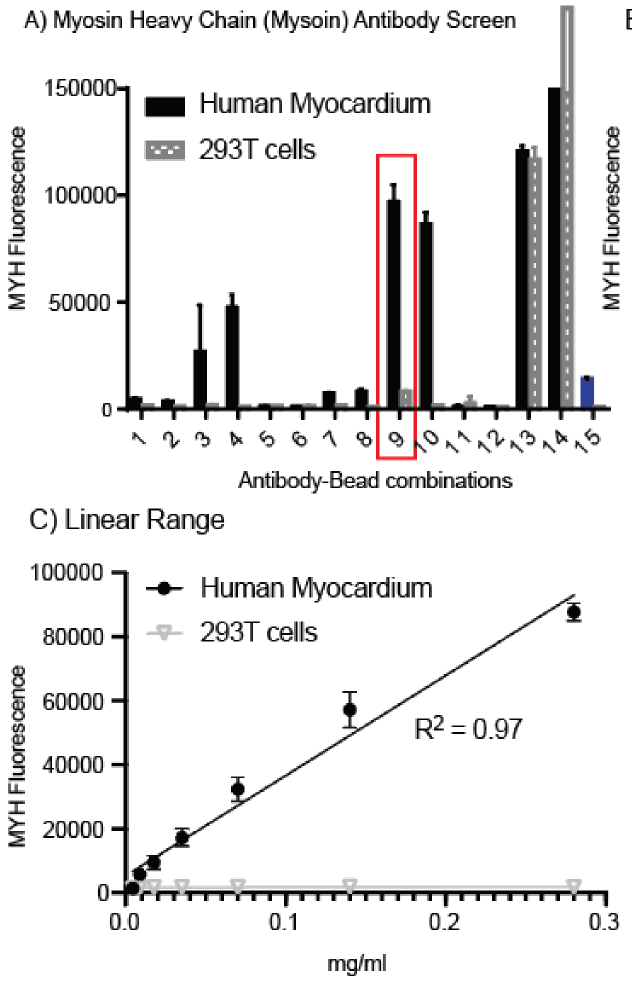


**Supplemental Figure 4: Optimization of Myosin Heavy Chain AlphaLISA Assay Conditions.** A) Screening various antibody combinations for detection of myosin heavy chain (MYH), including antibodies for both αMYH and βMYH7, were tested within human myectomy tissue and 293T cells (negative control known to lack aMYH and bMYH expression). Both samples were tested at a total protein concentration 0.3 mg/ml. Antibody combination 9 was selected (red box) and includes antibodies targeting both MYH6 and MYH7. As a positive control optimized MyBP-C antibody and bead combination was tested in 15 (blue bar graph). B) Time dependence of MYH Fluorescence was tested. C) The relationship of total protein concentration in MYH Fluorescence was evaluated in human myocardial tissue and *MYBPC3* (-/-) iPSC-CMs was evaluated. Average and standard error of mean for samples tested in triplicate are shown in each graph.

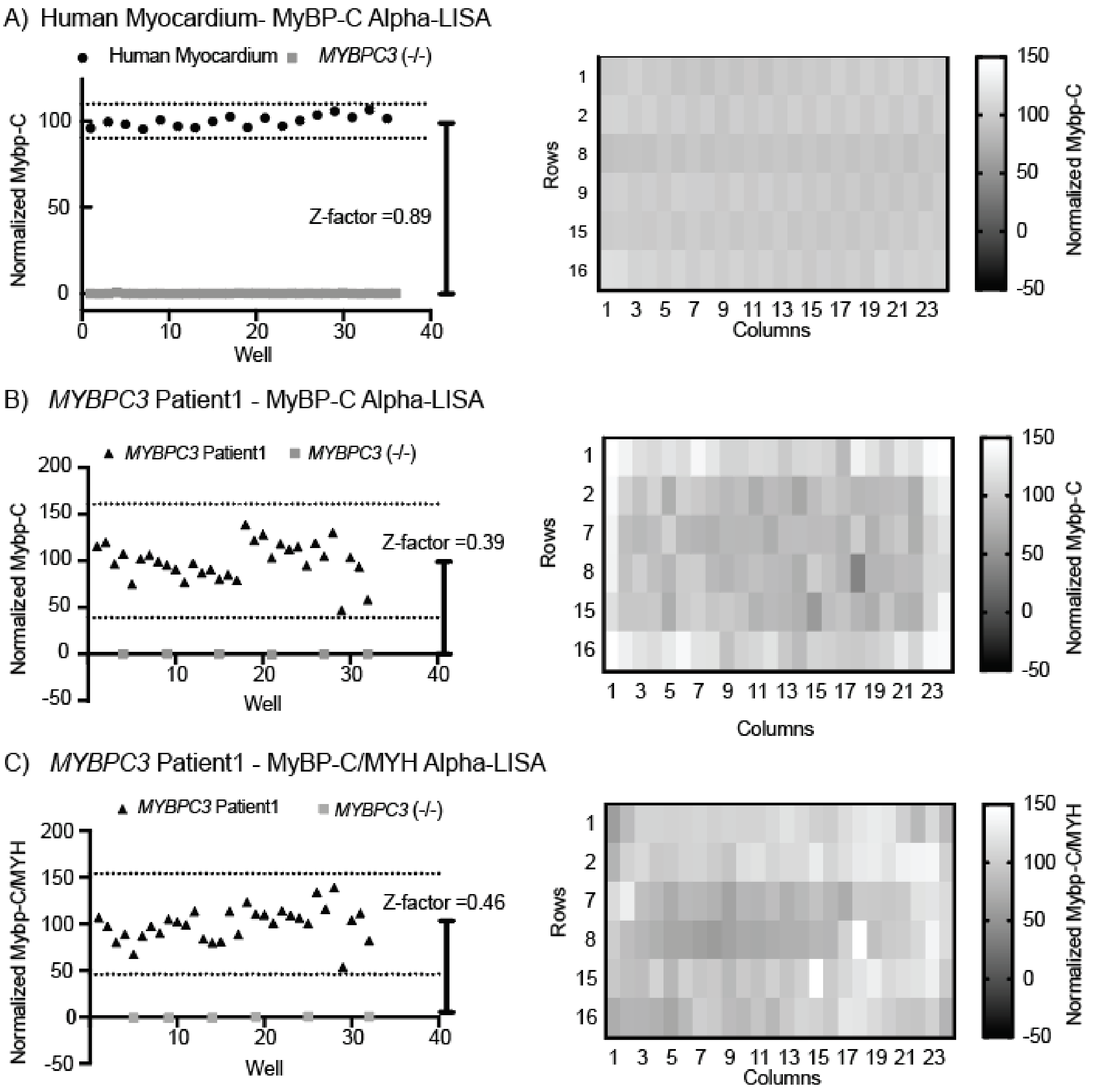


**Supplemental Figure 5: Plate Uniformity.** For (A) Human myocardium- MyBP-C Alpha-LISA and (B) *MYBPC3* Patient1 iPSC-CMs grown in 384 well plates an MyBP-C Alpha-LISA plate uniformity test was performed as shown on the right. Further, a Z-factor analysis was performed using *MYBPC3* (-/-) as a negative control (0%) as shown on the left. Culturing *MYBPC3* Patient1 iPSC-CMs does result in increased variability within the MyBP-C Alpha-LISA assay with reduced signal observed within outer edges of the plate. This is improved when MyBP-C Alpha-LISA assay and MYH Alpha-LISA assay are performed in parallel on cellular lysate from *MYBPC3* patient 1 iPSC-CMs grown in 384 well plates using MyBP-C/MYH as the primary assay endpoint (C).

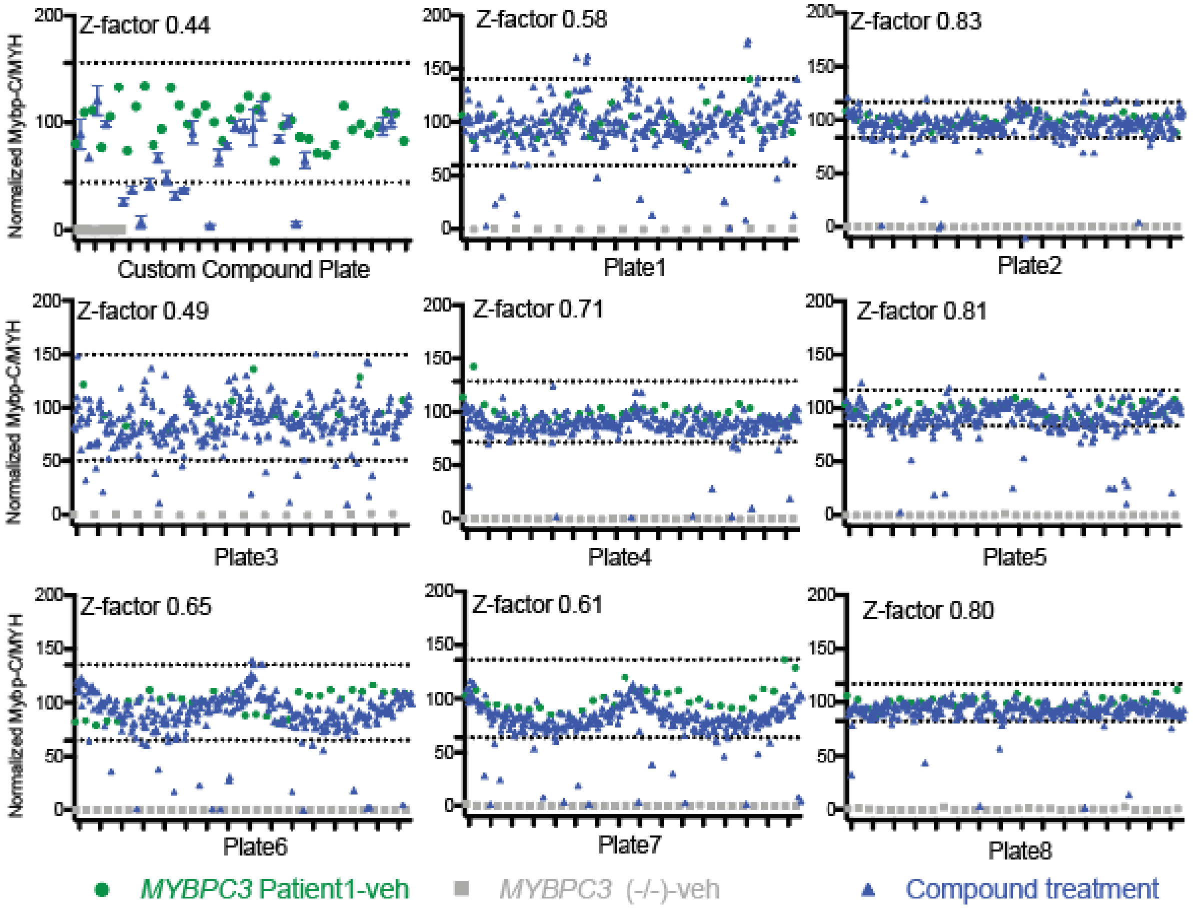


**Supplemental Figure 6: Full Screen results.** Full screen results for every plate tested is shown. Results are normalized with positive control set at 100%- *MYBPC3* Patient1 -veh (iPSC-CMs treated with vehicle control dmso), shown in green circles. The negative controls are set at 0%- *MYBPC3* (-/-)-veh, shown in grey squares. *MYBPC3* Patient1 iPSC-CMs treated with compound are shown in blue triangles. A Z-factor is calculated for each plate and hits are defined as 3 standard deviations above or below the mean of the positive control. This cut-off is marked by a dotted line for each plate.

**Actives**

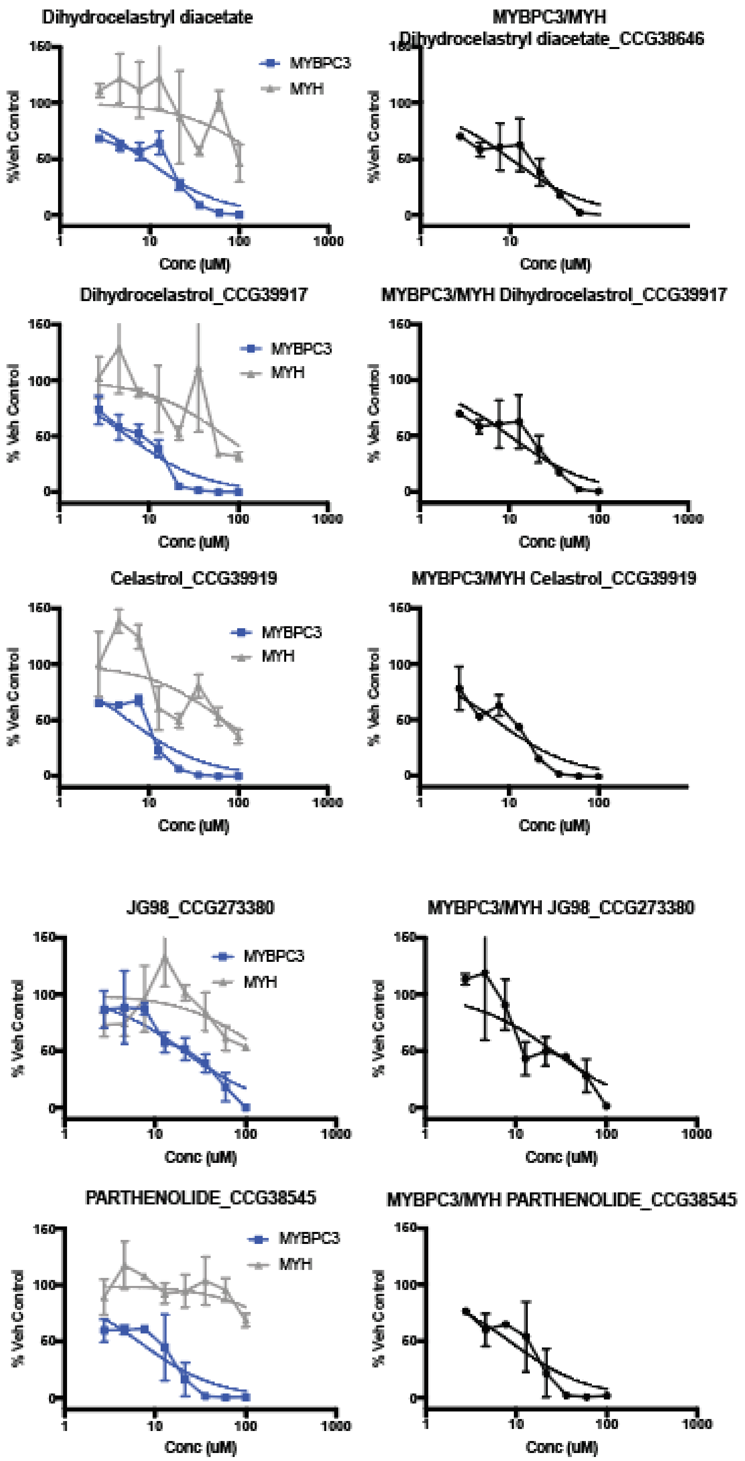
Dihydrocelastryl diacetate
MYBPC3
MYH
%Veh Control
Conc (uM)
MYBPC3/MYH
Dihydrocelastryl diacetate_CCG38646
Dihydrocelastrol_CCG39917
MYBPC3/MYH Dihydrocelastrol_CCG39917
% Veh Control
Celastrol_CCG39919
MYBPC3/MYH Celastrol_CCG39919
JG98_CCG273380
MYBPC3/MYH JG98_CCG273380
PARTHENOLIDE_CCG38545
MYBPC3/MYH PARTHENOLIDE_CCG38545

### Actives Continued

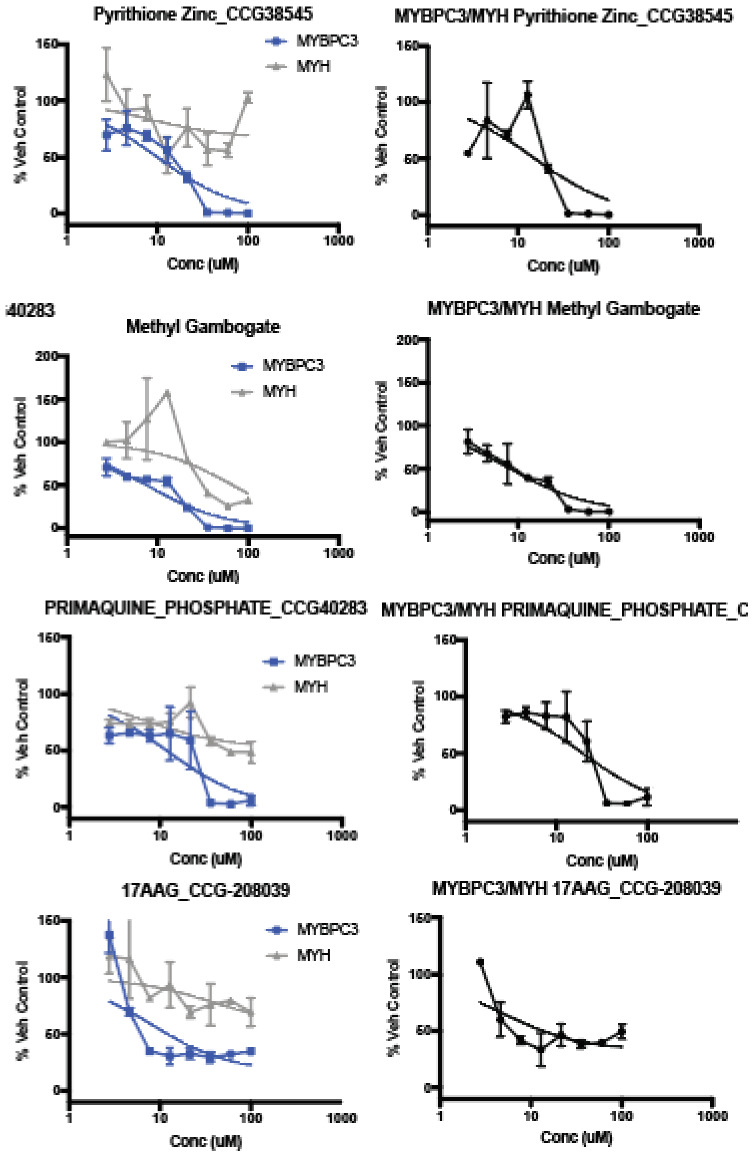

**Actives Continued**

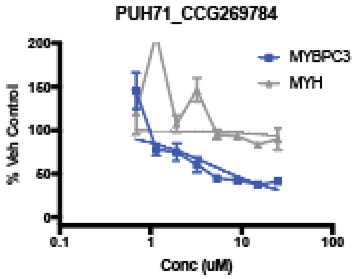


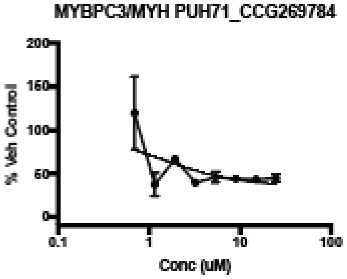


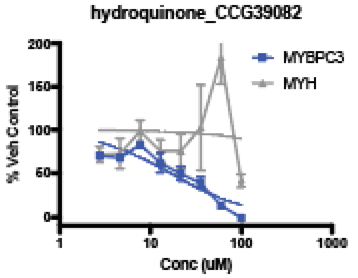


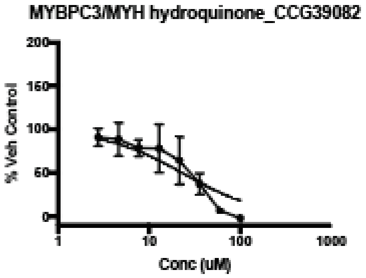


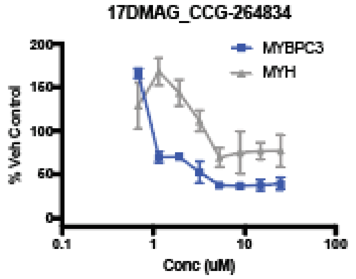


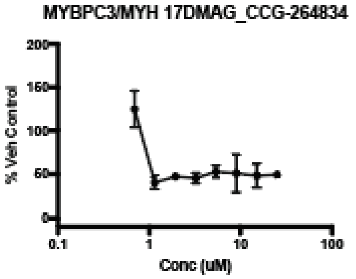


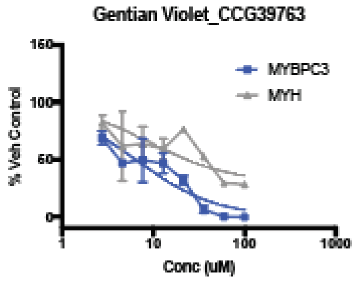


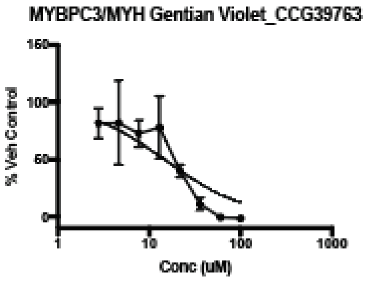


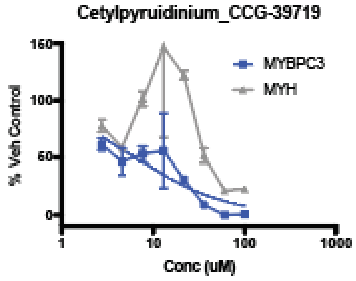


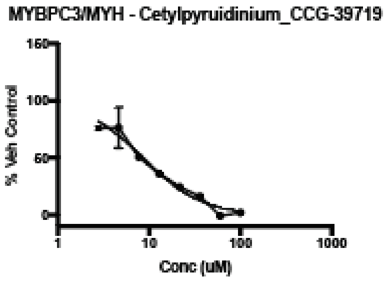

**Inactive**

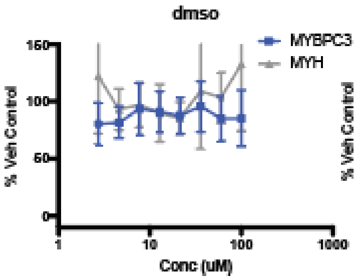
dmso
MYBPC3
MYH
% Veh Control
Conc (uM)

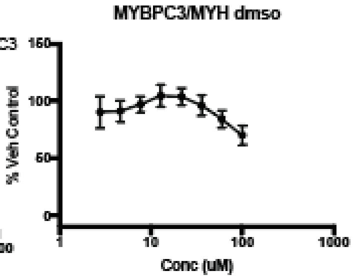
MYBPC3/MYH dmso
% Veh Control
Conc (uM)

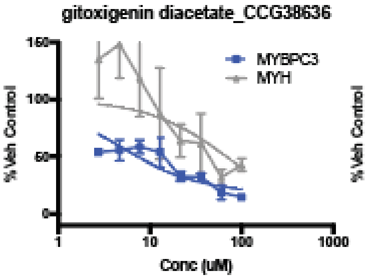
gitoxigenin diacetate_CCG38636
MYBPC3
MYH
% Veh Control
Conc (uM)

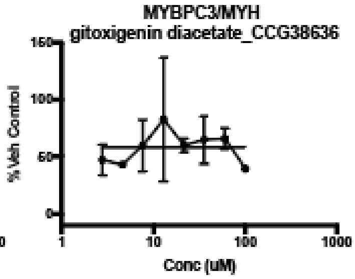
MYBPC3/MYH
gitoxigenin diacetate_CCG38636
% Veh Control
Conc (uM)

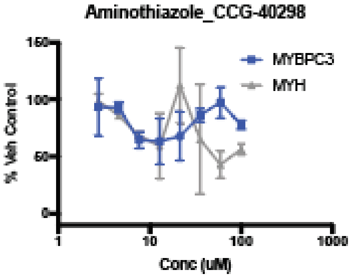
Aminothiazole_CCG-40298
MYBPC3
MYH
% Veh Control
Conc (uM)

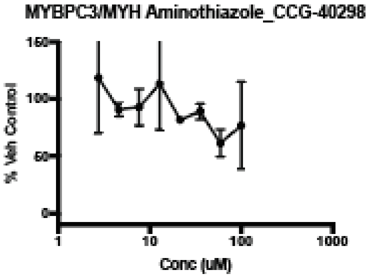
MYBPC3/MYH Aminothiazole_CCG-40298
% Veh Control
Conc (uM)

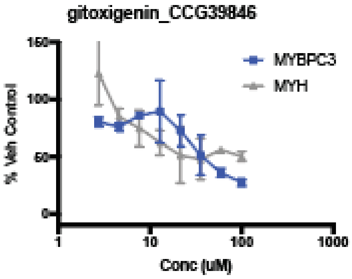
gitoxigenin_CCG39846
MYBPC3
MYH
% Veh Control
Conc (uM)

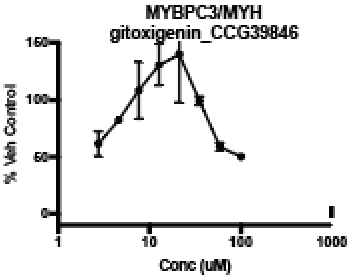
MYBPC3/MYH
gitoxigenin_CCG39846
% Veh Control
Conc (uM)

**Indeterminant**

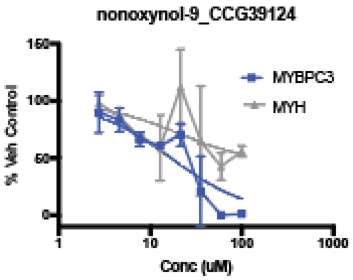


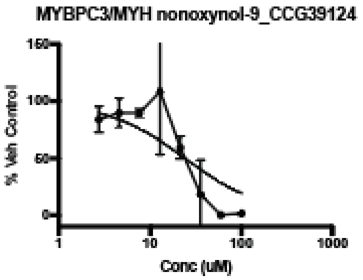


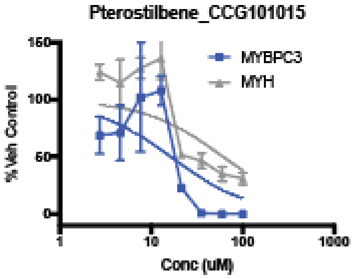


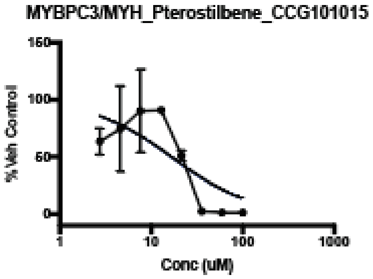


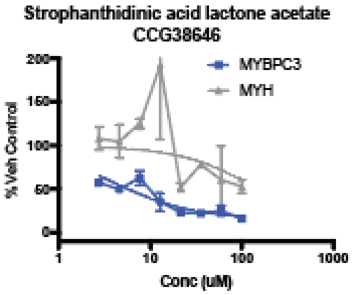


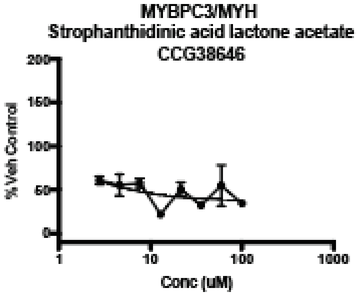


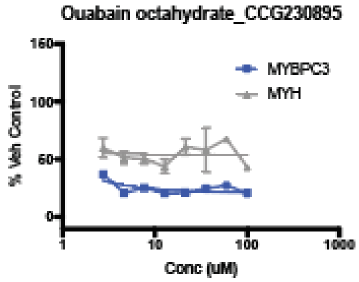


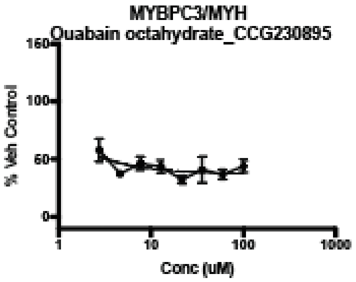


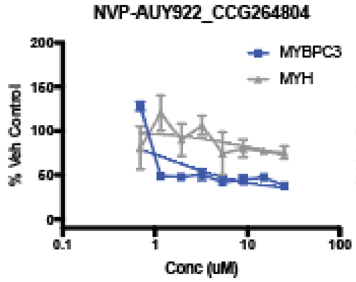


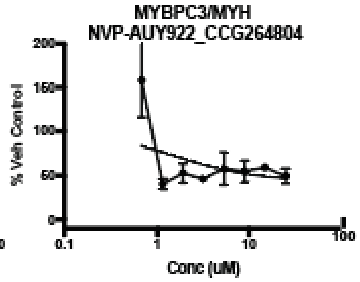

**Indeterminant Continued**

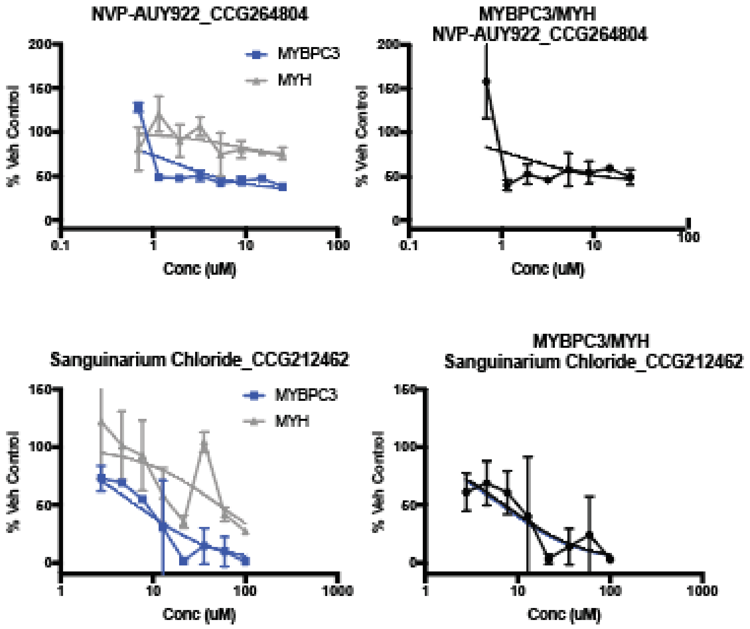


**Supplemental Figure 7: Full Concentration Response Curve (CRC) results-** *MYBPC3* Patient1 iPSC-CMs pooled from 3 independent differentiates were treated with compounds at the indicated concentrations (µM) tested in duplicate for 24 hours. MyBP-C and MYH Alpha-LISA was performed. Results were normalized to 100% *MYBPC3* Patient1 treated with vehicle control (dmso). 0% is set by the negative control. For MyBP-C and MyBP-C/MYH this was *MYBPC3* (-/-) treated with vehicle control (dmso). For MYH 0% is defined by a negative control of 293T cells treated with vehicle control (dmso). The mean and standard error of the mean for MyBP-C, MYH and the ratio of MyBP-C/MYH is shown for each compound. CRC results are shown for each of the 23 validated hit (Decreases MyBP-C/MYH, Decreases MyBP-C, does not decrease MYH). 14 compounds were identified as active. Concentration response curves were fit using GraphPad prism inhibitor vs response curves assuming bottom is greater than 0, top = 100, and hill slope -1. Results are further summarized in Supplemental Table 7.

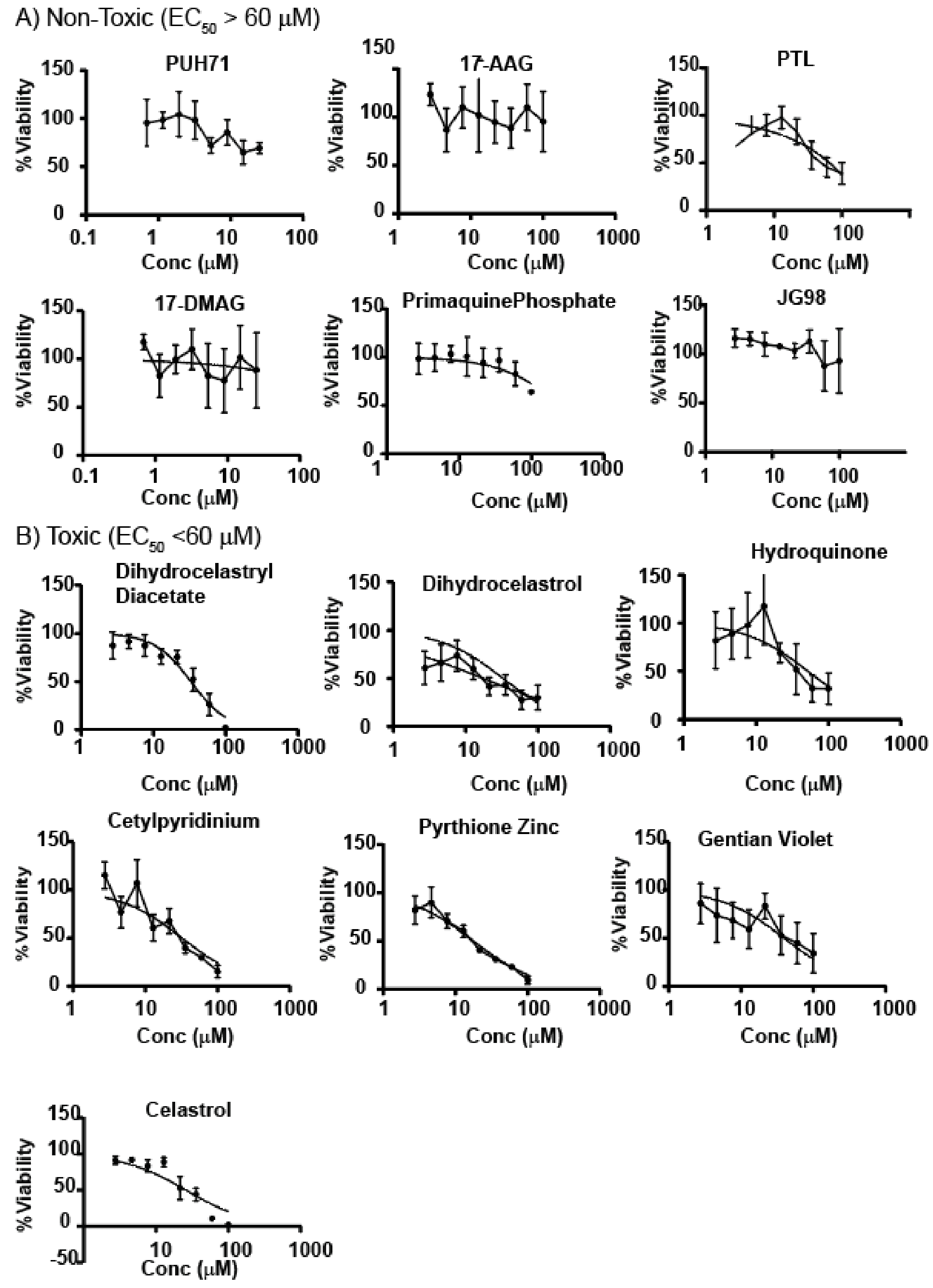


**Supplemental Figure 8: Full Cell Toxicity Results-** Tested in duplication on 2 biologic replicates of *MYBPC3* Patient1 iPSC-CMs (N=4). Results are normalized to %Viability, where 100% = *MYBPC3* Patient1 iPSC-CMs treated with vehicle control (dmso) and 0% = buffer only control (no live cells). Average and standard error of mean shown and fitted using GraphPad Prism inhibitor vs response non-linear fit assuming Top = 100, Bottom = 0 and hill-slope = -1. A) Non-toxic compounds have $EC_{50}$ of > 60 μM. B) Toxic compounds have $EC_{50}$ of <60 μM.

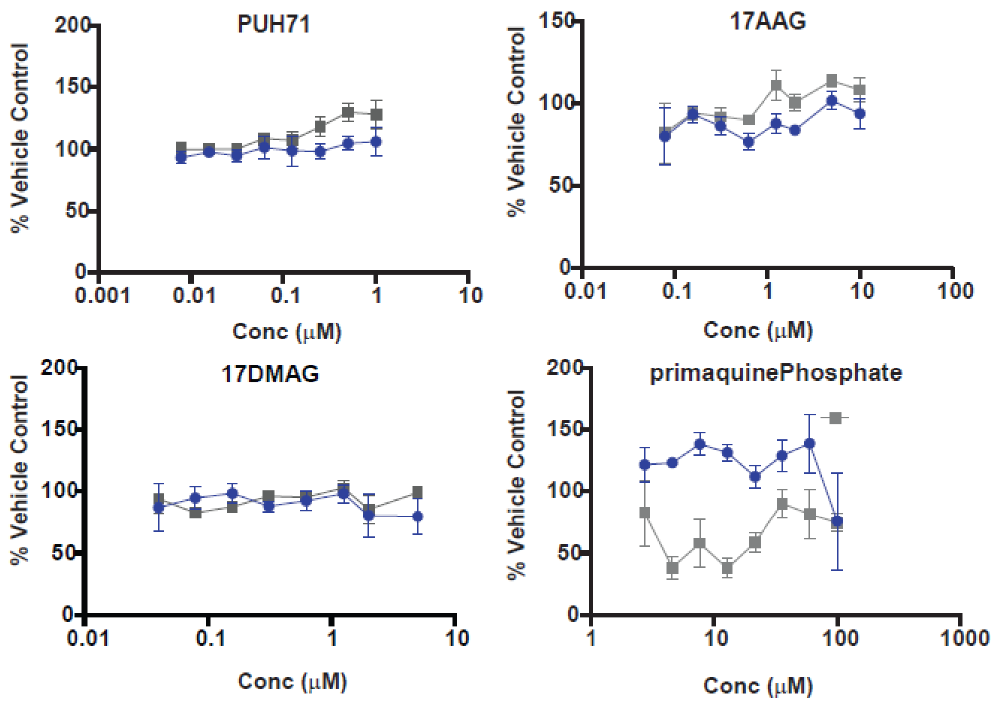


**Supplemental Figure 9: Inactive Hits from powder retest.** 4/6 compounds lost activity observed during screening when fresh powder was tested (PUH71, 17-AAG, 17-DMAG, primaquine phosphate). The results of compound that were inactive during powder retest are shown above. All compounds were tested in duplicate on independently differentiated and pooled MYBPC3 Patient1 iPSC-CMs (n =4). Average and standard error of mean is depicted for each concentration tested. MyBP-C Alpha-LISA results are shown in blue circles and MYH Alpha-LISA result are shown in grey squares. Of note, the Hsp90 inhibitors did not exhibit typical dose dependent activity in CRC performed, but rather a rapid decrease in MyBP-C at lower concentrations (Supplemental Figure 7). As such a lower maximum concentration based on their known Hsp90 inhibitor $EC_{50}$ was selected for their powder retest.

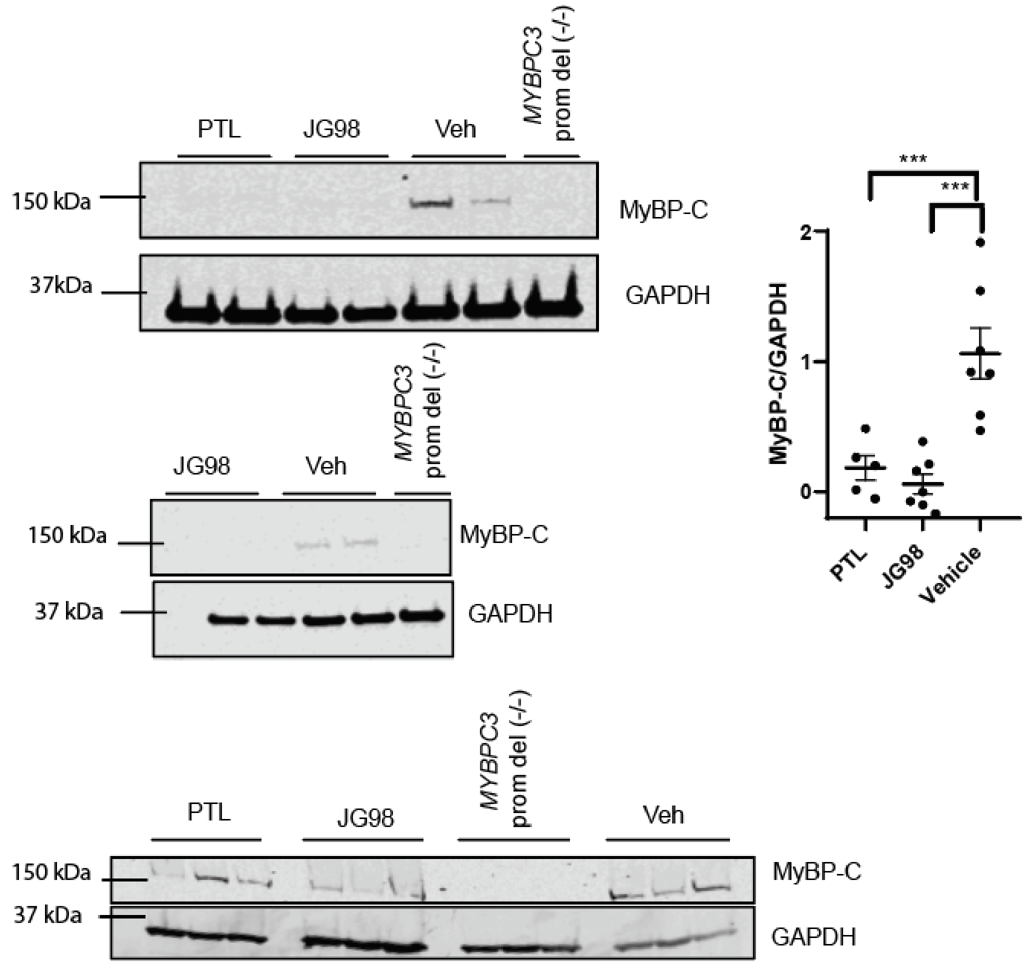


**Supplemental Figure 10: Western Blot confirmation of PTL and JG-98 Activity.** Western blot using Mouse monoclonal *MYBPC3* (Santa Cruz sc-137180) on denaturing SDS-page gel were performed on cellular lysate from Ctrl-1 iPSC-CMs treated with JG-98, PTL at 20 mM or vehicle control are shown. was utilized. MyBP-C levels are normalized using GAPDH loading control. In all graphs, mean and standard error of the mean are shown, *** indicates p-value <0.0001.

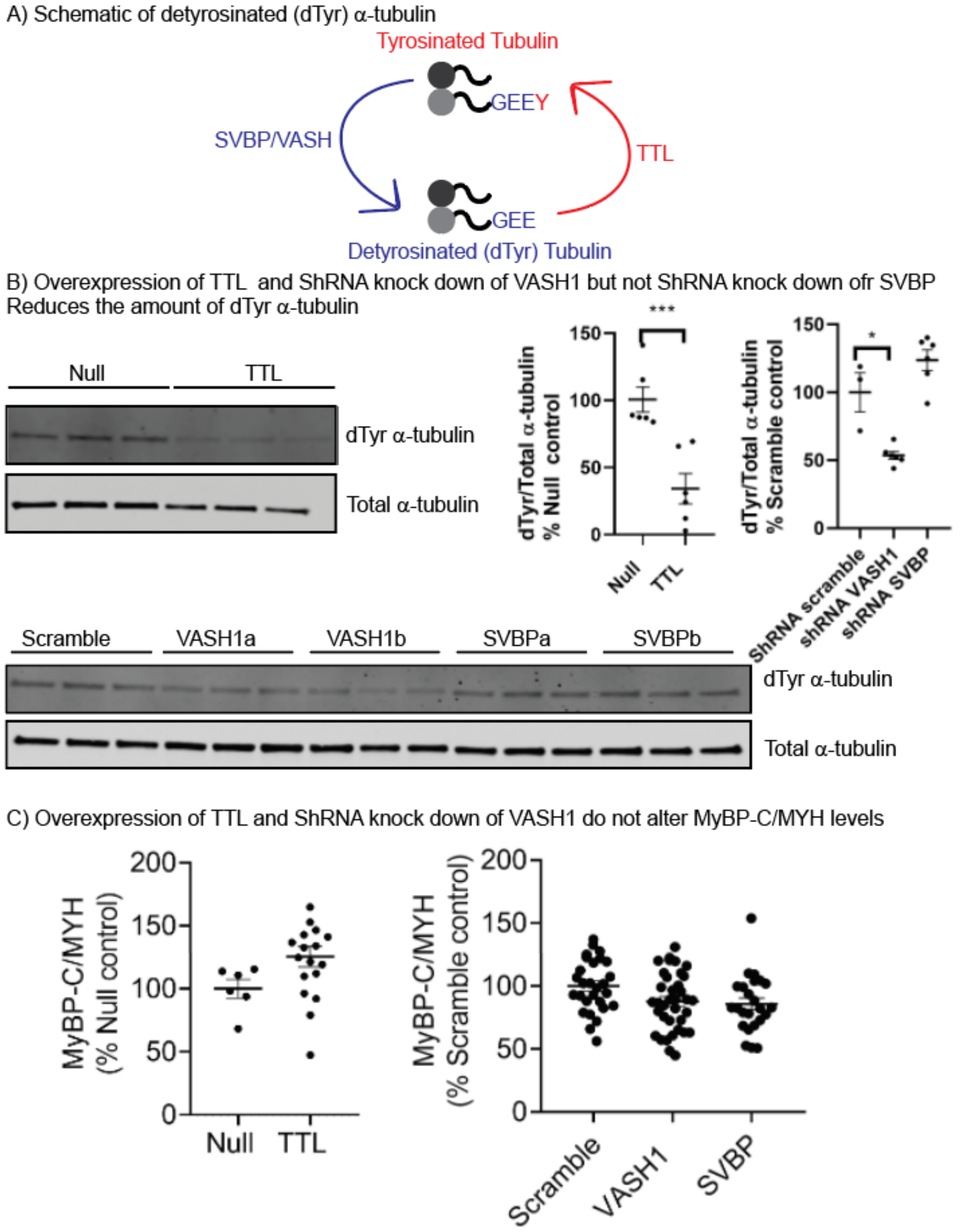


**Supplemental Figure 11: Genetic Experiments to evaluate if detyrosinated α-tubulin (dTyr) regulates MyBP-C protein homeostasis**. (A) The proteins that drive tyrosination and detyrosination of α-tubulin are depicted. B) *MYBPC3* Patient3 iPSC-CMs were treated with adenovirus MOI 10 carrying a null control, the enzyme TTL, or shRNA (Scrambled, or targeting- VASH1, SVBP). Two shRNA constructs were used for both VASH1 and SVBP (a,b).(14) Treated cells were lysed and analyzed by (B) western blot to evaluate their effect on dTyr-α-tubulin or (C) Alpha-LISA to evaluate their effect on MyBP-C/MYH protein levels. Graphs show mean and standard error of the mean, p-value is inducated by; * <0.05, ** < 0.005, *** <0.0005.

**Supplemental Table 1: Induced pluripotent stem cell (iPSC lines)**

| Name | Original Source | *MYBPC3* genotype | Gender | Previously Published |
|---|---|---|---|---|
| Ctrl-1 | WiCell Repository Madison WI (Helms Lab) | +/+ | Male | (15) |
| Ctrl-2 | Ncyte, Ncardia The Netherlands | +/+ | Male | |
| *MYBPC3* knock-out (*MYBPC3* -/-) | Gene Edited (Helms Lab) | -/- | Male | (15) |
| *MYBPC3* Patient1 | Patient Derived (Mummery Lab) | +/c.2373dupG | Male | (15-17) |
| *MYBPC3* Patient2 | Newly generated, (Helms Lab) | +/ c.506-1G>A | Female | |
| *MYBPC3* Patient3 | Newly generated | +/ c.2373dupG | Male | |

### Supplemental Table 2: Custom Compounds- MyBP-C Homeostasis Literature Review

| (18,19)Compound Target | Compound Name | Source | MW g/mol | EC50 μM | Ref | Molecular Formula | CAS Number |
|---|---|---|---|---|---|---|---|
| Hsp70(5,15) | YM-1 | Gestwicki | 418.0 | 2.4 | (5,20) | $C_{20}H_{20}ClN_3OS_2$ | 409086-68-6 |
| | 115-7c | Gestwicki | 477.3 | 120 | (21) | $C_{23}H_{22}Cl_2N_2O_5$ | 908074-72-6 |
| | MKT-077 | Gestwicki | 433.0 | 2 | (20) | $C_{21}H_{23}ClN_3OS_2$ | 147366-41-4 |
| | IT2-144 | Gestwicki | 400.1 | 6.9 | (22) | | |
| | JG-98 | Gestwicki | 534.5 | 0.12 | (23) | $C_{24}H_{21}Cl_2N_3OS_3$ | 1456551-16-8 |
| Hsp90*(24-27) | 17-AAG | Santa Cruz | 585.7 | 0.12 | (28,29) | $C_{31}H_{43}N_3O_8$ | 75747-14-7 |
| | Geldanamycin | Santa Cruz | 560.6 | 0.20 | (30) | $C_{29}H_{40}N_2O_9$ | 30562-34-6 |
| | 17-DMAG | Cayman | 653.2 | 0.024 | (29,31) | $C_{32}H_{49}ClN_4O_8$ | 467214-21-7 |
| | CCT018159 | Tocris | 374.9 | 5.7 | (32-34) | $C_{20}H_{20}N_2O_4$ | 171009-07-7 |
| | radicicol | Cayman | 364.7 | 0.27 | (35,36) | $C_{18}H_{17}ClO_6$ | 12772-57-5 |
| | NVP-AUY922 | Cayman | 465.5 | 0.02 | (37) | $C_{26}H_{31}N_3O_5$ | 747412-49-3 |
| | PU-H71 | Axon Medchem | 621.6 | 0.05 | (38) | $C_{18}H_{21}IN_6O_2S$ | 973436-91 |
| smHsp(5) | Compound 28 | Geswticki | 402.7 | >10 | (39) | $C_{27}H_{46}O_2$ | PUBCHEM CID: 6710665 |
| VCP/p97(5) | NMS-873 | Tocris | 520.7 | >10 | (40-42) | $C_{27}H_{28}N_4O_3S_2$ | 1418013-75-8 |
| | CB5083 | Cayman | 413.5 | 0.011 | (42,43) | $C_{24}H_{23}N_5O_2$ | 1542705-92-9 |
| **Heat Shock activators(15,44)** | KNK437 | Sigma | 245.2 | 95 | (45,46) | $C_{13}H_{11}NO_4$ | 218924-25-5 |
| | Arimoclomol | Carbosynth | | NR | (47-50) | $C_{14}H_{20}ClN_3O_3$ | 289893-25-0 |
| **Proteasome(5,51)** | MG132 | Sigma | 475.6 | 0.004 | (52-56) | $C_{26}H_{41}N_3O_5$ | 133407-82-6 |
| | Lactacystin | Sigma | 376.4 | 0.60 | (57,58) | $C_{15}H_{24}N_2O_7S$ | 133343-34-7 |
| | Bortezomib | Sigma | 384.2 | 0.05 | (59,60) | $C_{19}H_{25}BN_4O_4$ | 179324-69-7 |
| ***MYBPC3* phos(12)** | | | | | | | |
| PKA-I | PKI (5-24) | Cayman | 2,222 | 0.002 | (61,62) | $C_{94}H_{148}N_{32}O_{31}$ | 99534-03-9 |
| CAMKII-I | A484954 | Sigma | 289.3 | 0.28 | (63) | $C_{13}H_{15}N_5O_3$ | 142557-61-7 |
| PP1 and PP2A-I | Cantharidin | Cayman | 196.2 | 1.00 | | $C_{10}H_{12}O_4$ | 56-25-7 |
| **HCM therapies** | | | | | | | |
| Myosin | Mavacamten | Cayman | 273.3 | 0.3 | (64-66) | $C_{15}H_{19}N_3O_2$ | 1642288-47-8 |
| **Hypertrophic(8,9) agonists** | | | | | | | |
| | Endothelin1 | Sigma | 2491.9 | 0.02 | (67,68) | | |
| | TGF-beta | Sigma | | 0.5ng/ml | (9,69) | | |

Compounds were tested at 20 μM if $EC_{50}$ < 20 and > 0.1 μM, at 200 μM if $EC_{50}$ > 20 μM and 2 μM if $EC_{50}$ <0.1 μM. Well established concentrations for testing in cardiomyocytes 20nM and 0.5 ng/ml were utilized for Endothelin-1 and TGF-beta respectively. Hsp70 and smHsp compounds were provided by Jason Gestwicki's laboratory.

### Supplemental Table 3: Compounds for Powder Retest and Structure activity relationships

| Compounds for Powder Retest | | | | | |
|---|---|---|---|---|---|
| **CCG#** | **Compound Name** | CAS# | Source | Pubchem CID | Pubchem Structure |
| **40283** | **Primaquine Phosphate** | 63-45-6 | Microsource #1500500 | 6135 | 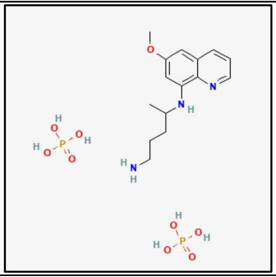 |
| **39638** | **Parthenolide (PTL)** | 20554-84-1 | Microsource #1503640 | 5702252 | 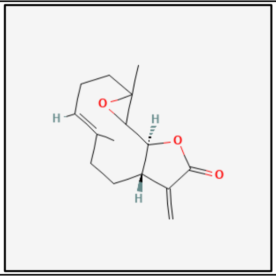 |
| 208039 | **17-AAG** | 75747-14-7 | MolPort-003-983-836 (TargetMol #T6290) | 66576986 | 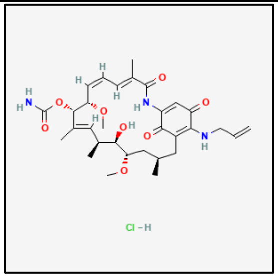 |
| 264834 | 17-DMAG | 467214-20-6 | Cayman #11036 | 53316138 | 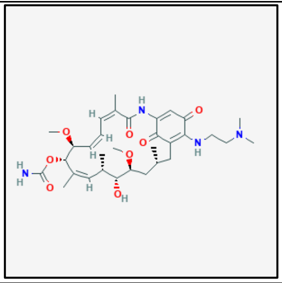 |
| 269784 | PU-H71 | 873436-91-0 | MolPort-042-665-769 (Axon Medchem #1856) | 9549213 | 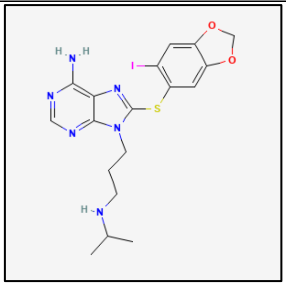 |

| | | | | | |
|---|---|---|---|---|---|
| 273380 | JG-98 | 1456551-16-8 | Gestwicki Lab | 72547053 | 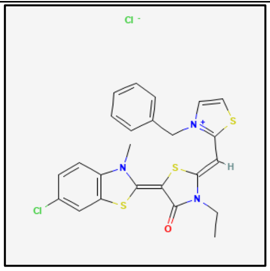 |
| **JG-98: Compounds to test structure activity relationships (SAR)** | | | | | |
| **CCG#** | **Compound Name** | **CAS#** | **Source** | **PubchemID*** | **Pubchem Structure** |
| 208765 | MKT-077 | 147366-41-4 | CCG Library | 6444403 | 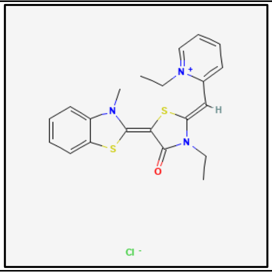 |
| 208766 | YM-1 | 409086-68-6 | Gestwicki Lab & CCG Library | Miyata ACS Chem. Neurosci 2013. | |
| 208767 | YM-8 | 812647-88-4 | CCG Library | Miyata ACS Chem. Neurosci 2013. | |
| - | JG-258 | - | Gestwicki Lab | Shao 2018 J. Med Chem Eftekharzadeh 2019 Nature Communications | |
| - | JG-231 | 1627126-59-3 | Gestwicki Lab | 134817195 | 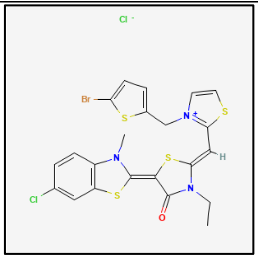 |
| - | JG-345 | | Gestwicki Lab | Shao 2018 J. Med Chem | |
| PTL – Compounds to test SAR | | | | | |
| **CCG#** | **Compound Name** | **CAS#** | **Source** | **Pubchem CID** | **Pubchem Structure** |

| | | | | | |
|---|---|---|---|---|---|
| 36373 | NSC157035 Parthenolide (PTL) | 20554-84-1 | CCG Library | 108068 | 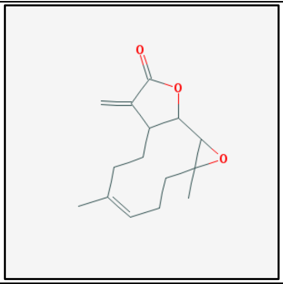 |
| 205026 | Parthenolide | 20554-84-1 | CCG Library | 5420804 | 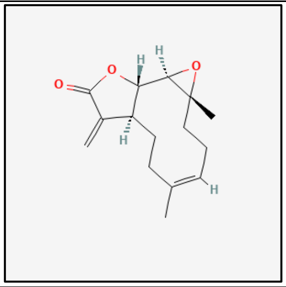 |
| 35235 | NSC 36437 | 28028-68-4 | CCG Library | 6440981 | 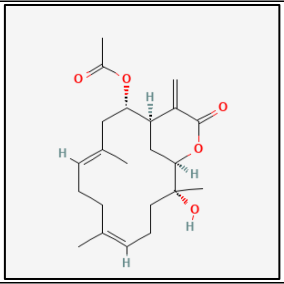 |
| 35566 | NSC 282752 (Eunicin) | 22551-45-7 | CCG Library | 5358827 | 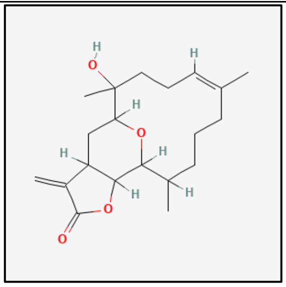 |
| 266942 | Micheliolide | 68370-47-8 | CCG Library | 442279 | 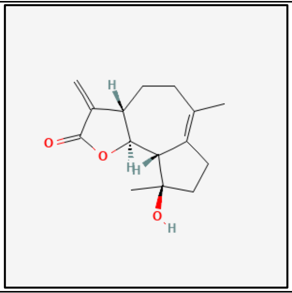 |
| 266810 | Costunolide | 553-21-9 | CCG Library | 5281437 | 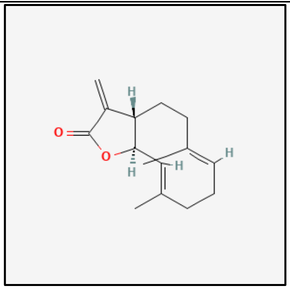 |

| | | | | | |
|---|---|---|---|---|---|
| 208244 | Parthenolide | 20554-84-1 | CCG Library | 7251185 | 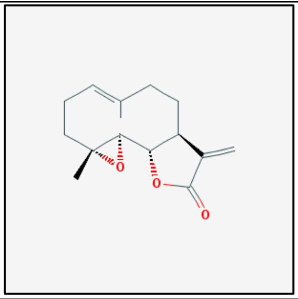 |
| 36035 | NSC 114566 (Eupatundin) | 20071-53-8 | CCG Library | 5281464 | 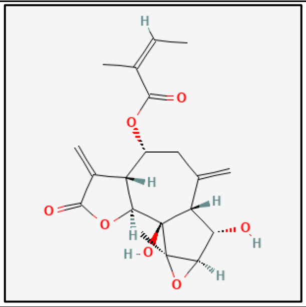 |
| 36184 | NSC 22070 (Pyrethrosin) | 28272-18-6 | CCG library | 5281496 | 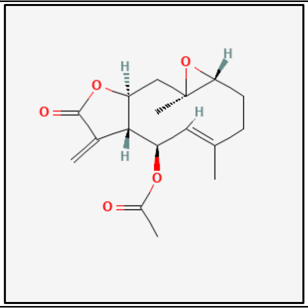 |
| 36376 | NSC302289 (Bohlmann K2631) | 546-43-0 | CCG Library | 327378 | 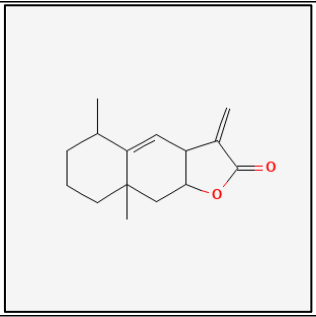 |
| 36468 | NSC93131 (Alantolactone) | 546-43-0 | CCG library | 72724 | 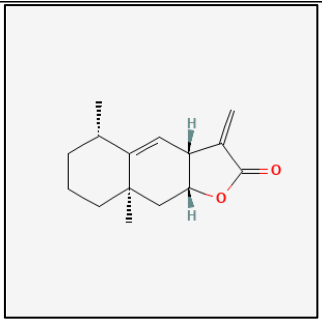 |

| | | | | | |
|---|---|---|---|---|---|
| 266943 | Parthenolide ((-)-Parthenolide) | 20554-84-1 | CCG library | 6473881 | 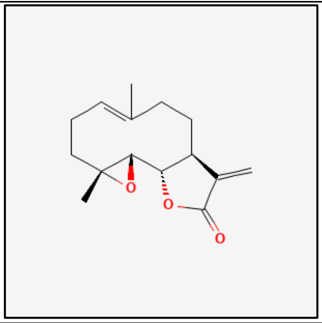 |

**Supplemental Table 4: Initial Hits.**

| Increase MyBP-C/MYH N =70 | CCG_Number | % by Plate | SD_Plate _by_Controls | NAME_STRAIN_GENEID |
|---|---|---|---|---|
| 1 | CCG-38932 | 179.1 | -6.8 | DOXEPIN HYDROCHLORIDE |
| 2 | CCG-40211 | 175.5 | -6.5 | DOPAMINE HYDROCHLORIDE |
| 3 | CCG-39709 | 163.7 | -5.5 | LINCOMYCIN HYDROCHLORIDE |
| 4 | CCG-213826 | 130.4 | -5.4 | TENATOPRAZOLE |
| 5 | CCG-40113 | 162.5 | -5.4 | CARBINOXAMINE MALEATE |
| 6 | CCG-40121 | 158.5 | -5.0 | LEVONORDEFRIN |
| 7 | CCG-39781 | 125.9 | -4.7 | ALLANTOIN |
| 8 | CCG-41384 | 126.1 | -4.5 | RIMANTADINE HYDROCHLORIDE |
| 9 | CCG-39370 | 124.1 | -4.3 | RAMIFENAZONE |
| 10 | CCG-39238 | 121.2 | -3.8 | PROPANTHELINE BROMIDE |
| 11 | CCG-229675 | 143.7 | -3.7 | APIXABAN |
| 12 | CCG-39234 | 142.6 | -3.6 | PROCAINE HYDROCHLORIDE |
| 13 | CCG-39190 | 120.1 | -3.6 | LISINOPRIL |
| 14 | CCG-39483 | 141.8 | -3.6 | METHICILLIN SODIUM |
| 15 | CCG-39237 | 141.6 | -3.6 | PROGESTERONE |
| 16 | CCG-213585 | 120.2 | -3.5 | FLORFENICOL |
| 17 | CCG-34161 | 119.4 | -3.5 | CINCHOPHEN |
| 18 | CCG-229979 | 119.1 | -3.5 | ESOMEPRAZOLE POTASSIUM |
| 19 | CCG-40143 | 119.7 | -3.4 | EQUILIN |
| 20 | CCG-214597 | 139.9 | -3.4 | 3-HYDROXY-4-(SUCCIN-2-YL)-CARYOLANE delta-LACTONE |
| 21 | CCG-39634 | 118.7 | -3.4 | VIDARABINE |
| 22 | CCG-39219 | 137.9 | -3.3 | PHENYTOIN SODIUM |
| 23 | CCG-38999 | 137.2 | -3.2 | METHOXAMINE HYDROCHLORIDE |
| 24 | CCG-38738 | 117.5 | -3.2 | LEVOTHYROXINE |
| 25 | CCG-213210 | 150.5 | -3.1 | IOXILAN |
| 26 | CCG-214635 | 135.7 | -3.1 | 7,4'-DIMETHOXYISOFLAVONE |
| 27 | CCG-38691 | 135.8 | -3.1 | 2',4'-DIHYDROXYCHALCONE 4'-GLUCOSIDE |
| 28 | CCG-212566 | 135.6 | -3.1 | DEXAMETHASONE SODIUM PHOSPHATE |
| 29 | CCG-40328 | 135.7 | -3.1 | BECLOMETHASONE DIPROPIONATE |

| Decrease MyBP-C/MYH (N = 241) | CCG_Number | % by Plate | SD_Plate_ by_Controls | NAME_STRAIN_GENEID |
|---|---|---|---|---|
| 1 | CCG-39264 | 83.3 | 3.0 | SULINDAC |
| 2 | CCG-213547 | 82.6 | 3.0 | DIATRIZOIC ACID |
| 3 | CCG-38728 | 68.4 | 3.0 | DALBERGIONE, 4-METHOXY-4'-HYDROXY- |
| 4 | CCG-38433 | 64.7 | 3.0 | MUUROLLADIE-3-ONE |

| | | | | |
|---|---|---|---|---|
| 5 | CCG-213530 | 82.6 | 3.0 | PANTOTHENIC ACID(d) Na salt |
| 6 | CCG-38983 | 83.0 | 3.0 | ACETAMINOSALOL |
| 7 | CCG-38984 | 83.0 | 3.0 | ACETANILIDE |
| 8 | CCG-204168 | 83.2 | 3.0 | RESERPINE |
| 9 | CCG-39018 | 83.0 | 3.1 | MEMANTINE HYDROCHLORIDE |
| 10 | CCG-213542 | 82.3 | 3.1 | ERYTHROSINE SODIUM |
| 11 | CCG-213612 | 82.4 | 3.1 | PROTIRELIN |
| 12 | CCG-264834 | 43.0 | 3.1 | 17-DMAG |
| 13 | CCG-213614 | 82.3 | 3.1 | ACESULFAME POTASSIUM |
| 14 | CCG-214274 | 63.9 | 3.1 | DIFUCOL HEXAMETHYL ETHER |
| 15 | CCG-213453 | 82.2 | 3.1 | PYRIDOXINE |
| 16 | CCG-213690 | 82.2 | 3.1 | DIFLORASONE DIACETATE |
| 17 | CCG-213742 | 82.6 | 3.1 | PIPEMIDIC ACID |
| 18 | CCG-213778 | 82.6 | 3.1 | BENFOTIAMINE |
| 19 | CCG-214375 | 67.4 | 3.1 | PIPLARTINE |
| 20 | CCG-213992 | 82.5 | 3.1 | CYROMAZINE |
| 21 | CCG-39708 | 82.6 | 3.1 | ACETARSOL |
| 22 | CCG-39017 | 82.7 | 3.1 | MEFEXAMIDE HYDROCHLORIDE |
| 23 | CCG-213228 | 48.4 | 3.1 | THONZONIUM BROMIDE |
| 24 | CCG-213989 | 82.4 | 3.2 | NIFUROXAZIDE |
| 25 | CCG-39629 | 82.6 | 3.2 | TAMOXIFEN CITRATE |
| 26 | CCG-40150 | 82.3 | 3.2 | ZOXAZOLAMINE |
| 27 | CCG-39983 | 82.3 | 3.2 | MECYSTEINE HYDROCHLORIDE |
| 28 | CCG-213464 | 81.7 | 3.2 | IFOSFAMIDE |
| 29 | CCG-38526 | 62.8 | 3.2 | GARCINOLIC ACID |
| 30 | CCG-39973 | 82.1 | 3.2 | BETAMIPRON |
| 31 | CCG-39265 | 82.3 | 3.2 | TERBUTALINE HEMISULFATE |
| 32 | CCG-40148 | 66.6 | 3.2 | CHRYSIN |
| 33 | CCG-213527 | 81.6 | 3.2 | PREDNISOLONE SODIUM PHOSPHATE |
| 34 | CCG-213993 | 82.0 | 3.2 | PROXYPHYLLINE |
| 35 | CCG-36432 | 82.0 | 3.2 | PIDOLIC ACID |
| 36 | CCG-38995 | 82.2 | 3.2 | SULFAMONOMETHOXINE |
| 37 | CCG-231323 | 66.4 | 3.2 | 16(17)-EPOXY-5-PREGNENOLONE |
| 38 | CCG-213671 | 81.5 | 3.2 | CAPTAMINE |
| 39 | CCG-100832 | 82.0 | 3.2 | PIDOTIMOD |
| 40 | CCG-40004 | 62.4 | 3.2 | 3-DESMETHYL-5-DESHYDROXYSCLEROIN |
| 41 | CCG-213456 | 81.5 | 3.2 | SULFISOXAZOLE ACETYL |
| 42 | CCG-213438 | 81.3 | 3.3 | HALOTHANE |
| 43 | CCG-100878 | 81.7 | 3.3 | BUFLOMEDIL HYDROCHLORIDE |
| 44 | CCG-38959 | 81.7 | 3.3 | CLOFIBRIC ACID |

| | | | | |
|---|---|---|---|---|
| 45 | CCG-213218 | 46.3 | 3.3 | ZIPRASIDONE MESYLATE |
| 46 | CCG-39745 | 81.7 | 3.3 | AMINOPTERIN |
| 47 | CCG-213463 | 81.2 | 3.3 | CEFONICID SODIUM |
| 48 | CCG-40146 | 61.5 | 3.3 | AMODIAQUINE DIHYDROCHLORIDE |
| 49 | CCG-39147 | 81.1 | 3.3 | DICHLORVOS |
| 50 | CCG-213445 | 81.1 | 3.3 | METHYLENE BLUE |
| 51 | CCG-38919 | 61.4 | 3.3 | AMITRIPTYLINE HYDROCHLORIDE |
| 52 | CCG-213675 | 81.0 | 3.3 | AMIKACIN HYDRATE |
| 53 | CCG-39774 | 81.7 | 3.3 | SISOMICIN SULFATE |
| 54 | CCG-39322 | 45.5 | 3.3 | METHYLBENZETHONIUM CHLORIDE |
| 55 | CCG-39610 | 80.9 | 3.3 | ALPRENOLOL |
| 56 | CCG-213741 | 81.4 | 3.3 | RUTIN |
| 57 | CCG-220176 | 81.4 | 3.3 | IPRONIAZID PHOSPHATE |
| 58 | CCG-39493 | 80.8 | 3.3 | PINACIDIL |
| 59 | CCG-39694 | 81.3 | 3.3 | METHOPRENE (S) |
| 60 | CCG-38893 | 60.9 | 3.3 | BATYL ALCOHOL |
| 61 | CCG-212786 | 81.4 | 3.4 | RIFAMPIN |
| 62 | CCG-269784 | 37.6 | 3.4 | PU-H71 |
| 63 | CCG-230630 | 80.6 | 3.4 | EDITOL |
| 64 | CCG-213034 | 81.3 | 3.4 | RASAGILINE |
| 65 | CCG-214537 | 60.6 | 3.4 | 4,4'-DIISOTHIOCYANOSTILBENE-2,2'-SUFONIC ACID SODIUM SALT |
| 66 | CCG-208039 | 37.4 | 3.4 | 17-AAG |
| 67 | CCG-39272 | 81.2 | 3.4 | TIMOLOL MALEATE |
| 68 | CCG-38529 | 64.2 | 3.4 | CARMINIC ACID |
| 69 | CCG-213493 | 43.6 | 3.4 | FLUTICASONE PROPIONATE |
| 70 | CCG-38622 | 64.1 | 3.4 | CYTIDINE |
| 71 | CCG-204551 | 80.0 | 3.5 | ETHAMIVAN |
| 72 | CCG-213221 | 79.9 | 3.5 | DECOQUINATE [5mM] |
| 73 | CCG-213019 | 79.8 | 3.5 | FULVESTRANT |
| 74 | CCG-39523 | 80.3 | 3.5 | PRIDINOL METHANESULFONATE |
| 75 | CCG-40209 | 79.7 | 3.5 | DISOPYRAMIDE PHOSPHATE |
| 76 | CCG-207958 | 80.2 | 3.5 | DOCONEXENT |
| 77 | CCG-39446 | 79.4 | 3.6 | TRIADIMEFON |
| 78 | CCG-213427 | 79.5 | 3.6 | BURAMATE |
| 79 | CCG-213477 | 79.5 | 3.6 | DILOXANIDE FUROATE |
| 80 | CCG-213982 | 80.0 | 3.6 | CHLOROPYRAMINE HYDROCHLORIDE |
| 81 | CCG-213027 | 80.3 | 3.6 | NIFURSOL |
| 82 | CCG-40109 | 80.0 | 3.6 | ABAMECTIN (avermectin B1a shown) |
| 83 | CCG-214000 | 79.7 | 3.6 | IFENPRODIL TARTRATE |
| 84 | CCG-39260 | 79.9 | 3.6 | SULFAMETHIZOLE |

| | | | | |
|---|---|---|---|---|
| 85 | CCG-214610 | 61.9 | 3.6 | ERYTHROSE |
| 86 | CCG-39344 | 40.1 | 3.6 | BENZALKONIUM CHLORIDE |
| 87 | CCG-40292 | 61.6 | 3.7 | HUPERZINE A |
| 88 | CCG-264804 | 31.8 | 3.7 | NVP-AUY922 |
| 89 | CCG-231621 | 78.7 | 3.7 | CAPTAN |
| 90 | CCG-39715 | 79.2 | 3.7 | PENTAMIDINE ISETHIONATE |
| 91 | CCG-214765 | 78.6 | 3.7 | HYDROQUININE HYDROBROMIDE HYDRATE |
| 92 | CCG-39082 | 38.8 | 3.7 | HYDROQUINONE |
| 93 | CCG-40115 | 56.5 | 3.7 | NIFEDIPINE |
| 94 | CCG-39305 | 79.1 | 3.7 | BROMOPRIDE |
| 95 | CCG-39117 | 79.1 | 3.7 | ERYTHROMYCIN STEARATE |
| 96 | CCG-39630 | 79.0 | 3.8 | HEXESTROL |
| 97 | CCG-214611 | 60.7 | 3.8 | GARDENIN B |
| 98 | CCG-38376 | 56.1 | 3.8 | EPICATECHIN MONOGALLATE |
| 99 | CCG-213448 | 78.4 | 3.8 | MINOXIDIL HYDROCHLORIDE |
| 100 | CCG-212466 | 78.9 | 3.8 | PYRETHRINS |
| 101 | CCG-213566 | 78.3 | 3.8 | ATRACURIUM BESYLATE |
| 102 | CCG-39494 | 78.3 | 3.8 | LOPERAMIDE HYDROCHLORIDE |
| 103 | CCG-213831 | 78.8 | 3.8 | SODIUM CYCLAMATE |
| 104 | CCG-213561 | 78.2 | 3.8 | IRINOTECAN HYDROCHLORIDE |
| 105 | CCG-213428 | 78.1 | 3.8 | CAPOBENIC ACID |
| 106 | CCG-38835 | 78.7 | 3.8 | HYDROQUINIDINE |
| 107 | CCG-213834 | 78.6 | 3.8 | CYACETACIDE |
| 108 | CCG-213032 | 78.9 | 3.8 | QUINESTROL |
| 109 | CCG-39531 | 37.1 | 3.8 | AMLODIPINE BESYLATE |
| 110 | CCG-39077 | 77.8 | 3.8 | CHLOROTHALONIL |
| 111 | CCG-208032 | 59.8 | 3.9 | PLUMBAGIN |
| 112 | CCG-213231 | 36.7 | 3.9 | PREDNICARBATE |
| 113 | CCG-39001 | 78.2 | 3.9 | GLAFENINE |
| 114 | CCG-208156 | 77.5 | 3.9 | ARTESUNATE |
| 115 | CCG-40051 | 58.9 | 3.9 | DIHYDROCELASTRYL DIACETATE |
| 116 | CCG-213915 | 77.9 | 4.0 | FORMESTANE |
| 117 | CCG-40182 | 77.9 | 4.0 | DROPROPIZINE |
| 118 | CCG-213983 | 77.8 | 4.0 | FAMPROFAZONE |
| 119 | CCG-204987 | 77.8 | 4.0 | PALMIDROL [5mM] |
| 120 | CCG-273380 | 26.4 | 4.0 | JG-98 |
| 121 | CCG-213835 | 77.6 | 4.0 | ACECLOFENAC |
| 122 | CCG-39550 | 77.6 | 4.0 | ACEMETACIN |
| 123 | CCG-38354 | 77.9 | 4.0 | ARECOLINE HYDROBROMIDE |
| 124 | CCG-213543 | 76.8 | 4.0 | GLYCOPYRROLATE |

| | | | | |
|---|---|---|---|---|
| 125 | CCG-39945 | 77.3 | 4.1 | PIMETHIXENE MALEATE |
| 126 | CCG-213832 | 77.2 | 4.1 | ORBIFLOXACIN |
| 127 | CCG-213252 | 32.7 | 4.1 | TACROLIMUS |
| 128 | CCG-213487 | 76.3 | 4.1 | TIOCONAZOLE |
| 129 | CCG-40159 | 76.0 | 4.2 | 2,6-DIMETHOXYQUINONE |
| 130 | CCG-269859 | 3.9 | 4.2 | NMS-873 |
| 131 | CCG-213436 | 75.7 | 4.2 | FOMEPIZOLE HYDROCHLORIDE |
| 132 | CCG-39746 | 76.5 | 4.2 | PUROMYCIN DIHYDROCHLORIDE |
| 133 | CCG-39480 | 76.2 | 4.3 | QUININE ETHYL CARBONATE |
| 134 | CCG-39784 | 76.1 | 4.3 | OCTOPAMINE HYDROCHLORIDE |
| 135 | CCG-39719 | 49.4 | 4.3 | CETYLPYRIDINIUM CHLORIDE |
| 136 | CCG-213545 | 75.0 | 4.4 | OCTISALATE |
| 137 | CCG-38787 | 54.2 | 4.4 | CURCUMIN |
| 138 | CCG-39674 | 53.9 | 4.4 | STROPHANTHIDIN |
| 139 | CCG-39227 | 48.2 | 4.4 | PRAZOSIN HYDROCHLORIDE |
| 140 | CCG-40298 | 75.1 | 4.5 | AMINOTHIAZOLE |
| 141 | CCG-39135 | 73.7 | 4.6 | AMOXAPINE |
| 142 | CCG-39608 | 73.9 | 4.7 | CORTISONE |
| 143 | CCG-213454 | 72.6 | 4.8 | PHENSUCCIMIDE |
| 144 | CCG-213110 | 21.5 | 4.8 | BEPRIDIL HYDROCHLORIDE |
| 145 | CCG-40224 | 73.1 | 4.8 | FENDILINE HYDROCHLORIDE |
| 146 | CCG-214004 | 73.0 | 4.8 | OXINIACIC ACID |
| 147 | CCG-38364 | 49.4 | 4.8 | QUERCETIN 5,7,3',4'-TETRAMETHYL ETHER |
| 148 | CCG-39878 | 19.3 | 4.9 | THALIDOMIDE |
| 149 | CCG-40333 | 72.5 | 4.9 | BROXYQUINOLINE |
| 150 | CCG-39155 | 72.2 | 5.0 | BENFLUOREX HYDROCHLORIDE |
| 151 | CCG-208040 | 7.6 | 5.0 | Geldanamycin |
| 152 | CCG-39161 | 72.1 | 5.0 | EBSELEN |
| 153 | CCG-213695 | 71.4 | 5.0 | OXTRIPHYLLINE |
| 154 | CCG-40020 | 17.4 | 5.0 | OXICONAZOLE NITRATE |
| 155 | CCG-213424 | 71.0 | 5.1 | BISMUTH SUBSALICYLATE |
| 156 | CCG-231419 | 46.9 | 5.1 | DIMETHYL GAMBOGINATE |
| 157 | CCG-39927 | 5.6 | 5.1 | CANTHARIDIN |
| 158 | CCG-39254 | 71.5 | 5.2 | SPIRONOLACTONE |
| 159 | CCG-40290 | 71.4 | 5.2 | TRIFLUOPERAZINE HYDROCHLORIDE |
| 160 | CCG-38542 | 38.0 | 5.3 | DIHYDROGAMBOGIC ACID |
| 161 | CCG-39361 | 12.0 | 5.4 | SELAMECTIN |
| 162 | CCG-39317 | 10.9 | 5.4 | PERHEXILINE MALEATE |
| 163 | CCG-213042 | 69.7 | 5.5 | URACIL |
| 164 | CCG-40027 | 36.0 | 5.5 | THYMOQUINONE |

| | | | | |
|---|---|---|---|---|
| 165 | CCG-214012 | 68.5 | 5.5 | TROMETHAMINE |
| 166 | CCG-220939 | 69.7 | 5.5 | IODIPAMIDE |
| 167 | CCG-39124 | 9.8 | 5.5 | NONOXYNOL-9 |
| 168 | CCG-214002 | 68.3 | 5.7 | IDAZOXAN HYDROCHLORIDE |
| 169 | CCG-213841 | 67.9 | 5.7 | BORNYL ACETATE |
| 170 | CCG-38966 | 68.1 | 5.8 | ECONAZOLE NITRATE |
| 171 | CCG-213700 | 66.7 | 5.8 | AUROTHIOGLUCOSE |
| 172 | CCG-39639 | 39.0 | 5.8 | HELENINE |
| 173 | CCG-39846 | 30.8 | 5.9 | GITOXIGENIN |
| 174 | CCG-39763 | 30.8 | 5.9 | GENTIAN VIOLET |
| 175 | CCG-207937 | 65.7 | 6.0 | BUTYLATED HYDROXYTOLUENE |
| 176 | CCG-39553 | 28.8 | 6.1 | CLOMIPHENE CITRATE |
| 177 | CCG-38636 | 27.7 | 6.2 | GITOXIGENIN DIACETATE |
| 178 | CCG-38362 | 26.6 | 6.3 | DIGITOXIN |
| 179 | CCG-38826 | 25.8 | 6.4 | DIGOXIN |
| 180 | CCG-213570 | 23.7 | 6.5 | DORAMECTIN |
| 181 | CCG-38852 | 23.3 | 6.6 | GITOXIN |
| 182 | CCG-38382 | 30.5 | 6.6 | AVOCATIN A |
| 183 | CCG-214256 | 28.7 | 6.8 | DIGOXIGENIN |
| 184 | CCG-38646 | 18.4 | 7.0 | STROPHANTHIDINIC ACID LACTONE ACETATE |
| 185 | CCG-38819 | 17.2 | 7.1 | CONVALLATOXIN |
| 186 | CCG-39020 | 17.1 | 7.1 | PERUVOSIDE |
| 187 | CCG-231469 | 24.9 | 7.2 | EMICYMARIN |
| 188 | CCG-212631 | 14.6 | 7.3 | GRAMICIDIN (gramicidin A shown) |
| 189 | CCG-39756 | 13.6 | 7.4 | MICONAZOLE NITRATE |
| 190 | CCG-40283 | 13.1 | 7.4 | PRIMAQUINE PHOSPHATE |
| 191 | CCG-40047 | 56.9 | 7.5 | DIBENZOYLMETHANE |
| 192 | CCG-231353 | 19.6 | 7.7 | STROPHANTHIDIN ACETATE |
| 193 | CCG-38545 | 8.5 | 7.8 | PYRITHIONE ZINC |
| 194 | CCG-214431 | 4.9 | 8.1 | AVERMECTIN A1a |
| 195 | CCG-213819 | 53.6 | 8.3 | OLTIPRAZ |
| 196 | CCG-212462 | 3.2 | 8.3 | SANGUINARIUM CHLORIDE |
| 197 | CCG-214187 | 2.6 | 8.3 | METHYL GAMBOGATE METHYL ETHER |
| 198 | CCG-39830 | 1.4 | 8.4 | TOTAROL |
| 199 | CCG-39917 | 1.3 | 8.4 | DIHYDROCELASTROL |
| 200 | CCG-39638 | 1.1 | 8.5 | PARTHENOLIDE |
| 201 | CCG-38944 | 1.1 | 8.5 | OXYPHENBUTAZONE |
| 202 | CCG-36465 | 0.9 | 8.5 | alpha-MANGOSTIN |
| 203 | CCG-39919 | -0.1 | 8.6 | CELASTROL |
| 204 | CCG-39752 | 51.8 | 8.6 | HEXETIDINE |

| | | | | |
|---|---|---|---|---|
| 205 | CCG-39613 | 8.2 | 8.8 | 4-NONYLPHENOL |
| 206 | CCG-214482 | 8.0 | 8.8 | METHYL GAMBOGATE |
| 207 | CCG-38640 | 4.1 | 9.2 | TOTAROL-19-CARBOXYLIC ACID, METHYL ESTER |
| 208 | CCG-39847 | 3.5 | 9.2 | MUNDULONE |
| 209 | CCG-36362 | 3.4 | 9.2 | beta-LAPACHONE |
| 210 | CCG-38659 | 2.9 | 9.3 | GAMBOGIC ACID |
| 211 | CCG-36078 | 1.7 | 9.4 | POMIFERIN |
| 212 | CCG-214401 | 1.3 | 9.4 | ISOOSAJIN |
| 213 | CCG-38435 | 1.3 | 9.4 | CEDROL |
| 214 | CCG-39987 | 32.3 | 11.7 | CHLORANIL |
| 215 | CCG-213416 | 31.4 | 12.0 | MOXIDECTIN |
| 216 | CCG-40300 | 32.6 | 12.0 | LEVOMENTHOL |
| 217 | CCG-39607 | 28.3 | 12.5 | AMIODARONE HYDROCHLORIDE |
| 218 | CCG-39311 | 27.5 | 12.9 | CLOFOCTOL |
| 219 | CCG-230895 | 25.4 | 13.3 | OUABAIN OCTAHYDRATE |
| 220 | CCG-35584 | 25.1 | 13.4 | MILTEFOSINE [5mM] |
| 221 | CCG-38866 | 25.8 | 13.4 | LANATOSIDE C |
| 222 | CCG-214013 | 24.8 | 13.4 | BUCETIN |
| 223 | CCG-230495 | 19.1 | 14.1 | PROSCILLARIDIN A |
| 224 | CCG-213889 | 20.7 | 14.2 | ORNITHINE HYDROCHLORIDE |
| 225 | CCG-39619 | 20.4 | 14.2 | REBAMIPIDE |
| 226 | CCG-213774 | 18.9 | 14.5 | gamma-AMINOBUTYRIC ACID HYDROCHLORIDE |
| 227 | CCG-39201 | 13.6 | 15.0 | 1-BENZYLOXYCARBONYLAMINOPHENETHYL CHLOROMETHYL KETONE |
| 228 | CCG-40014 | 9.9 | 15.7 | MITOTANE |
| 229 | CCG-213865 | 10.4 | 16.0 | CITIOLONE |
| 230 | CCG-40030 | 3.2 | 16.8 | ELAIDYLPHOSPHOCHOLINE |
| 231 | CCG-39862 | 2.6 | 17.0 | LINDANE |
| 232 | CCG-213459 | 2.2 | 17.0 | TRICLOSAN |
| 233 | CCG-213693 | 2.0 | 17.1 | IVERMECTIN |
| 234 | CCG-101015 | 1.5 | 17.1 | PTEROSTILBENE |
| 235 | CCG-101072 | 1.8 | 17.1 | TOREMIFENE CITRATE |
| 236 | CCG-213065 | 4.5 | 17.3 | EPRINOMECTIN |
| 237 | CCG-38719 | 3.1 | 17.3 | ESCIN |
| 238 | CCG-39270 | 1.8 | 17.7 | THIORIDAZINE HYDROCHLORIDE |
| 239 | CCG-39015 | 1.7 | 17.8 | SULCONAZOLE NITRATE |
| 240 | CCG-39725 | -1.1 | 18.3 | THIMEROSAL |
| 241 | CCG-39751 | -48.7 | 26.9 | PHENYLMERCURIC ACETATE |

SD = standard deviation, %_by_Plate where 100% indicated MyBP-C/MYH signal for *MYBPC3* Patient1 iPSC-CMs vehicle control for that plate and 0% indicates MyBP-C/MYH signal for *MYBPC3* (-/-) vehicle control for that plate.

### Supplemental Table 5: Validated by retesting – Compounds that decrease MyBP-C/MYH

| Count | CCG_Number | NAME_STRAIN_GENEID |
|---|---|---|
| 1 | CCG-264834 | 17-DMAG |
| 2 | CCG-269784 | PU-H71 |
| 3 | CCG-208039 | 17-AAG |
| 4 | CCG-264804 | NVP-AUY922 |
| 5 | CCG-40113 | CARBINOXAMINE MALEATE |
| 6 | CCG-40146 | AMODIAQUINE DIHYDROCHLORIDE |
| 7 | CCG-39490 | ASPIRIN |
| 8 | CCG-273380 | JG98 |
| 9 | CCG-213835 | ACECLOFENAC |
| 10 | CCG-40051 | DIHYDROCELASTRYL DIACETATE |
| 11 | CCG-39719 | CETYLPYRIDINIUM CHLORIDE |
| 12 | CCG-40298 | AMINOTHIAZOLE |
| 13 | CCG-213210 | IOXILAN |
| 14 | CCG-39082 | HYDROQUINONE |
| 15 | CCG-40333 | BROXYQUINOLINE |
| 16 | CCG-39674 | STROPHANTHIDIN |
| 17 | CCG-208040 | Geldanamycin |
| 18 | CCG-39927 | Cantharidin |
| 19 | CCG-269859 | NMS-873 |
| 20 | CCG-39124 | NONOXYNOL-9 |
| 21 | CCG-39846 | GITOXIGENIN |
| 22 | CCG-39317 | PERHEXILINE MALEATE |
| 23 | CCG-38636 | GITOXIGENIN DIACETATE |
| 24 | CCG-38826 | DIGOXIN |
| 25 | CCG-38362 | DIGITOXIN |
| 26 | CCG-38646 | STROPHANTHIDINIC ACID LACTONE ACETATE |
| 27 | CCG-208032 | PLUMBAGIN |
| 28 | CCG-38819 | CONVALLATOXIN |
| 29 | CCG-39020 | PERUVOSIDE |
| 30 | CCG-231469 | EMICYMARIN |
| 31 | CCG-39756 | MICONAZOLE NITRATE |
| 32 | CCG-214256 | DIGOXIGENIN |
| 33 | CCG-40283 | PRIMAQUINE PHOSPHATE |
| 34 | CCG-231353 | STROPHANTHIDIN ACETATE |
| 35 | CCG-38545 | PYRITHIONE ZINC |
| 36 | CCG-212462 | SANGUINARIUM CHLORIDE |
| 37 | CCG-39917 | DIHYDROCELASTROL |
| 38 | CCG-39638 | PARTHENOLIDE |
| 39 | CCG-36465 | alpha-MANGOSTIN |
| 40 | CCG-39919 | CELASTROL |
| 41 | CCG-39763 | GENTIAN VIOLET |
| 42 | CCG-214482 | METHYL GAMBOGATE |
| 43 | CCG-230895 | OUABAIN OCTAHYDRATE |

| | | |
|---|---|---|
| 44 | CCG-38866 | LANATOSIDE C |
| 45 | CCG-230495 | PROSCILLARIDIN A |
| 46 | CCG-39619 | REBAMIPIDE |
| 47 | CCG-213774 | gamma-AMINOBUTYRIC ACID HYDROCHLORIDE |
| 48 | CCG-213459 | TRICLOSAN |
| 49 | CCG-101015 | PTEROSTILBENE |
| 50 | CCG-38719 | ESCIN |
| 51 | CCG-39725 | THIMEROSAL |
| 52 | CCG-39751 | PHENYLMERCURIC ACETATE |

Criteria: MyBP-C/MYH > 3 stdev below vehicle control for three of four available data points.

**Supplemental Table 6: Evaluating MyBP-C and MYH assay results for validated MyBP-C/MYH compounds**

**Active MyBP-C, inactive MYH**

| Count | CCG_Number | MyBP-C % by_Plate | MyBP-C SD_Plate by_Controls | NAME_STRAIN_GENEID |
|---|---|---|---|---|
| 1 | CCG-40298 | 72.6 | 3.87 | AMINOTHIAZOLE |
| 2 | CCG-264834 | 32 | 5.55 | 17-DMAG |
| 3 | CCG-208039 | 30 | 5.71 | 17-AAG |
| 4 | CCG-269784 | 23 | 6.28 | PU-H71 |
| 5 | CCG-264804 | 17 | 6.77 | NVP-AUY922 |
| 6 | CCG-273380 | 15 | 6.94 | JG98 |
| 7 | CCG-39082 | 43.5 | 7.98 | HYDROQUINONE |
| 8 | CCG-39719 | 29.4 | 9.97 | CETYLPYRIDINIUM CHLORIDE |
| 9 | CCG-39846 | 25.6 | 10.93 | GITOXIGENIN |
| 10 | CCG-39124 | 19.1 | 11.88 | NONOXYNOL-9 |
| 11 | CCG-38636 | 18.8 | 11.93 | GITOXIGENIN DIACETATE |
| 12 | CCG-38646 | 15.8 | 12.37 | STROPHANTHIDINIC ACID LACTONE ACETATE |
| 13 | CCG-40051 | 44.1 | 12.56 | DIHYDROCELASTRYL DIACETATE |
| 14 | CCG-230895 | 13.9 | 13.7 | OUABAIN OCTAHYDRATE |
| 15 | CCG-39763 | 1.2 | 13.95 | GENTIAN VIOLET |
| 16 | CCG-39917 | 3.9 | 14.12 | DIHYDROCELASTROL |
| 17 | CCG-39919 | 3.5 | 14.17 | CELASTROL |
| 18 | CCG-39638 | 1.7 | 14.43 | PARTHENOLIDE |
| 19 | CCG-40283 | 1.7 | 14.45 | PRIMAQUINE PHOSPHATE |
| 20 | CCG-38545 | 0.9 | 14.56 | PYRITHIONE ZINC |
| 21 | CCG-212462 | 0.1 | 14.67 | SANGUINARIUM CHLORIDE |
| 22 | CCG-101015 | 0.7 | 16.69 | PTEROSTILBENE |
| 23 | CCG-214482 | 0.4 | 17.24 | METHYL GAMBOGATE |

**Active MyBP-C,**
**excluded due to > 3 stdev change in MYH assay above or below control**

| Count | CCG_Number | MyBP-C % by_Plate | MyBP-C SD_Plate by_Controls | NAME_STRAIN_GENEID |
|---|---|---|---|---|
| 1 | CCG-38362 | 9.1 | 12.84 | DIGITOXIN |
| 2 | CCG-38866 | 8.9 | 12.87 | LANATOSIDE C |
| 3 | CCG-38826 | 11.7 | 12.98 | DIGOXIN |
| 4 | CCG-230495 | 11.3 | 13.03 | PROSCILLARIDIN A |
| 5 | CCG-39020 | 10.1 | 13.2 | PERUVOSIDE |
| 6 | CCG-38819 | 9.2 | 13.34 | CONVALLATOXIN |
| 7 | CCG-39317 | 2.2 | 14.36 | PERHEXILINE MALEATE |
| 8 | CCG-213459 | 1.1 | 14.52 | TRICLOSAN |
| 9 | CCG-39756 | 0.9 | 14.55 | MICONAZOLE NITRATE |
| 10 | CCG-36465 | 0.7 | 14.59 | alpha-MANGOSTIN |
| 11 | CCG-214256 | 15.1 | 14.69 | DIGOXIGENIN |
| 12 | CCG-39751 | -0.2 | 14.71 | PHENYLMERCURIC ACETATE |

| | | | | |
|---|---|---|---|---|
| 13 | CCG-39725 | -0.1 | 14.71 | THIMEROSAL |
| 14 | CCG-231353 | 9.1 | 15.74 | STROPHANTHIDIN ACETATE |
| 15 | CCG-39674 | 14.6 | 19.20 | STROPHANTHIDIN |
| 16 | CCG-208040 | 3 | 7.91 | GELDANAMYCIN |
| 17 | CCG-269859 | 1 | 8.08 | NMS-873 |
| 18 | CCG-39927 | 2 | 8.00 | CANTHARIDIN |

**Supplemental Table 7: CRC results**
**MyBP-C/MYH**

| Count | CCG_Number | EC50(μM) | 95% CI | Bottom | 95% CI | NAME_STRAIN_GENEID |
|---|---|---|---|---|---|---|
| *1* | *CCG-40298* | *inactive* | - | - | - | *AMINOTHIAZOLE* |
| **2** | **CCG-264834** | **1.1** | **0.1-4.7** | **41.2** | **12.3-61.8** | **17-DMAG** |
| **3** | **CCG-208039** | **4.7** | **1.3-12.4** | **33.1** | **12.6-59.7** | **17-AAG** |
| **4** | **CCG-269784** | **1.3** | **0.2-5.5** | **35.2** | **2.9-57.1** | **PU-H71** |
| 5 | CCG-264804 | 1.7 | 0-20 | 43.7 | 0-74.4 | NVP-AUY922 |
| **6** | **CCG-273380** | **25.5** | **9.6-52.1** | **0** | **NA** | **JG98** |
| **7** | **CCG-39082** | **23.1** | **13.7-36.8** | **0** | **NA** | **HYDROQUINONE** |
| **8** | **CCG-39719** | **10.6** | **7.7-14.3** | **0** | **NA** | **CETYLPYRIDINIUM CHLORIDE** |
| *9* | *CCG-39846* | *Inactive* | - | - | - | *GITOXIGENIN* |
| 10 | CCG-39124 | 24.4 | 10.4-50.2 | 0 | NA | NONOXYNOL-9 |
| *11* | *CCG-38636* | *inactive* | - | - | - | *GITOXIGENIN DIACETATE* |
| 12 | CCG-38646 | indeterminate | - | - | - | STROPHANTHIDINIC ACID LACTONE ACETATE |
| **13** | **CCG-40051** | **10.2** | **5.9-15.1** | **0** | **NA** | **DIHYDROCELASTRYL DIACETATE** |
| 14 | CCG-230895 | 0.8 | 0.1-1.8 | 37.6 | 31.1-43.6 | OUABAIN OCTAHYDRATE |
| **15** | **CCG-39763** | **14.6** | **8.3-24.3** | **0** | **NA** | **GENTIAN VIOLET** |
| **16** | **CCG-39917** | **8.0** | **5.1-11.9** | **0** | **NA** | **DIHYDROCELASTROL** |
| **17** | **CCG-39919** | **7.0** | **4.8-9.8** | **0** | **NA** | **CELASTROL** |
| **18** | **CCG-39638** | **8.5** | **5.1-11.9** | **0** | **NA** | **PARTHENOLIDE** |
| **19** | **CCG-40283** | **19.7** | **11.1-32.1** | **0** | **NA** | **PRIMAQUINE PHOSPHATE** |
| **20** | **CCG-38545** | **15.6** | **5.9-32.1** | **0** | **NA** | **PYRITHIONE ZINC** |
| 21 | CCG-212462 | 7.2 | 2.2-13.4 | 0 | NA | SANGUINARIUM CHLORIDE |
| 22 | CCG-101015 | 16.8 | 7.0-33.2 | 0 | NA | PTEROSTILBENE |
| **23** | **CCG-214482** | **8.4** | **6.0-11.5** | **0** | **NA** | **METHYL GAMBOGATE** |

**MyBP-C**

| Count | CCG_Number | EC50(μM) | 95% CI | Bottom | 95% CI | NAME_STRAIN_GENEID |
|---|---|---|---|---|---|---|
| *1* | *CCG-40298* | *inactive* | - | - | - | *AMINOTHIAZOLE* |
| **2** | **CCG-264834** | **4.3** | **0.8-14.7** | **16.0** | **0-54.9** | **17-DMAG** |
| **3** | **CCG-208039** | **8.2** | **2.2-24.5** | **16.5** | **0-43.0** | **17-AAG** |
| **4** | **CCG-269784** | **4.8** | **1.2-14.6** | **18.6** | **0-50.9** | **PU-H71** |

| | | | | | | |
|---|---|---|---|---|---|---|
| 5 | CCG-264804 | 1.6 | 0.4-5.5 | 32.6 | 2.6-53.2 | NVP-AUY922 |
| **6** | **CCG-273380** | **20.8** | **13.1-30.5** | **0** | **NA** | **JG98** |
| **7** | **CCG-39082** | **16.9** | **10.7-23.9** | **0** | **NA** | **HYDROQUINONE** |
| **8** | **CCG-39719** | **7.4** | **3.7-11.7** | **0** | **NA** | **CETYLPYRIDINIUM CHLORIDE** |
| *9* | *CCG-39846* | *41.5* | *15.9-61.7* | *0* | *NA* | *GITOXIGENIN* |
| 10 | CCG-39124 | 24.4 | 10.4-50.2 | 0 | NA | NONOXYNOL-9 |
| *11* | *CCG-38636* | *4.7* | *2.0-9.4* | *18.6* | *3.6-30.9* | *GITOXIGENIN DIACETATE* |
| 12 | CCG-38646 | 4.0 | 2.1-6.8 | 15.4 | 4.4-25.2 | STROPHANTHIDINIC ACID LACTONE ACETATE |
| **13** | **CCG-40051** | **8.9** | **6.0-12.5** | **0** | **NA** | **DIHYDROCELASTRYL DIACETATE** |
| 14 | CCG-230895 | 0.42 | 0.1-0.8 | 21.3 | 17.9-24.7 | OUABAIN OCTAHYDRATE |
| **15** | **CCG-39763** | **6.5** | **4.3-8.9** | **0** | **NA** | **GENTIAN VIOLET** |
| **16** | **CCG-39917** | **5.9** | **4.3-8.1** | **0** | **NA** | **DIHYDROCELASTROL** |
| **17** | **CCG-39919** | **6.0** | **4.0-8.6** | **0** | **NA** | **CELASTROL** |
| **18** | **CCG-39638** | **6.6** | **4.1-9.8** | **0** | **NA** | **PARTHENOLIDE** |
| **19** | **CCG-40283** | **11.9** | **5.2-20.0** | **0** | **NA** | **PRIMAQUINE PHOSPHATE** |
| **20** | **CCG-38545** | **10.2** | **6.8-14.7** | **0** | **NA** | **PYRITHIONE ZINC** |
| 21 | CCG-212462 | 6.6 | 3.8-10.2 | 0 | NA | SANGUINARIUM CHLORIDE |
| 22 | CCG-101015 | 16.5 | 5.6-37.3 | 0 | NA | PTEROSTILBENE |
| **23** | **CCG-214482** | **7.7** | **5.5-10.5** | **0** | **NA** | **METHYL GAMBOGATE** |

**MYH**

| Count | CCG_Number | $EC_{50}$($\mu$M) | 95% CI | Bottom | 95% CI | NAME_STRAIN_GENEID |
|---|---|---|---|---|---|---|
| *1* | *CCG-40298* | *inactive* | - | - | - | *AMINOTHIAZOLE* |
| **2** | **CCG-264834** | **indeterminant** | | | | **17-DMAG** |
| **3** | **CCG-208039** | **36.8** | **2.4-444** | **57.5** | **0-87.2** | **17-AAG** |
| **4** | **CCG-269784** | **inactive** | | | | **PU-H71** |
| 5 | CCG-264804 | 11.7 | 0-185.6 | 60.6 | 0-90.7 | NVP-AUY922 |
| **6** | **CCG-273380** | **inactive** | | | | **JG98** |
| **7** | **CCG-39082** | **inactive** | | | | **HYDROQUINONE** |
| **8** | **CCG-39719** | **67.4** | **4.7-290.6** | **0** | **NA** | **CETYLPYRIDINIUM CHLORIDE** |
| *9* | *CCG-39846* | *13.7* | *3.5-56.2* | *37.6* | *0-60.5* | *GITOXIGENIN* |
| 10 | CCG-39124 | 24.4 | 10.4-50.2 | 0 | NA | NONOXYNOL-9 |
| *11* | *CCG-38636* | *indeterminant* | - | - | - | *GITOXIGENIN DIACETATE* |

| | | | | | | |
|---|---|---|---|---|---|---|
| 12 | CCG-38646 | indeterminate | - | - | - | STROPHANTHIDINIC ACID LACTONE ACETATE |
| **13** | **CCG-40051** | **inactive** | | | | **DIHYDROCELASTRYL DIACETATE** |
| 14 | CCG-230895 | indeterminate | | | | OUABAIN OCTAHYDRATE |
| **15** | **CCG-39763** | **9.3** | **1.8-37.8** | **30.4** | **0-51.8** | **GENTIAN VIOLET** |
| **16** | **CCG-39917** | **71.5** | **5.5-233** | **0** | **NA** | **DIHYDROCELASTROL** |
| **17** | **CCG-39919** | **68.6** | **8.1-186.7** | **0** | **NA** | **CELASTROL** |
| **18** | **CCG-39638** | **inactive** | | | | **PARTHENOLIDE** |
| **19** | **CCG-40283** | **indeterminant** | | | | **PRIMAQUINE PHOSPHATE** |
| **20** | **CCG-38545** | **inactive** | | | | **PYRITHIONE ZINC** |
| 21 | CCG-212462 | 51.9 | 2.9-154.2 | 0 | NA | SANGUINARIUM CHLORIDE |
| 22 | CCG-101015 | 61.6 | 15.3-168.1 | 0 | NA | PTEROSTILBENE |
| **23** | **CCG-214482** | **68.9** | **12.6-240** | **0** | **NA** | **METHYL GAMBOGATE** |

**Compound Conclusion: Active = Bold,** *Inactive = italic*, Indeterminant = grey
Analysis: non-linear fit using inhibitor vs. response model assuming bottom > 0, top = 100, hill-slope -1 was performed using GraphPad Prism.

**Supplemental Table 8: Cell Toxicity Results**

| Count | CCG_Number | $EC_{50}$ (μM) | 95% CI | NAME_STRAIN_GENEID |
|---|---|---|---|---|
| **1** | **CCG-264834** | **>60** | **-** | **17-DMAG** |
| **2** | **CCG-208039** | **>60** | **-** | **17-AAG** |
| **3** | **CCG-269784** | **>60** | **-** | **PU-H71** |
| **4** | **CCG-273380** | **>60** | **-** | **JG98** |
| *5* | *CCG-39082* | *52.1* | *21.6-148* | HYDROQUINONE |
| *6* | *CCG-39719* | *32.1* | *19.2-55.1* | CETYLPYRIDINIUM CHLORIDE |
| *7* | *CCG-40051* | *33.2* | *26.0-42.7* | DIHYDROCELASTRYL DIACETATE |
| *8* | *CCG-39763* | *39.3* | *17.8-91.9* | GENTIAN VIOLET |
| *9* | *CCG-39917* | *18.0* | *10.3-31.3* | DIHYDROCELASTROL |
| *10* | *CCG-39919* | *25.9* | *20.1-33.4* | CELASTROL |
| **11** | **CCG-39638** | **60.8** | **35.9-108.7** | **PARTHENOLIDE** |
| **12** | **CCG-40283** | **>60** | **-** | **PRIMAQUINE PHOSPHATE** |
| *13* | *CCG-38545* | *17.6* | *14.9-20.7* | PYRITHIONE ZINC |
| *14* | *CCG-214482* | *15.7* | *11.4-21.8* | METHYL GAMBOGATE |

Bold = non-toxic ($EC_{50}$>60μM). Non-linear fit using inhibitor vs response curves assuming bottom = 0% viability, top = 100% viability, hill-slope -1, were performed using GraphPad Prism.

**Supplemental Table 9. Structure Activity Relationships (SAR) for JG98 and PTL**

| CCG#-Compound Name | MyBP-C | | | | MYH | | | | |
|---|---|---|---|---|---|---|---|---|---|
| | EC50 µM | 95% CI | Top % Veh | 95% CI | EC50 µM | 95% CI | Top % Veh | 95% CI | N |
| **JG98** | **22.6** | **18.2-28.4** | **138** | **124-152** | **48.9** | **37.7-64.4** | **115** | **104-127** | **5** |
| 208765-MKT-077 | >50 | - | 76 | 65-87 | >50 | - | 156 | 130-184 | 2 |
| 208766 -YM-1 | >50 | - | 110 | 99-120 | >50 | - | 142 | 127-158 | 2 |
| 208767 -YM-8 | >50 | | 78 | 74-82 | >50 | - | 117 | - | 2 |
| JG-258 | >50 | - | 105 | 93-119 | >50 | - | 113 | 103-122 | 2 |
| JG-231 | 15.9 | 9.6-28.4 | 132 | 102-169 | 31.5 | 25.2-39.9 | 121 | 110-132 | 5 |
| JG-345 | 5.8 | 3.7-8.9 | 144 | 107-204 | 21.8 | 17.8-27.0 | 124 | 113-136 | 5 |
| **CCG #-Compound Name** | **MyBP-C** | | | | **MYH** | | | | |
| | **EC50 µM** | **95% CI** | **Top % Veh** | **95% CI** | **EC50 µM** | **95% CI** | **Top % Veh** | **95% CI** | **N** |
| **PTL** | **20.3** | **10.8-36.5** | **126** | **100-161** | **>50** | **-** | **97** | **-** | **4** |
| 36373-PTL* | 13.7 | 10.8-17.6 | 98 | 87-111 | >50 | - | 131 | 115-149 | 2 |
| 205026-PTL* | 17.6 | 12.3-26.0 | 88 | 73-105 | >50 | - | 119 | - | 2 |
| 208244-PTL* | 13.9 | 10.1-19.3 | 93 | 79-109 | >50 | - | 114 | 89-141 | 2 |
| 266943-PTL* | 52.3 | 31.7-93.2 | 83 | 65-101 | >50 | - | 123 | 95-152 | 2 |
| 36184-Pyrethrosin* | 52.4 | 30.2-106 | 100 | 75-127 | >50 | - | 126 | - | 2 |
| 35566-Eunicin | 15.9 | 10.2-25.5 | 66 | 52-82 | 21.3 | 11.0-48.7 | 113 | 77-154 | 2 |
| 35235-NSC36437 | >50 | - | 101 | - | >50 | - | 114 | - | 2 |
| 36035-Eupatundin* | 54.1 | 34.3-92.9 | 85 | 69-101 | >50 | - | 117 | 81-1154 | 2 |
| 266942-Micheliolide | >50 | - | 91 | 81-102 | >50 | - | 143 | - | 2 |
| 266810-Costuolide | >50 | - | 104 | 85-123 | >50 | - | 117 | - | 2 |
| 36376-Bohlmann | >50 | - | 93 | 79-108 | >50 | - | 118 | 91-146 | 2 |
| 36468-Alantolactone | >50 | - | 89 | 79-98 | >50 | - | 121 | 107-136 | 2 |

Legend notes: Inhibitor vs. response-variable slope = bottom equals 0, hill slope -2. * indicates compounds with epoxide moiety.

**Supplemental Table 10. Antibodies tested in the development of MyBP-C and MYH Alpha-LISA assays**

| **MyBP-C Alpha-LISA antibody screen** | |
|---|---|
| **Donor Bead Antibody** | **Acceptor Bead Antibody** |
| C0, rabbit, polyclonal, Samantha Harris | C5-C7, rabbit, polyclonal, Samantha Harris |
| C0, rabbit, polyclonal, Samantha Harris | C0, mouse, monoclonal Santa Cruz, sc-137180 |
| C5-C7, rabbit, polyclonal, Samantha Harris | C0, rabbit, Samantha Harris |
| C5-C7, rabbit, polyclonal, Samantha Harris | C0, mouse, monoclonal Santa Cruz, sc-137180 |
| C0, mouse, monoclonal Santa Cruz, sc-137180 | C0, rabbit, polyclonal, Samantha Harris |
| C0, mouse, monoclonal Santa Cruz, sc-137180 | C5-C7, rabbit, polyclonal, Samantha Harris |
| **MYH Alpha-LISA antibody screen** | |
| **Donor Bead Antibody** | **Acceptor Bead Antibody** |
| MYH7 mouse, monoclonal sc53090 | MYH7 Nterm, rabbit abgent |
| MYH7 Nterm, rabbit abgent | MYH7 mouse, monoclonal sc53090 |
| MYH7 mouse, monoclonal sc53090 | MYH7, rabbit, polyclonal ab228353 |
| MYH7, rabbit, polyclonal ab228353 | MYH7 mouse, monoclonal sc53090 |
| MYH7 mouse, monoclonal sc53090 | Myosin-light chain 2, rabbit monoclonal ab92721 |
| Myosin-light chain 2, rabbit monoclonal ab92721 | MYH7 mouse, monoclonal sc53090 |
| MYH6, mouse, monoclonal ab50967 | MYH7 Nterm, rabbit abgent |
| MYH7 Nterm, rabbit abgent | MYH6, mouse, monoclonal ab50967 |
| MYH6, mouse, monoclonal ab50967 | MYH7, rabbit, polyclonal ab228353 |
| MYH7, rabbit, polyclonal ab228353 | MYH6, mouse, monoclonal ab50967 |
| MYH6, mouse, monoclonal ab50967 | Myosin-light chain 2, rabbit monoclonal ab92721 |
| Myosin-light chain 2, rabbit monoclonal ab92721 | MYH6, mouse, monoclonal ab50967 |
| MYH7 Nterm, rabbit abgent | MYH7 Nterm, rabbit abgent |
| MYH7, rabbit, polyclonal ab228353 | MYH7, rabbit, polyclonal ab228353 |
